# The macroevolutionary impact of an innovation reversal in ray-finned fishes

**DOI:** 10.64898/2026.08.25.747124

**Authors:** Chase D. Brownstein, Richard C. Harrington, Julia E. Wood, Ava Ghezelayagh, Laura R.V. Alencar, Martha M. Muñoz, Christine E. Thacker, Thomas J. Near

## Abstract

The evolution of new traits can drive species diversification by facilitating the use of new resources,^1,2^ but environmental change may turn these same adaptations into liabilities.^3–5^ Trait loss is also often associated with the origin of new ecologies, but how losses modulate diversification remains unclear.^1,2,6–10^ The swim bladder allows ray-finned fishes to regulate their buoyancy and exploit ecosystems throughout the water column, yet this organ has been lost many times among species-rich lineages. Here, we show that timing and ecological context control the macroevolutionary effects of swim bladder loss. Many lineages of fishes lost the swim bladder over the last 66 million years as they specialized for benthic habitats where buoyancy regulation is unnecessary. Swim bladder loss enabled the descendants of these benthic fishes to diversify in the deep sea where extreme pressure makes its inflation untenable,^11^ and in the frigid, oxygen-saturated Southern Ocean, where loss of the oxygen delivery mechanisms required for swim bladder inflation carries little physiological cost.^12,13^ Yet, we detect a selective filter associated with swim bladder loss during extreme global warming 56 to 50 million years ago, when its absence limited the capacity of fishes to escape ecological disruptions on the ocean floor. These contrasting patterns explain how the loss of a complex trait promoted major ecological transitions without increasing overall diversification through deep time. As human activity drives rapid global warming, the evolutionary legacies of swim bladder loss may again shape the fate of marine fish diversity.

## Introduction

The evolution of traits exerts a profound influence on the origination and extinction of clades in deep time.^1,14–17^ Across many of the most widely studied evolutionary radiations, the origins of novel features that enable the use of new resources and the invasion of new habitats, called key innovations, appear to have driven rapid species diversification.^2,15,18–21^ However, the functional and ecological context in which novelties evolve^5,22–25^ and the ease by which they are gained or lost^4,8,26,27^ are equally important for understanding how they contribute to species richness. Ecological change can render these same innovations functionally unnecessary or even costly to survival.^5,28,29^

The loss of complex functional innovations is another classic pattern in many species that have invaded new environments. For example, cave-adapted animals often lose functional eyes and pigmented bodies as they evolve in darkness,^30,31^ and birds commonly lose the ability to fly after colonizing islands.^32–34^ Nonetheless, the impact of trait loss on diversification has remained intensely debated for over two centuries. Does the loss of functionally important traits tend to instigate an evolutionary trajectory towards extinction, as Charles Darwin hypothesized,^6^ or can losing a functional trait facilitate diversification, as suggested by a growing number of biological systems?^35,36^

Most of the more than 35,720 living species of ray-finned fishes possess a swim bladder, an organ that regulates buoyancy through its inflation and deflation.^37,38^ In some fishes, the swim bladder attaches to the esophagus and can be used for respiration and buoyancy control ^37,38^ Fishes that lack this connection inflate the swim bladder solely using oxygen in the bloodstream through a web of vasculature called the *rete mirabile*.^38–40^ The rapid inflation of the swim bladder in these fishes is achieved by a physiological phenomenon called the Root effect, in which oxygen is rapidly released from the bloodstream after slight decreases in pH.^37,41–43^ Because it allows fishes to move throughout the water column and access a variety of resources in different habitats, the swim bladder is hypothesized to be a key innovation driving the accumulation of ray-finned fish diversity.^39,40^ Yet, many rapidly-diversifying lineages of fishes,^44^ including deep-sea anglerfishes,^45,46^ flatfishes,^47,48^ darters,^49–51^ snailfishes,^52^ and the adaptive radiation of Antarctic notothenioids,^53–55^ have secondarily lost the swim bladder in adults. These lineages occupy benthic habitats, where buoyancy control is thought to be functionally unnecessary, or environments where this organ is costly, such as the extreme pressure conditions of the deep sea.^8,11,56–58^ Despite decades of interest in the functional implications of swim bladder loss and its association with transitions between habitats,^11,37,39,52,58,59^ this association has not been tested at broad evolutionary scales, and it is unclear how the loss of this trait might have shaped the diversity of fishes through deep time. Here, we examine the contrasting effects of swim bladder loss on ecological transition and species diversification throughout 90 million years of global environmental perturbation. We show that the loss of this complex trait and its associated physiochemical systems has promoted invasions of new habitats and ecologies, providing the substrate for species diversification. Yet, the loss of the swim bladder has also functioned to increase the exposure of fishes to ecological perturbations when global climate change induced major extinctions in habitats associated with its loss. As anthropogenic climate change rapidly warms the planet, these macroevolutionary patterns may be recapitulated.

## Results

### Tempo and mode of swim bladder loss in deep time

Swim bladders have been lost more than 30 times in ray-finned fishes, and a similarly long list of mutations is responsible.^52^ The invariant loss of swim bladders across all species in several taxonomic orders of ray-finned fishes, as well as the observation that this organ degenerates in different ways within^52^ and among^60^ lineages in a manner dependent on mutations involved,^52^ suggests that regains of this complex trait are unlikely after its loss at deep time scales. Losses of the swim bladder are disproportionately concentrated in *Eupercaria*, a clade of more than 7,800 species^61,66,67^ of fishes, including many evolutionary radiations posited to have been driven by the gain of functional, physiological, and life history innovations.^8,55,68–74^ The common ancestor of *Eupercaria* lacked a connection between the swim bladder and esophagus, and species within *Eupercaria* must inflate the swim bladder using oxygen delivered from the bloodstream.^39,43,52^ The small number of swim bladder losses that occur in lineages outside *Eupercaria* are phylogenetically isolated across hundreds of million years of Earth history.^52,62,75^ As such, we focused on constructing a species-rich phylogeny of *Eupercaria* to investigate the evolutionary impacts of swim bladder loss. The time-calibrated phylogeny that we present (Figure 1a; Extended Data Figures 1–2) is based on 1,314 ultraconserved element loci, includes 95% of taxonomic families, 40% of genera, and 19% (n=1,301) of species in *Eupercaria*, and is the most complete phylogenetic hypothesis built for this clade based on genome-wide marker data. The phylogeny of *Eupercaria* that we present is congruent with previously trees inferred using genomic data.^61,62,64,71^ The divergence times that we estimate for major lineages in *Eupercaria* (Figure 1a; Extended Data Figures 1–2; Supplementary Information) are also comparable to previous estimates. ^61,62,64,76^

**Figure 1.**
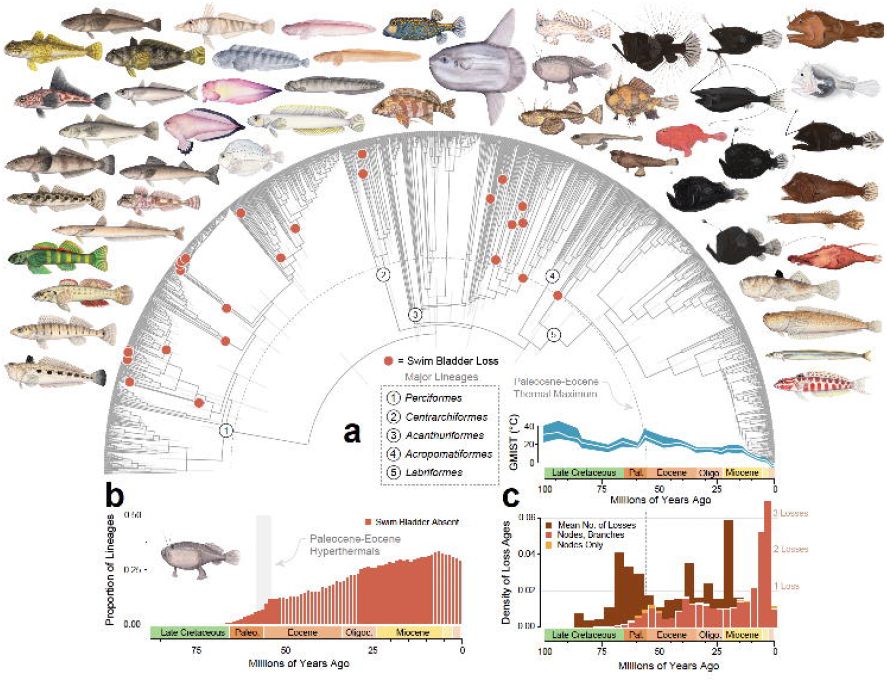
Evolutionary Tempo and Mode of Swim Bladder Loss Across *Eupercaria*. (a) Time calibrated phylogeny of 1,332 tips fixed to the maximum likelihood single partition topology for the 75% complete matrix estimated using Bayesian node-dating with 48 fossils. Outgroups are removed. Bars at nodes indicate 95% highest posterior density intervals of divergence times. Red dots denote independent losses of the swim bladder along the phylogeny. The turquoise line to the right is the global mean sea temperature curve from Judd et al.,^100^ with 95% confidence intervals shaded. (b) Proportion of lineages in each state through time in million-year-intervals. Shown in the foreground of (c) is the density plot of ages estimated across 10,000 randomly sampled posterior trees of crown and stem lineages where independent swim bladder loss are inferred to take place in our ancestral state reconstruction in (a). The density plot shows the density of ages when only losses at nodes (dark orange) are recorded, or when losses along branches (i.e., along stem lineages; light oranges) are additionally considered. In the background is a curve showing the mean number of swim bladder losses reconstructed across 1,000 simulated stochastic mappings along the consensus time-calibrated phylogeny. Illustrations are by Julie Johnson (lifesciencestudios.com).

Ancestral state reconstruction of swim bladder evolution using this time-calibrated phylogeny and a new dataset of swim bladder presence for 1271 species suggests that this organ was lost a minimum of 25 times in *Eupercaria* (Figure 1a). These losses, which represent more than half of known instances across ray-finned fishes, began to accumulate after the Cretaceous-Paleogene Mass Extinction 66.02 million years ago (Figure 1b) in the marine realm (Extended Data Figure 3a). There is a lag time of 21.74 million years between the origin of *Eupercaria* in the Coniacian Stage of the Late Cretaceous, 87.74 Ma (95% highest posterior density [HPD] intervals: 79.53, 98.12 Ma), and the loss of swim bladders among living lineages, which is a fourth of the total evolutionary history of *Eupercaria.* Several independent losses of the swim bladder occur in darters, perches, and walleyes (*Percidae*), which is the most species-rich freshwater lineage of *Eupercaria*.^77^ We also reconstruct losses of the swim bladder in the common ancestors of the three most species-rich lineages of fishes in the polar oceans,^73,74^ eelpouts and wolffishes (*Zoarcoidea*), sculpins, snailfishes, and relatives (*Cottoidea*), and the Antarctic notothenioid radiation (*Notothenioidei*), as well as in these and other species-rich deep-sea fish clades, including pelagic anglerfishes (*Ceratioidea*) and coffinfishes (*Chaunacidae*), monkfishes (*Lophiidae*), batfishes (*Ogcocephalidae*), and some lineages of rockfishes (*Sebastidae*) (Figure 1a). Notably, snailfishes (*Liparidae*) include the deepest-dwelling vertebrate species.^72,80,81^

We examined the timescale of swim bladder loss across *Eupercaria* using methods that account for uncertainty in the estimation of both lineage ages and loss times. We generated density curves of ages of nodes and branches reconstructed to have lost the swim bladder (Figure 1a) taken across 10,000 time trees from the posterior sample in our Bayesian time-calibration analysis. These density curves alternately account for the loss of the swim bladder along the branch leading to a node or singleton tip where this organ is absent in our summary ancestral state reconstruction (Figure 1a). When plotted, these curves demonstrate an abrupt downturn in the number of independent loss times at the Paleocene-Eocene boundary 56 million years ago, following the steady accumulation of losses in the first ten million years after the Cretaceous-Paleogene Mass Extinction (Figure 1c). Independent losses of the swim bladder do not rebound until 40 Ma, well into the Eocene (Figure 1c). Similarly, the mean number of losses across the 1,000 stochastic mappings used to generate our summary ancestral state reconstruction (Figure 1c) shows a downturn in the average number of losses during the Paleocene, and a low point in the Eocene, following a peak in new losses at the terminal Cretaceous (Figure 1c); this corresponds to a stagnation in the proportion of lineages without a swim bladder during the Paleocene-Eocene transition (Figure 1b). These results illustrate a period of suppressed swim bladder losses during the late Paleocene and early Eocene.

### The Exaptive Nature of Swim Bladder Loss

We next examined how swim bladder loss is associated with major transitions across habitats and ecologies. We first observe that swim bladder losses are concentrated in species tied to benthic habitats (Figure 2a; Extended Data Figure 3). Across our sample, 78% of species lacking a swim bladder are benthic (i.e., remaining in contact with the bottom for the majority of their life),^82^ compared to only 7% of species with a swim bladder. Our ancestral state reconstruction of water column occupation demonstrates that, across the 25 independent losses of the swim bladder, none precede transitions into benthic habitats, 16 occur in lineages that had already become benthic, and eight occur at the same nodes as transitions to benthic habitats (Figure 2a; Extended Data Figure 3a). Except for oceanic sunfishes, all pelagic lineages that lack a swim bladder evolved from benthic ancestors (Extended Data Figure 3a). A likelihood ratio test of models where swim bladder presence and benthic habitat occupation are alternatively treated as evolving in a correlated manner and swim bladder loss is irreversible supports the hypothesis that swim bladder loss and benthic habitat occupancy are correlated (LR = 117.5066, df = 3, p = 2.657031 x 10^-25^). The swim bladder loss rate in benthic lineages is 43.68 times greater than the rate in demersal and pelagic lineages (Figure 2a), and the small proportion of benthic lineages that leave benthic habitats and lose the swim bladder all transition to demersal ecosystems (Figure 2a). In contrast, transitions to fully pelagic ecologies are only nine times faster in fishes with swim bladders than in fishes without them.

**Figure 2.**
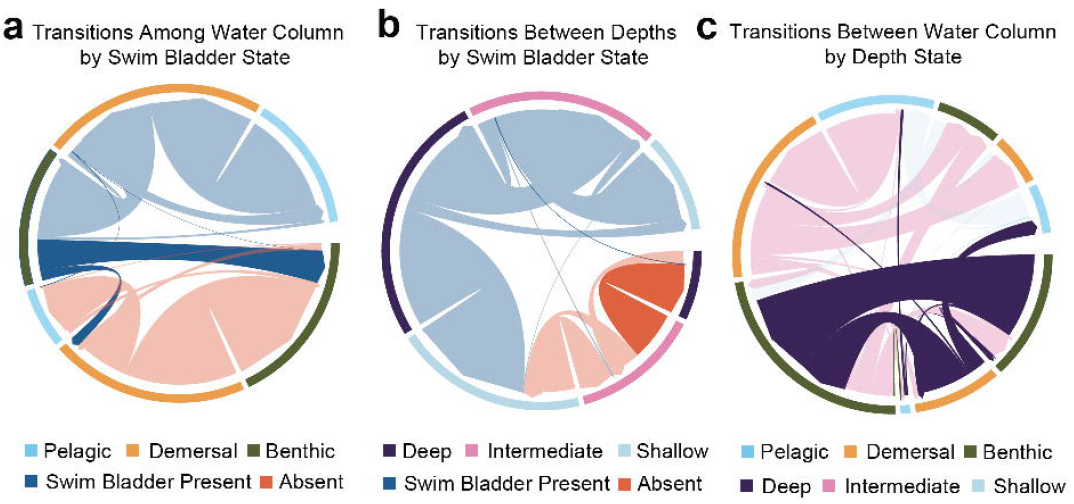
Major Transitions and the Exaptive Nature of Swim Bladder Loss. (a) Chord diagrams showing transitions between ecological and depth states by swim bladder state (left, center) and depth (right). For the left and center chord diagrams, blue chords denote transitions by lineages where the swim bladder is ancestrally present, and orange chords denote transitions by lineages where the swim bladder is absent. For the right chord diagram, dark blue chords denote transitions by lineages at 1000 + m depth, light blue chords denote transitions by lineages at 0 to 1999 m depth, and orange chords denote transitions by lineages at 200 to 999 m depth. The rate of loss in benthic-dwelling fish lineages is over 43 times larger than the rate of loss in non-benthic fish lineages. In the left plot, transitions from swim bladder presence to absence are highlighted. In the center plot, transitions to deep water without the swim bladder are highlighted. In the right plot, transitions by deep-sea lineages are highlighted. (b) Distributions of body aspect ratio are shown for species both possessing and lacking a swim bladder. There is a significant association between the absence of the swim bladder and higher aspect ratio according to phylogenetic logistic regression. (c) Distributions of body aspect ratio are shown for benthic and non-benthic species. There is a significant association between benthic habitat occupancy and higher aspect ratio according to phylogenetic logistic regression.

Loss of the swim bladder is also associated with the characteristic specialization for benthic habitats in ray-finned fishes: flattened bodies.^82^ Phylogenetic logistic regression reveals a significant (p = 1.755 x 10^-7^, regression coefficient = -1.69499) association between swim bladder absence and increased flatness, quantified by the ratio of maximum body depth to body width (Figure 2b). We also find a significant relationship between increased body flatness and benthic ecology (p = 7.38 x 10^-4^, regression coefficient = -2.93411), as well as flatness and water depth (p = 2.897 1.755 x 10^-3^, regression coefficient 0.84778). Flattened body shapes in our sample have a wide phylogenetic distribution across fishes in our phylogeny that lack a swim bladder, although species of batfishes and monkfishes show particularly extreme values (Extended Data Figure 4a). Phylogenetic principal components analysis reveals that many species that lack a swim bladder are concentrated in a region of morphospace that corresponds to shallow, elongated body plans (Extended Data Figure 4b; Table S2). These observations are consistent with the hypothesis that swim bladder loss is associated with specialization for benthic habitats where buoyancy regulation might pose an unnecessary energetic demand.

We also examined associations between swim bladder evolution and transitions to the deep sea, where its loss in many lineages of deepwater fishes has been considered an evolutionary response to the energetic demands of its inflation with oxygen under the extreme pressures that characterize life at more than 1,000 meters below sea level.^11^ We find a significant correlation between swim bladder loss and transitions to the deep ocean (LR = 66.8854, df = 3, p = 1.981436 x 10^-14^). All transitions to deep water in fish lineages that have lost the swim bladder occur after the invasion of benthic habitats in these lineages (Figure 2a; Extended Data Figure 1a), and consist of both deep-water benthic clades with flattened bodies, such as batfishes and eelpouts, or secondarily pelagic fishes, such as ceratioid anglerfishes and several lineages in *Notothenioidei*, that achieve buoyancy by means of lipid-rich tissues.^59,83,84^ Across *Eupercaria*, the vast majority of transitions to and from depths exceeding 1000 meters occur among benthic fishes (Figure 2a). This suggests that the loss of the swim bladder in benthic fishes has served as an exaptation that enabled repeated invasions of the deep sea, driving the accumulation of deep marine lineage diversity.

### Swim bladder loss indirectly imparts a signature of diversification

We tested the impact of swim bladder losses on lineage diversification in *Eupercaria* by estimating swim bladder-associated, benthic habitat-associated, and depth-associated diversification rates under Binary (BiSSE) and (II) Hidden State Speciation and Extinction (HiSSE) models, which analyze differences in trait-associated diversification rates with and without hidden states influencing diversification. We tested these on both the complete phylogeny and a pruned version of our time-calibrated tree in which genera were represented by single species in order to assess whether recent divergences among species-rich genera might drive or suppress signals of diversification. BiSSE analyses favor a non-significant net increase in diversification rate associated with swim bladder loss and benthic habitat occupancy, but a significant increase associated with invasions of the deep sea (Figure 3a; Extended Data Figures 4–5). These results are consistent when our dataset is pruned to include only one representative species per genus, except for BiSSE analysis of depth-driven diversification, in which rates associated with deep and shallow-intermediate depths largely overlap (Extended Data Figure 4). Across all these traits and both analyses of the complete and subsampled phylogeny, HiSSE model comparisons favor a character-dependent hidden state model (Figure 2a; Table S4), suggesting these traits only partially explain diversification regimes across *Eupercaria* (Figure 2a; Extended Data Figures 4–5).

**Figure 3.**
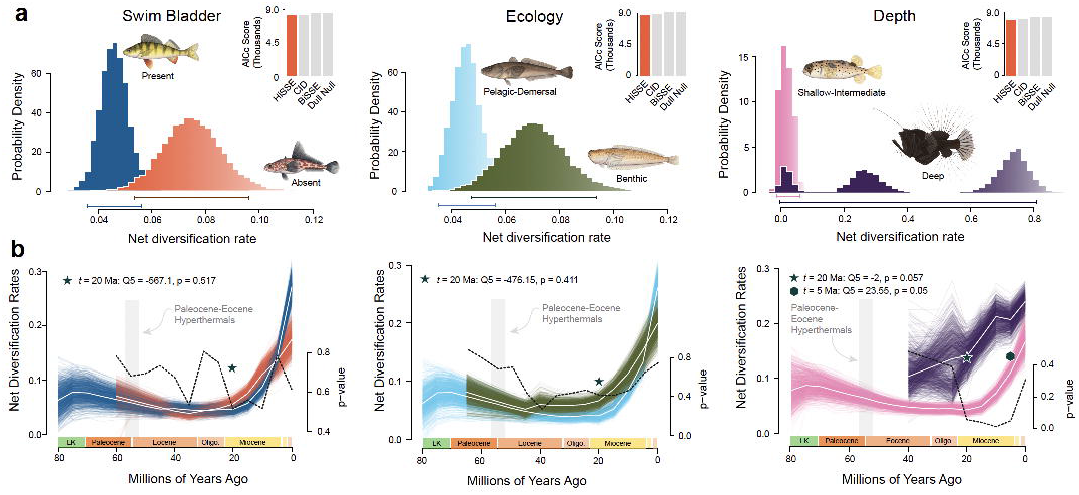
Context-Specific Signals of Diversification Associated With Swim Bladder Loss. (a) Trait-associated diversification rate distributions estimated under BiSSE using an irreversible trait model and HiSSE model fitting. In the distribution plots, bars below distributions indicate 95% highest posterior density intervals. The inset bars show AICc scores for each of four models tested in the HiSSE framework, with the best-fit in red. (b) Trait-associated diversification rate curves estimated using *deepSTRAPP*. Mean rate curves are highlighted among the estimated posterior rate curves. The dotted line denotes the p-values from STRAPP tests for differences in trait-associated diversification rates across 5 million year intervals. The scales for the p values are displayed to the right of each plot. All analyses shown in this figure were conducted using the full phylogeny. Illustrations are by Julie Johnson (lifesciencestudios.com).

To test whether swim bladder loss and ecological transitions promote diversification in a manner that varies across timescales, we estimated variation in trait-associated diversification rates and variation in these rates through time using STRAPP tests. ^85^ These analyses show net diversification of benthic lineages and lineages that lack a swim bladder decreases around the Cretaceous-Paleocene transition before rebound starting ∼20 Ma (Figure 3b). Net diversification rates do not significantly differ at any time step between lineages possessing or lacking the swim bladder or lineages that occupy benthic or non-benthic habitats (Figure 3b; Extended Data Figures 6–7). However, we do detect a significant (p < 0.05) difference between rates of diversification associated with occupation of deep and shallow-intermediate habitats between 20 and 5 million years ago (Figure 3b; Extended Data Figure 6). Diversification rates associated with deep habitats also remain, on average, elevated for the entire history of deep sea colonization by lineages of *Eupercaria* (Figure 3b). This elevated diversification rate is driven by recent speciation within genera of deep-sea fishes in *Perciformes* and *Lophioidei*. When depth-associated diversification rates through time are estimated on a pruned version of the phylogeny in which genera are only represented by one species, this significant increase in deep-sea diversification starting 20 million years ago disappears (Extended Data Figure 8). These analyses demonstrate that losing the swim bladder has not increased lineage diversification through time, contrasting with expectations in a scenario where losing this organ has acted as a key innovation. However, habitats associated with swim bladder loss do differ in their diversification impacts: transitions to deep water habitats are significantly associated with an increased rate of lineage diversification over the last 20 million years (Figure 3b).

### Swim Bladder Loss and Extinction Risk in Extreme Habitats

Our time-calibrated phylogeny suggests that fishes lacking swim bladders invaded and diversified in physiochemically extreme ecosystems in the deep sea^8,83,84,86^ and polar oceans,^44,61,87^ following the Eocene. Among these, snailfishes, eelpouts, and Antarctic notothenioid adaptive radiation are the most species-rich lineages in the Arctic and Southern oceans^78,88^ and drive the inverse latitudinal speciation rate gradient observed in ray-finned fishes.^44^ The 40 million years of near-continuous global cooling since the Eocene (Figure 2) is associated with the degradation of the physiochemical basis of swim bladder function in some of these polar marine radiations, which evolved antifreeze proteins that enable their survival in freezing water after losing the swim bladder.^53,55,72,89^ The Root effect, which describes the rapid release of oxygen by ray-finned fish hemoglobins during slight increases in blood acidity, allows fishes in *Eupercaria* to inflate the swim bladder via the *rete mirabile* in the absence of a connection between the bladder and esophagus.^13,37,39,42^ However, many species of Antarctic notothenioids and eelpouts do not exhibit the Root effect.^13,90,91^ Our ancestral state reconstruction of Root effect magnitude shows that multiple lineages in each of these polar fish radiations evolved reduced Root effects soon after evolving specialized antifreeze proteins (Figure 4; Extended Data Figure 9a). Lag times of 10 to 20 million years exist between the loss of the swim bladder in *Notothenioidei* and *Zoarcoidea* and the loss of the Root effect in lineages nested within these radiations (Figure 1a; Figure 4). Hemoglobin, the oxygen delivery protein in blood and the molecule that experiences the Root effect, is lost in the notothenioid clade *Channichthyidae* (icefishes), and myoglobin, which binds oxygen in tissues, is also lost in several lineages of icefishes^92^ during their diversification in the thermally stable, cool, and oxygen-rich waters of the Southern Ocean.^91–95^ Other independent, complete losses of the Root effect occur in the Longfin Icedevil *Aethotaxis micropteryx* and the Antarctic Ploughfish *Gymnodraco acuticeps* (Figure 4).

**Figure 4.**
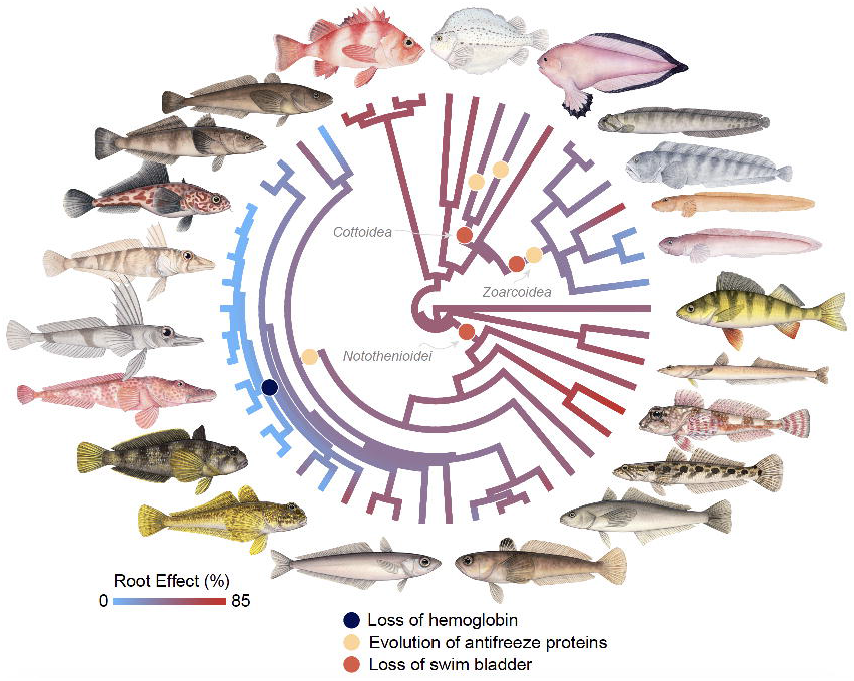
Ecological Legacies of Swim Bladder Loss. Ancestral state reconstruction for percent reduction in oxygen saturation from across one pH value decrease (see di Prisco et al. 2007) for species in *Perciformes* (n=49). Non-perciform species have been removed. Note the complete sampling of all lineages at the level of taxonomic families in *Notothenioidei*, and sampling of most taxonomic families in *Zoarcoidea*. Illustrations are by Julie Johnson (lifesciencestudios.com).

The observation that Root effect hemoglobins allow fish to enhance oxygen delivery to tissues^43,96^ provides an explanation for the lag time between swim bladder and Root effect loss. Root effect losses only occur within lineages of fishes inhabiting the Southern Ocean. Living in the oxygen-supersaturated waters of the Southern Ocean might allow these fishes to compensate for the loss of Root effect hemoglobins,^97^ as well as reducing metabolic costs associated with life at higher temperatures ^98^. This hypothesis is consistent with the pattern of Root effect disparity we observe in notothenioids, which falls under the 95% confidence intervals of disparity accumulation under Brownian motion and contrasts with the expected pattern if strong stabilizing selection was maintaining similar Root effects among notothenioid lineages (Extended Data Figure 9b). Thus, environmental context appears to have enabled the stepwise degeneration of the Root effect and other oxygen delivery systems in polar fishes.

## Discussion and Conclusions

Our analyses establish that the relationship between swim bladder loss, ecological transitions, and lineage diversification is contingent on temporal and environmental context. There are highly significant associations between swim bladder loss and ecological transitions among ecologies and depth levels.^1,2,99^ Despite this, loss of the swim bladder is not associated with any clear signature of increased species diversification. Instead, losing the swim bladder is associated with decreased survival during ecological upheaval in the deep past, especially during the Paleocene and Eocene 56 to 50 million years ago (Figure 1b-c, Figure 2), when global sea temperature spiked to over 35° Celsius in the most extreme fluctuations of the past 80 million years (Figure 1).^100^ Both the accumulation of lineages lacking a swim bladder and the inferred rate of new swim bladder losses decline at the Paleocene-Eocene Thermal Maximum (PETM) 56 Ma and stay low throughout the Eocene Climatic Optimum (ECO) from 55-50 Ma, which together form a prolonged period of extreme global warming (Figure 1a). The proportion of benthic lineages in *Eupercaria* continues to steadily increase throughout this interval, and the proportion of deep-water lineages remains constant (Extended Data Figure 2b-c). These contrasting patterns suggest a complex relationship between swim bladder evolution and diversification that is only partially mediated by ecology.

Our results explain how swim bladder loss simultaneously promotes ecological transition and extinction risk in the long term. Loss of the swim bladder removes an energetic cost for fishes that have become specialized for benthic habitats where buoyancy regulation is unnecessary and is significantly associated with transitions and subsequent specializations for these environments. The selective extinctions of fishes in *Eupercaria* that lost the swim bladder indicated by our analyses are also consistent with fossil evidence for the effects of extreme Eocene global warming on marine diversity. During the PETM and ECO, more than half of deep-dwelling benthic foraminifera went extinct as global ocean acidification impacted calcifying organisms and forced many lineages toward the poles.^101–104^ The major benthic turnover event that occurred at the PETM was felt by faunas across the ocean depth gradient,^105–109^ and would have impacted fishes without swim bladders that were ecologically and physically tied to the ocean floor.

Although swim bladders rarely fossilize, the hypothesis of a selective extinction of fishes lacking swim bladders during the PETM and ECO is supported by the fossil record of living lineages that lack this organ. †*Enchodontoidei*, a diverse, potentially paraphyletic lineage nested within living aulopiform fishes that was a major component of Cretaceous and Paleogene marine teleost faunas, went extinct in the earliest Eocene after losing a considerable proportion of their diversity at the K-Pg boundary.^110,111^ Living species of *Aulopiformes* all lack swim bladders,^52^ implying that this organ was lost in their common ancestor, and therefore absent in †*Enchodontoidei.* The extinction of †*Enchodontoidei* during the earliest Eocene is consistent with selection against swim bladder loss during Paleocene-Eocene warming. The major lineages of fishes outside *Aulopiformes* and *Eupercaria* that invariably lack a swim bladder, such as oarfishes, ribbonfishes, flatfishes, only appear in the fossil record after 50 million years ago.^47,112^ Time-calibrated phylogenies of these lineages indicate rapid radiations after the Cretaceous-Paleogene Mass Extinction and around the Paleocene-Eocene boundary ^48,112,113^. These observations suggest that nearly all major fish lineages that lost the swim bladder did so following global temperature spikes that mark the K-Pg and Paleocene-Eocene boundaries.

The evolutionary history of the swim bladder illuminates how ecological context and historical contingency shape the macroevolutionary impact of complex trait loss. Many ray-finned fishes, from the adaptive radiation of Antarctic notothenioids ^53,55,114^ to the temperate continental radiation of darters,^51,56,60^ lost the swim bladder after undergoing major transitions into benthic habitats. In the descendants of some of these benthic lineages, swim bladder loss enabled invasions of extreme ecosystems and the degeneration of related, energetically costly aspects of organismal biology. For example, ceratoid anglerfishes evolved from a benthic ancestor that lacked a swim bladder (Figure 2)^8^ and diversified in the extreme pressures of the deep open ocean, where a gas-inflated organ is a functional liability.^11,59^ The impact of the swim bladder is dependent on when this trait reversal occurs over geological timescales, as our results substantiate a selective bottleneck against species that lacked a swim bladder during Eocene warming events that induced major extinctions of benthic marine faunas. This selective filter is consistent with the classic scenario in which specialization increases extinction risk during environmental perturbation.^28,115^ In these lineages of fishes, the loss of a complex functional trait might have been energetically advantageous in the short term, but critically maladaptive in the long term.

Today, as rapid anthropogenic global warming approaches the severity of Eocene hyperthermals,^116^ our study emphasizes that the very feature which may have helped facilitate radiations of fishes in extreme environments, such as the Earth’s polar oceans, makes them particularly susceptible to extinction. Conceivably, warming of these polar waters will reverse the conditions that allow fishes to tolerate loss of oxygen delivery systems involved in swim bladder inflation. This compound cost of losing the swim bladder, the physiochemical adaptations that support its function, and the metabolic benefits of life in these extreme environments^98^ is an observable consequence of macroevolutionary processes with dire implications for the survival of species today. In an echo of the deep past, humanity may once more shift the fortunes of species with this innovation reversal.

## Methods

### Taxonomy

In this manuscript, we follow the phylogenetic rank-free taxonomy of ray-finned fishes proposed recently in several monographic treatments.^117,118^ In accordance with the conventions of the *PhyloCode*^119^ and emerging trends in the literature,^120^ we italicize all formal clade names.

### Sequence Dataset Assembly

We assembled a phylogenomic dataset of 1,750 individuals representing 1,302 eupercarian species and 33 outgroups by sequencing ultraconserved element (UCE) loci, a type of genome-wide marker, based on a probe set of 1,314 individual loci developed for acanthomorph teleosts ^64^. Our sequence dataset contains both previously published sequences used to infer the phylogeny of acanthomorphs ^61,64^, anglerfishes ^8^, angelfishes ^121^, eelpouts and relatives ^70^, and wrasses and parrotfishes ^71^, *Percidae*,8/25/2026 3:05:00 PM^122^ UCE loci skimmed from all genomes of eupercarians available on the NCBI GenBank sequence repository as of summer 2025, and newly sequenced individuals. Our phylogeny is the first phylogenomic analysis to sample several deeply divergent and species-rich lineages in *Perciformes*, *Acanthuriformes*, and *Acropomatiformes*, including *Hypoplectrodes* basslets, *Epinephelus* groupers, *Synagrops* splitfins, Grape-eyed Seabass *Hemilutjanus microphthalmos*, and Long-Finned Pike *Dinolestes lewini*.

### Sequence Extraction and Library Preparation

For newly sequenced individuals, we extracted DNA using Qiagen DNEasy Blood and Tissue Kits, quantified the DNA concentration of each extraction using a Qubit fluorometer (Life Technologies), assessed the DNA fragment size and integrity using gel electrophoresis, and sheared approximately 500 ng of DNA per sample using a QSonic Q800R3 sonicator, which produced fragments of 300 to 600 bp in length. Next, we used Kapa HyperPrep kits (Kapa Biosystems) and Illumina TruSeq iTru5 and iTru7 adapters ^123^ to prepare genomic libraries, which were then hybrid-enriched for the acanthomorph UCE probe set ^64^ using probes obtained from Daicel Arbor Biosciences following previously published methods ^61^. We sequenced libraries using 150 bp paired-end sequencing on Illumina NovaSeq platforms after creating an equimolar pool of enriched libraries.

### Read Processing and Sequence Alignment

Once we received raw reads from our new DNA libraries, we used *phyluce* 1.7.3 to process and assemble reads *de novo* ^124^. We trimmed reads using the *Trimmomatic* ^125–127^ wrapper *Illumiprocessor* v. 2.0 ^128^ to remove adapter index sequences and low quality bases, and assembled raw reads using *SPADES* v. 3.14.1 ^129^ with default settings. Next, we extracted UCE sequences from reads using the acanthomorph UCE probe set, which contains 1,314 target loci ^64^. After acquiring UCE contigs from newly sequenced specimens, we combined them with previously sequenced UCE loci and UCE loci that we extracted from available genome assemblies using custom scripts in *phyluce*. To assess for completeness of our dataset, we produced an occupancy table (Figure S22). This table was visualized via *Claude* Opus 4.7. We aligned the resulting sequence set using *MAFFT* ^124^ called from *phyluce* and then produced 75% and 90% complete taxon-sequence matrices. Next, we used *CIAlign* ^130^ to visualize and check our dataset for chimeric alignments. After this step, we retained a total of 995 UCE loci sampled for 1,750 individuals in the 75% complete matrix, and 877 UCE loci sampled for the same count in the 90% complete matrix. We ran all sequence processing and alignment steps on the Yale High Performance Computing Cluster McCleary.

### Phylogenetic Analyses and Concordance Factors

We conducted maximum likelihood phylogenetic analyses of our 75% and 90% UCE matrices using *IQ-TREE* 2 ^131^ on the Yale High Performance Computing Cluster McCleary. We ran analyses treating the dataset as a single set of concatenated sequences and also inferred gene trees to calculate gene and site concordance factors. For all analyses, we used ModelFinder Plus ^132^ to infer best-fit models of nucleotide evolution for each partition and calculated nodal support using 1,000 ultrafast bootstrap replicates. To further examine node support, we calculated gene and site concordance factors, which measure the number of decisive gene trees and sites that support a given node ^133^. Finally, we used custom scripts (https://www.robertlanfear.com/blog/files/concordance_factors.html) to plot the relationships of gene and site concordance factors, ultrafast bootstrap support values, and log-transformed branch lengths to assess their relationships. We conducted all phylogenetic analyses on the Yale High Performance Computing Cluster McCleary.

### Time Calibration

Using a Bayesian node-dating protocol in *BEAST* v. 2.6.7 ^134,135^, we time-calibrated the maximum likelihood phylogeny generated from analysis of the 75% complete UCE matrix as a single concatenated alignment. We randomly subsampled our UCE set for three sets of 50 UCE loci, each of which was inputted into the BEAUTi terminal to construct input xml files for analyses in *BEAST 2* with tree, substitution, and clock models linked across loci. We used a general-time-reversible (GTR) model of nucleotide evolution with the gamma among-site rate variation parameter, a relaxed log-normal molecular clock, and the *BEAST* implementation of the Fossilized Birth-Death (FBD) Model ^136^. We specified as 0.185, which is the approximate number of eupercarian species sampled as of September 2025 ^77^, the diversification rate prior to 0.05 (bounds of 0.0 and 1.0), which is the approximate background rate estimated for *Acanthomorpha* in a previous study ^61^, and the origin prior to 145.0 Ma, the approximate posterior age of *Acanthomorpha* estimated in ^61^, with bounds of 125.0 and 165.0 Ma. The upper bound of the origin corresponds to the traditional Barremian-Aptian boundary and the oldest putative acanthomorph body fossils ^137,138^, and the lower bound corresponds to the approximate boundary of the Middle and Late Jurassic, when the oldest crown teleosts appear in the fossil record ^139^. We set priors on the ages of 48 nodes based on a newly updated set of fossil calibrations, the majority of which have placements that are supported by phylogenetic analyses of morphological characters. A full list of fossil calibrations and justifications is included in the Supplementary Information. For each fossil calibration, we set a monophyletic MRCA prior with a lognormal age distribution and modified the shape of the distribution so that 97.5% fell before the age of the corresponding fossil. We ran each UCE set three times independently over 2.0 x 10^8^ generations, with a 1.0 x 10^8^ generation pre-burnin and storing every 5,000 generations, checked for convergence of the posteriors and high effective sample size (ESS) values across sets using *Tracer* v. 1.7 ^140^, combined posterior tree sets sampling every 20,000 generations and with a burnin of 75% on each in *LogCombiner* v. 2.6.7, and summarized the phylogenies with median node heights in *TreeAnnotator* v. 2.6.6. We ran all time-calibration analyses on the Yale High Performance Computing Cluster McCleary.

### Discrete Trait Datasets

We conducted a comprehensive literature search for swim bladder presence across our sample of fish species in *Eupercaria* (Supplementary Information). We next collected data on whether fishes occupied benthic, demersal, and pelagic habitats by expanding the dataset of Friedman et al.^82^ in accordance with their criteria for determining habitat occupation and to remain consistent with the literature:^141^ benthic species spend the vast majority of their adult life stages in contact with the substrate, demersal species interact with the substrate but also move within the water column, and pelagic species are found in open water and rarely, if ever, contact the substrate. We classified fish species in our phylogeny as shallow (0-199 m depth), intermediate (200-999 m depth), or deep (>1,000 m depth) by expanding the dataset of Friedman and Muñoz^44^ and applying their criteria for state coding. For both the depth and water column ecology traits, we alternatively coded species as being benthic and non-benthic, and deep and non-deep, for the purposes of downstream analyses. Finally, we coded salinity type as freshwater, brackish, marine, or a combination of the three, from the online data source FishBase,^142^ following previous studies.^122^ For the Root Effect dataset, we discretized the data into two states: Root Effect less than or greater than 20% for at least one hemoglobin. We discretized this trait because of its known within-individual and intraspecific variability^91^ and because a Root Effect of 20% has previously been considered a cutoff for the magnitude of this phenomenon ^42^. We were able to score 141 (11%) tips from all taxonomic orders in *Eupercaria* for the Root Effect trait, 1,333 (100%) tips for the habitat occupancy and salinity traits, and 1,300 (98%) species for the presence of the swim bladder.

### Exploring the body shape morphospace of fishes lacking a swim bladder

A proxy for habitat preference is body shape; fishes with vertically flattened bodies tend to be benthic specialists ^82^. We examined whether the loss of the swim bladder was associated with the evolution of novel body shapes across *Eupercaria* using two methods: phylogenetic logistic regressions and principal components analysis. For n=723 species in our time-calibrated phylogeny (54%), we used the *phyloglm* function in the R package *phylolm* v. 2.1 ^143^ to conduct a phylogenetic logistic regression between fish aspect ratio (maximum body width: maximum body depth) and swim bladder presence based on data from the FishShapes v. 1 ^144^ dataset. Next, we ran a phylogenetic principal components analysis of six continuous measurements of body shape and mass from FishShapes v. 1 ^144^ using the R packages *phytools* v. 2 ^145^ and *geiger* v. 2.0.11 ^146^. Although the FishShapes v. 1 dataset includes nine continuous traits, we were only able to include six owing to the absence of data for three for *Mola mola*.

### Ancestral State Reconstructions and Lineage Through Time Plots

We conducted ancestral state reconstructions of the continuous and discrete character datasets using the R package *phytools* v. 2.0 ^145^. For the traits related to polar and pelagic transitions, the Root Effect trait, and the habitat occupancy trait, we assessed the fit of models of character evolution in which all transition rates were equal (ER) and a model where all transition rates differed (ARD). For the swim bladder trait, we tested the fit of these two models, a model following Dollo’s Law of irreversibility in which only transitions from state 0 to state 1 were allowed, and a reverse irreversible evolution model in which only transitions from state 1 to state 0 were allowed. We used Akaike Information Criterion (AIC) scores to assess the best-fitting model in each case (Supplementary Information). For the swim bladder dataset, we used a reverse Dollo model even though the ER model was the best fit. This is because the swim bladder and its associated vascular anatomy and physiochemical systems comprise a complex trait, and there is no evidence among fishes for the ability to regain a lost swim bladder; this feature conforms to Dollo’s Law of Irreversibility. Evidence for this comes from the observation that the swim bladder varies considerably in anatomy across different lineages that possess a vestigial organ, including closely related species, and the large number of mutants within species that degenerate the organ.^147,148^ For the habitat ancestral state reconstruction, we treated habitat as a polymorphic character using the *fitpolyMk* function, an ARD transition rate model, and the root prior distribution **π** from FitzJohn et al.^149^ After assessing model fit, we generated 1,000 stochastic mappings using the best-fit model and summarized them in a single ancestral state reconstruction. To chart the number of swim bladder losses through time, we deployed custom scripts in *phytools* to calculate the average number of losses across the 1,000 mappings. For comparative purposes, we also produced a lineage-through time plot for the *Eupercaria* tree and used the posterior tree set obtained from *LogCombiner* to calculate 95% confidence intervals and a median lineage-through-time curve. For the continuous trait data, we used the *contMap* function to generate a continuous character ancestral state reconstruction for aspect ratio, which is given as the ratio between maximum body width and maximum body depth.

### Swim Bladder Loss Through Time

To explicitly test how sensitivity to estimated divergence times and species sampling modifies the inferred tempo of swim bladder losses, we constructed density plots of the ages of nodes and branches that correspond to swim bladder losses across our posterior time-calibrated phylogeny sets. Whereas we sample nearly all taxonomic families and most genera of *Eupercaria* in our time-calibrated phylogeny, species diversity is far more complete for some genera, such as *Sebastes* and *Scarus*, than for others, such as *Haemulon* and *Lutjanus*. A solution to this is to assess the timescale of single origins of discrete traits through time. We annotated lists of nodes where each swim bladder loss was inferred to occur in our ancestral state reconstruction. Next, we wrote custom R scripts with the aid of *Claude* Opus 4.7 on the Yale Clarity Platform to collect the ages of these nodes across 10,000 trees randomly selected from our posterior time tree set. We used *ggplot2* ^150^ to produce density curves for these ages to represent the probability densities of swim bladder losses through time. We ran this procedure twice to account for uncertainty in the timing of swim bladder losses by modifying our node sampling approach. In the first case, we selected ages for the nodes corresponding to the crown clades that we inferred ancestrally lost swim bladders. In the second, we selected nodes subtending the stems of the crown clades that we inferred ancestrally lost swim bladders. In this way, we were able to infer whether our density curve of swim bladder losses was robust to whether we reconstruct losses to occur at the base or the top of each branch leading to each crown clade of interest.

### Trait Coevolution

To infer evolutionary associations between swim bladder loss and transitions to benthic and deep-sea habitats, we fit custom models using the *fitMk* function in *phytools* v. 2^145^ that modeled the evolution of these trait pairs as associated or unassociated. For these analyses, we binarized the water column and depth traits to reflect the categories of interest: benthic versus non-benthic, and deep (1000+ m) versus shallow-intermediate (0-999 m). Custom model fitting was necessitated by our wish to treat swim bladder loss as irreversible following our ancestral state reconstructions. We compared the fit of custom models in which swim bladder presence and depth or swim bladder presence and ecology coevolved with models where they did not, with swim bladder re-evolution restricted in all cases. We also compared the fit of these custom models to classic Pagel models of correlated and uncorrelated discrete binary trait evolution fitted using the fitPagel() function in *phytools* v. 2. Finally, we built custom models of joint trait evolution where swim bladder loss was treated as irreversible for the three-state versions of the water column and depth traits and then used chord diagrams to examine the number of transitions between states. For these comparisons and analyses, we used likelihood ratio tests. We developed scripts for correlated trait evolution and checked them with the aid of *Claude* Sonnet 5.0 on the Yale Clarity computing platform.

### Binary and Hidden State Associated Diversification Rates

Swim bladder loss is associated with some of the most famous examples of evolutionary radiation, which features the rapid diversification of often phenotypically and ecologically diverse lineages from a single common ancestor ^14,15,19,20,151,152^, across ray-finned fishes, including the Antarctic icefishes ^83^, North American darters ^56^, and deep-sea anglerfishes ^45^. To assess the effects of swim bladder loss on diversification across *Eupercaria*, we used the R packages *diversitree* 0.10-1 ^153^ and *hisse* v. 2.1.11 ^154^ to fit models of trait-associated diversification rates across our time-calibrated phylogeny. For all analyses, we removed outgroups. We fit models of irreversible trait evolution (no secondary gains of swim bladders) in *diversitree* and estimated posterior distributions for diversification rates associated with the presence or absence of the swim bladder after running an MCMC chain for 10,000 generations with sampling every 100 generations, discarding the first 1000 samples as burn-in, and checking for convergence by examining plots of log-likelihood scores per generation.

Next, we ran HiSSE model testing. After building models of trait evolution where re-evolving the swim bladder was not allowed, we assessed the fit of four models of trait-associated diversification rate: a dull null model in which the turnover and extinction fractions were the same across all states, a binary state-dependent speciation and extinction (BiSSE) model in which turnover and extinction fractions were allowed to between lineages that possessed or lacked a swim bladder, a character-dependent hidden state speciation and extinction (HiSSE CD) model in which turnover and extinction fractions were allowed to vary between both lineages that possessed or lacked a swim bladder *and* between hidden states introduced into the analysis, and finally a character-independent hidden state speciation and extinction (HiSSE CID) model where turnover and extinction fractions were allowed to vary across hidden states, but not across known states. We compared the fit of these four models using AIC scores. Finally, we used the *MarginReconHiSSE* function to jointly reconstruct swim bladder evolution under the forward irreversible model (i.e., swim bladders were not allowed to re-evolve after being lost) and estimate trait-associated diversification rate along the phylogeny. We ran identical BiSSE and HiSSE model selection analyses on the binary depth and water column ecology datasets, except we did not use custom matrix construction to treat them as irreversible characters.

As a second test of how sampling might bias our results, we subsampled our phylogeny for one species per genus and then re-ran BiSSE diversification rate analysis. For *Labridae*, we recognized genera delimited in a recent phylogenetic rank-free taxonomy. We used a sampling fraction of 0.39, which is the proportion of genera in our sample (n=497) divided by the total number reported for *Eupercaria* as of December 2025. We then compared the estimated diversification rates to those found using the full phylogeny.

### Trait-Associated Diversification Through Time

To estimate trait-associated diversification rates through time and assess differences in these rate curves, we deployed a recently developed protocol in the R package *deepSTRAPP* v. 1.0 ^85^ to estimate variation between trait-associated diversification rates pulled from ancestral state reconstructions and lineage diversification rates. Our ancestral state reconstruction protocol was the same as for the reconstruction of swim bladder loss as an irreversible trait and the binary water column ecology and depth traits as reversible traits (see “Ancestral State Reconstructions and Lineage Through Time Plots”). Next, we used *BAMM* ^155^ to estimate lineage diversification rates across our phylogeny. In the R package *BAMMTools* v. 2.1.12 ^155^ piped through *deepSTRAPP* v. 1.0 ^85^, we estimated priors along our input time-calibrated phylogeny using the *setBAMMpriors* function, then ran an MCMC chain for 1.0 x 10^7^ generations sampling every 1000 generations. We confirmed convergence of the posteriors, burned in the run in R, then plotted diversification rates through time for the entirety of *Eupercaria* and for selected sublineages. Next, we used functions in *deepSTRAPP* to run STRAPP tests, which test for differences in diversification rates across trait states through time, calculated for five-million-year intervals. From these analyses, we obtained p-values for trait-associated diversification rate differences at each interval. We used Mann-Whitney-Wilcoxon tests to quantify rate differences across trait states and plotted diversification rates by state for individual intervals. Finally, we re-ran these analyses using the reduced-representation phylogeny to test for how sampling bias might change rate estimates.

### Root Effect Analysis

We compiled data on Root effect magnitude, defined as the decrease in percent oxygen saturation in blood during consecutive decreases in pH in the presence of ATP (following di Prisco et al.^156^). We used the package phytools to conduct a continuous character ancestral state reconstruction using the command *contMap*. We also used the R package *geiger* to estimate mean subclade disparity through time in Root effect magnitude for a pruned version of our phylogeny that only included notothenioids.

## Data Availability

All data needed to replicate the results of this study are available in the Supplementary Information or the repository associated with this article: https://doi.org/10.60600/YU/BT163K.

## Code Availability

All data needed to replicate the results of this study are available in the repository associated with this article under the ‘Scripts’ directory: https://doi.org/10.60600/YU/BT163K.

## Supporting information

Supplemental Text, Tables, Figures, and References

## Acknowledgements.

The authors thank members of the Near and Muñoz Labs for discussions related to this manuscript, as well as the following people and collections for sharing tissues sequenced as part of this effort and previous ones integrated into this study: Academia Sinica (Taipei), Academy of Natural Sciences (Philadelphia), American Museum of Natural History (New York), Australian Museum (Sydney), Australian National Fish Collection (Hobart), Biodiversity Research Museum (Taipei), Burke Museum of Natural History and Culture (Seattle), California Academy of Sciences (San Francisco), Cornell University Museum of Vertebrates (Ithaca), Field Museum of Natural History (Chicago), G. Hoffmann (UCSB), Harvard Museum of Comparative Zoology (Cambridge), Hokkaido University Museum (Sapporo), Illinois Natural History Survey (Champaign), Kagoshima University Museum (Korimoto), Kyoto University Museum, L. Liggins, Louisiana Museum of Natural History (Baton Rouge), M. Miya, Mie University Fish Collection of the Fisheries Research Laboratory (Shima), Museo Nacional de Ciencias Naturales (Madrid), Museo Nacional de Historia Natural (Santiago), Museum of New Zealand Te Papa Tongarewa (Wellington), Museum Victoria (Melbourne), Museums and Art Galleries of the Northern Territory (Darwin), National Museum of Natural History (New Delhi), National Museum of Nature and Science (Tokyo), Natural History Museum and Institute (Chiba), Natural History Museum of Los Angeles County, North Carolina Museum of Natural Sciences (Raleigh), O. Radchenko, Peking University, Queensland Museum (Brisbane), R. Roberston, Royal Ontario Museum (Toronto), S. Klanten, Scripps Institution of Oceanography (La Jolla), Seikai National Fisheries Research Institute (Nagasaki), Smithsonian National Museum of Natural History (Washington, DC), South African Institute for Aquatic Biodiversity (Grahamstown), Southeastern Louisiana University Museum of Biology (Hammond), Universidade Estadual Paulista (São Paulo), Universitetsmuseet i Bergen (Hordaland), University of Copenhagen Zoological Museum, University of Florida Museum of Natural History (Gainesville), University of Kansas Biodiversity Institute (Lawrence), University of Tennessee David A. Etnier Ichthyological Collection (Knoxville), University of Tokyo Ocean Research Institute, and Yale Peabody Museum (New Haven). We thank G. Watkins-Colwell for help with tissue collection processing.

---

**Extended Data Figure 1.**
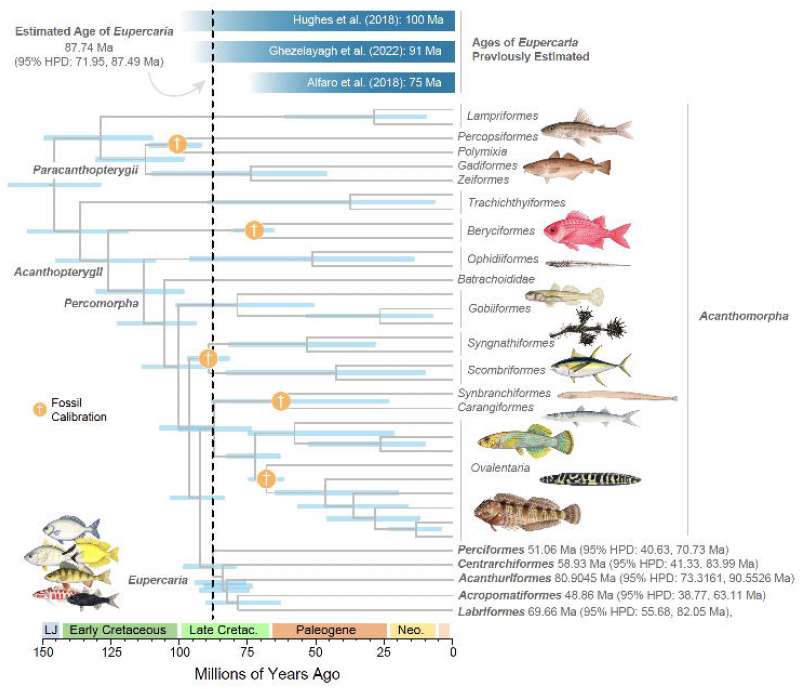
Evolutionary Origin of *Eupercaria*. Shown is the time-calibrated phylogeny from Bayesian node dating analysis of three sets of randomly 50 UCE loci. Major lineages of *Eupercaria* are labeled and shown alongside outgroup lineages. Important clades consisting of taxonomic orders are labeled. The approximate ages of *Eupercaria* estimated in previous studies, at top, are provided as a comparison to our estimate. Also provide are the median and 95% highest posterior density intervals (HPDs) of divergence times for major lineages in *Eupercaria*. The light orange dots at nodes with dagger (†) symbols denote the placement of fossil calibrations; ingroup calibrations are not shown. Illustrations are by Julie Johnson (lifesciencestudios.com).

**Extended Data Figure 2.**
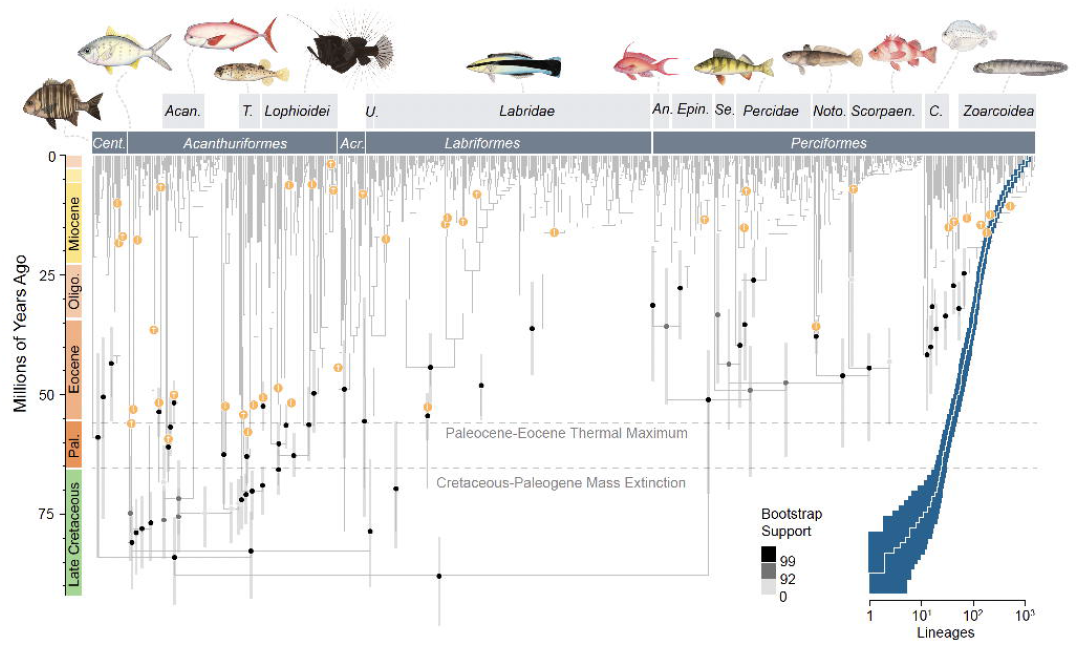
A Time-Calibrated Phylogeny of *Eupercaria* Based on Genomic Data. Shown is the time-calibrated phylogeny from Bayesian node dating analysis of three sets of randomly 50 UCE loci. Major lineages of *Eupercaria* are labeled. Abbreviations, from left to right, are: *Cent*., *Centrarchiformes*, *Acan*., *Acanthuroidei*, *T. Tetraodontoidei*, *U. Uranoscopoidei*, *An.*, *Anthiadidae*, *Epin*., *Epinephelidae*, *Se.*, *Serranidae*, *Noto.*, *Notothenioidei*, *Scorpaen*., *Scorpaenoidea*, *C*., *Cottoidea*. Dots at nodes indicate ultrafast bootstrap support values. Grey bars at nodes indicate 95% highest posterior density intervals of divergence times. The chartreuse dots at nodes with dagger (†) symbols denote the placement of fossil calibrations; outgroup calibrations are not shown. The plot at the right is the log-lineage through time plot, with 95% confidence intervals in dark blue computed from 1,000 trees using *phytools*. Illustrations are by Julie Johnson (lifesciencestudios.com).

**Extended Data Figure 3.**
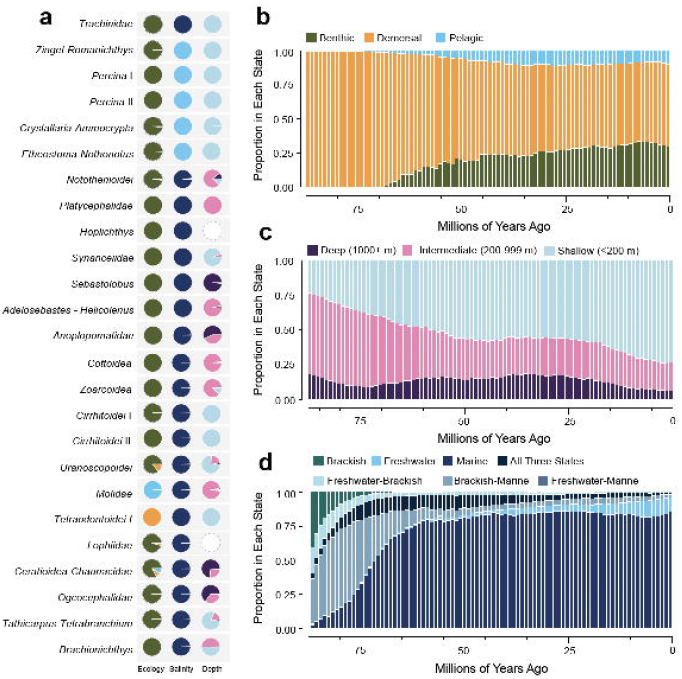
Major Transitions Across Eupercarian Phylogeny. (a) Ancestral state recontructions for water column occupation, depth, and salinity level for the nodes where swim bladder lossess are reconstructed to have occurred. Colors for states match labels in (b-d). Proportion of lineages in each state through time in million-year-intervals for water column ecology (b), depth (c), and salinity level (d).

**Extended Data Figure 4.**
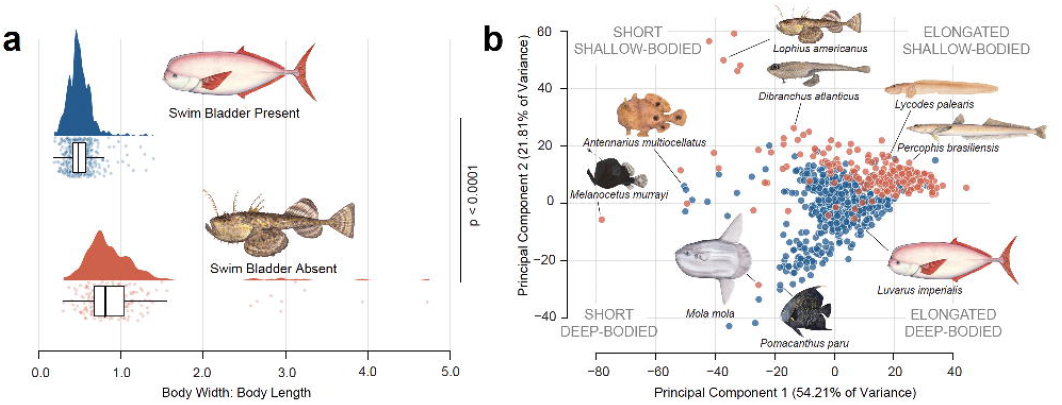
Body Shape Evolution Associated With Swim Bladder Loss. (a) Raincloud plot comparing body shape aspect ratios for fishes with and without swim bladders. Swim bladder loss and increasing flatness are significantly associated. (b) Plot of the first two principal components from phylogenetic principal components analysis of six continuous measurements related to body shape, with species colored by the presence of the swim bladder. Note that most species lacking a swim bladder cluster in the upper right corner of the morphospace, which corresponds to elongated, shallow bodies. Illustrations are by Julie Johnson (lifesciencestudios.com).

**Extended Data Figure 5.**
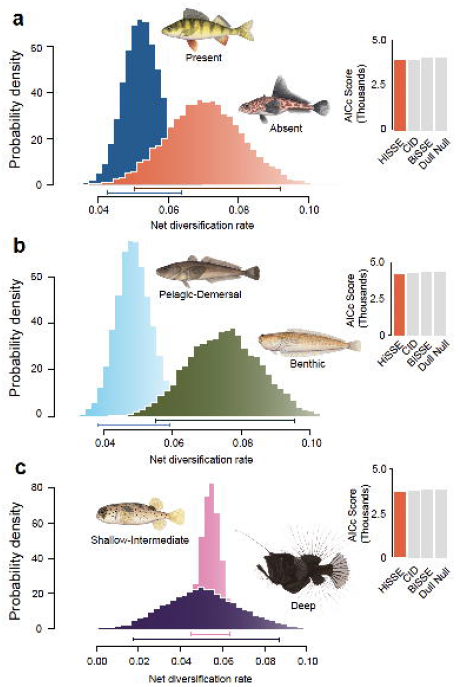
Binary (BiSSE) and Hidden State (HiSSE) Speciation and Extinction Sensitivity Analyses. Trait-associated diversification rate distributions estimated under BiSSE using an irreversible trait model, and HiSSE model fit, from analyses of the pruned tree (single species per genus sampled). Analyses are of (a) swim bladder presence, (b) water column ecology state, and (c) depth state. In the distribution plots, bars below distributions indicate 95% highest posterior density intervals. The inset bars show AICc scores for each of four models tested in the HiSSE framework, with the best-fit in red. Illustrations are by Julie Johnson (lifesciencestudios.com).

**Extended Data Figure 6.**
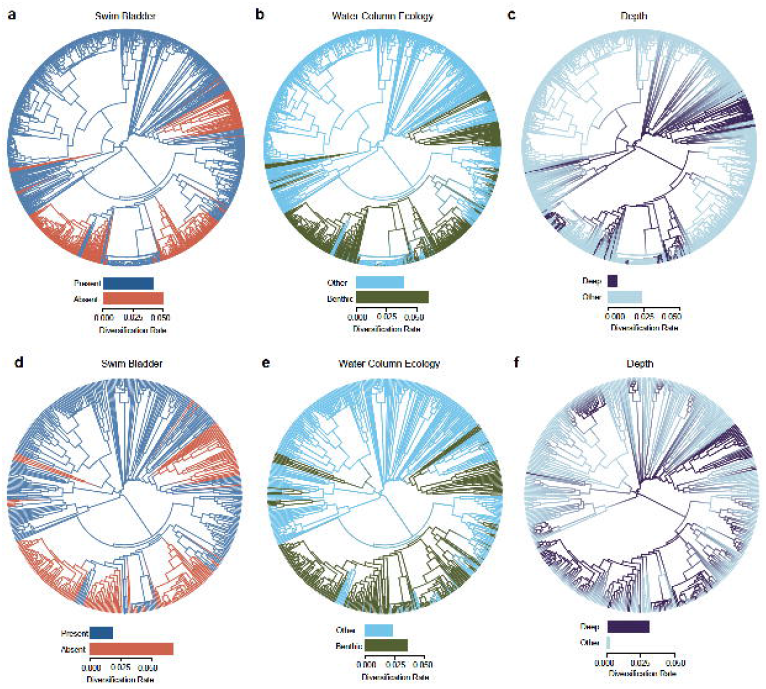
HiSSE Ancestral State Reconstruction and Diversification Rate Estimation. Full marginal ancestral state reconstructions and trait-associated net diversification rate estimates using the best fit HiSSE model for (a, d) swim bladder presence, (b, e) water column ecology state, and (c, f) depth state. Analyses of the full phylogeny are in (a, b, c), and analyses of the subsampled phylogeny are in (d, e, f). The estimated trait-associated diversification rates are presented in the barplots below each tree.

**Extended Data Figure 7.**
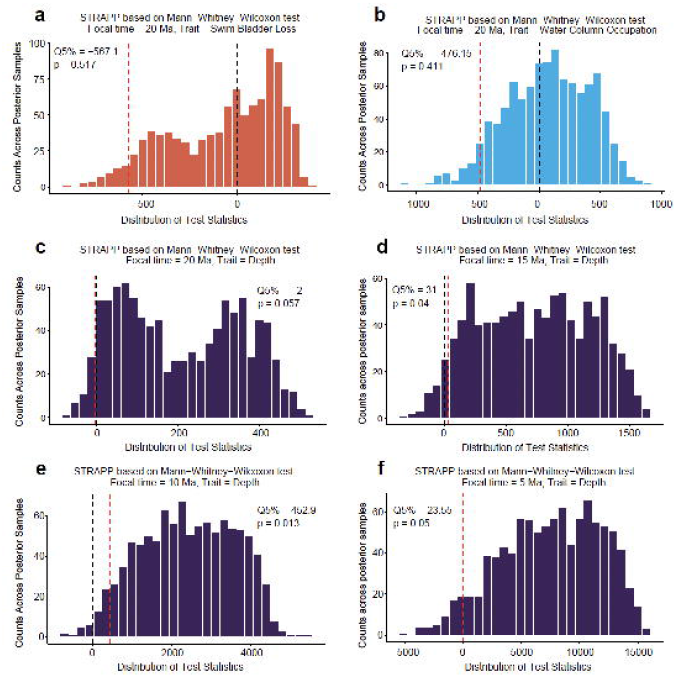
Statistical Tests of Significant Differences in Diversification Rates Through Time. Shown are the STRAPP analysis results based on Mann-Whitney-Wilcoxon tests of differences in trait-associated diversification rates across 1,000 posterior samples from the BAMM analyses at selected time steps for the curves shown in Figure 3b. In each plot, the dotted black line denotes the expected value under the hypothesis that diversification rates do not differ among states, and the red line represents the threshold at which 95% of observed STRAPP test statistics exhibit a higher-than expected value than the null hypothesis would suggest. Distributions are of (a) swim bladder presence-associated rates from analysis of the full phylogeny at 20 Ma, (b) water column ecology (benthic vs. others) associated rates from analysis of the full phylogeny at 20 Ma, and depth-associated rates from analysis of the full phylogeny at (c) 20 Ma, (d) 15 Ma, (e) 10 Ma, and (f) 5 Ma.

**Extended Data Figure 8.**
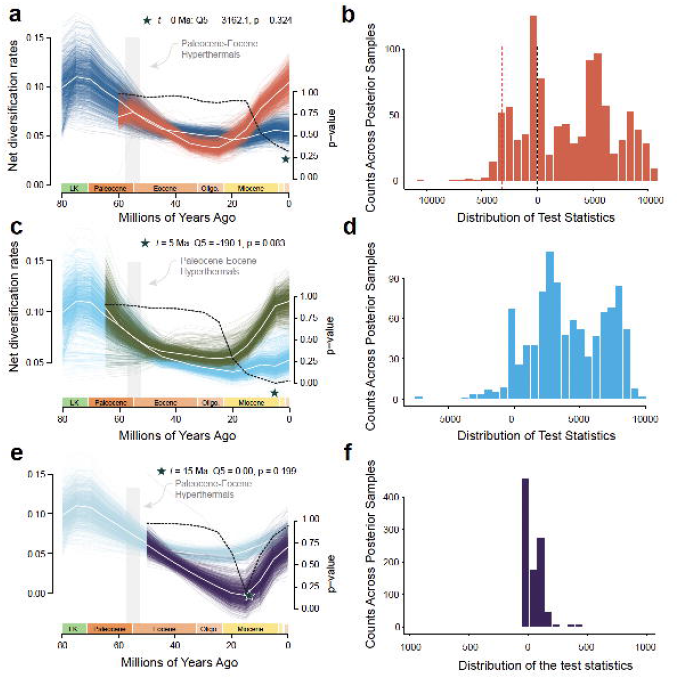
deepSTRAPP Sensitivity Analyses. Trait-associated diversification rates through time estimated using *deepSTRAPP* from analyses of the pruned tree (single species per genus sampled). Mean rate curves are highlighted among the estimated posterior rate curves. The dotted line denotes the p-values from STRAPP tests for differences in trait-associated diversification rates across 5 million year intervals. Analyses are of (a) swim bladder presence, (b) water column ecology state, and (c) depth state. Illustrations are by Julie Johnson (lifesciencestudios.com).

**Extended Data Figure 9.**
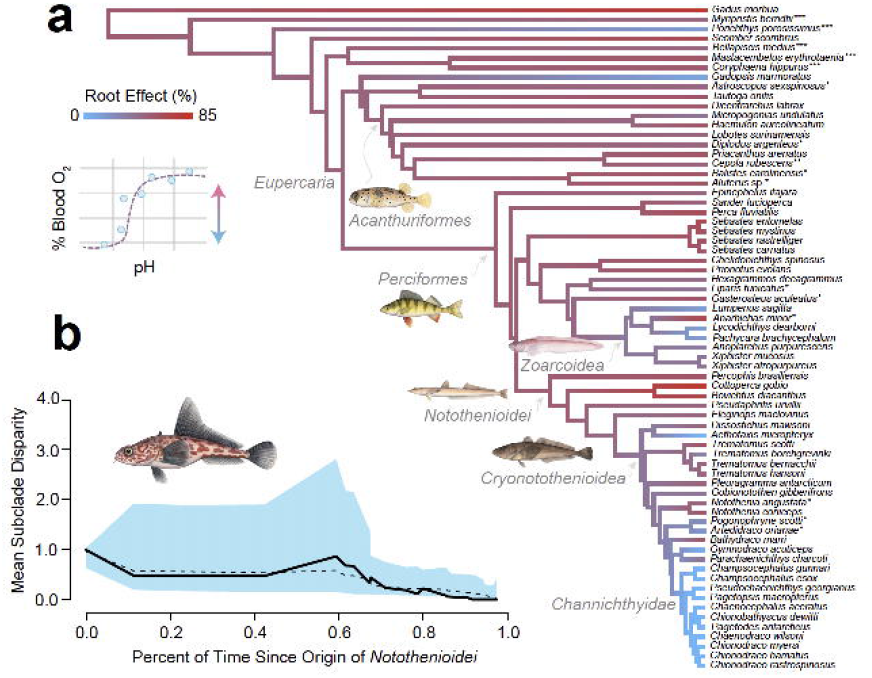
Loss of the Root Effect During Polar Fish Radiations. (a) Complete ancestral state reconstruction of Root effect magnitude across the pruned phylogeny. The inset shows the value used in the ancestral state reconstruction, which is the magnitude of blood oxygen saturation percent decrease across successive pH values. The gathered data corresponds to the biggest decrease in the oxygen saturation curve between pH 8 and pH 6, following previous studies.^156^ (b) Disparity-through-time plot for Root effect magnitude in *Notothenioidei*. The solid black line shows the average disparity through time observed along the tree, the dotted black line describes the expected disparity under a Brownian motion model, and the shaded grey region is the 95% confidence interval for the expected disparity through time under a Brownian motion model.

## References.

1. Heard, S. B. & Hauser, D. L. Key evolutionary innovations and their ecological mechanisms. Historical Biology 10, 151–173 (1995).

2. Miller, A. H., Stroud, J. T. & Losos, J. B. The ecology and evolution of key innovations. Trends in Ecology & Evolution 38, 122–131 (2023).

3. Wei, Z., et al. The contingent advantage of photosymbiosis in coral evolution. Proceedings of the National Academy of Sciences 123, e2532242123 (2026).

4. Peoples, N., Burns, M. D., Mihalitsis, M. & Wainwright, P. C. Evolutionary lability of a key innovation spurs rapid diversification. Nature 639, 962–967 (2025).

5. McGee, M. D., et al. A pharyngeal jaw evolutionary innovation facilitated extinction in Lake Victoria cichlids. Science 350, 1077–1079 (2015).

6. Darwin, C. On the Origin of Species by Means of Natural Selection, or the Preservation of Favoured Races in the Struggle for Life. (John Murray, London, 1859).

7. Title, P. O., et al. The macroevolutionary singularity of snakes. Science 383, 918–923 (2024).

8. Brownstein, C. D., et al. Synergistic innovations enabled the radiation of anglerfishes in the deep open ocean. Current Biology 34, 2541–2550.e4 (2024).

9. McGee, M. D., et al. A pharyngeal jaw evolutionary innovation facilitated extinction in Lake Victoria cichlids. Science 350, 1077–1079 (2015).

10. Borstein, S. R., Hammer, M. P., O’Meara, B. C. & McGee, M. D. The macroevolutionary dynamics of pharyngognathy in fishes fail to support the key innovation hypothesis. Nature Communications 15, 10325 (2024).

11. Marshall, N. B. Swimbladder structure of deep-sea fishes in relation to their systematics and biology. Fishery Bulletin 73, 95–109 (1975).

12. DeVries, A. L. & Eastman, J. T. Physiology and ecology of notothenioid fishes of the Ross Sea. Journal of the Royal Society of New Zealand 11, 329–340 (1981).

13. Verde, C., Vergara, A., Giordano, D., Mazzarella, L. & Prisco, G. D. The Root effect – a structural and evolutionary perspective. Antarctic Science 19, 271–278 (2007).

14. Gavrilets, S. & Losos, J. B. Adaptive Radiation: Contrasting Theory with Data. Science 323, 732–737 (2009).

15. Schluter, D. The Ecology of Adaptive Radiation. (OUP Oxford, 2000).

16. Wainwright, P. C. Functional Versus Morphological Diversity in Macroevolution. Annual Review of Ecology, Evolution, and Systematics 38, 381–401 (2007).

17. Stephen Jay Gould. The Structure Of Evolutionary Theory. (2002).

18. Burress, E. D. & Muñoz, M. M. Ecological Opportunity from Innovation, not Islands, Drove the Anole Lizard Adaptive Radiation. Systematic Biology 71, 93–104 (2022).

19. Gillespie, R. G., et al. Comparing Adaptive Radiations Across Space, Time, and Taxa. Journal of Heredity 111, 1–20 (2020).

20. Stroud, J. T. & Losos, J. B. Ecological Opportunity and Adaptive Radiation. Annual Review of Ecology, Evolution, and Systematics 47, 507–532 (2016).

21. Liem, K. F. Evolutionary Strategies and Morphological Innovations: Cichlid Pharyngeal Jaws. Systematic Zoology 22, 425–441 (1973).

22. Blount, Z. D., Borland, C. Z. & Lenski, R. E. Historical contingency and the evolution of a key innovation in an experimental population of Escherichia coli. Proceedings of the National Academy of Sciences 105, 7899–7906 (2008).

23. Roberts-Hugghis, A. S., Burress, E. D., Lam, B. & Wainwright, P. C. The cichlid pharyngeal jaw novelty enhances evolutionary integration in the feeding apparatus. Evolution 77, 1917–1929 (2023).

24. Alencar, L. R. V., et al. Opportunity begets opportunity to drive macroevolutionary dynamics of a diverse lizard radiation. Evolution Letters 8, 623–637 (2024).

25. Chen, H., et al. The rise of polyploids during environmental upheaval. Cell 0, (2026).

26. Albertson, R. C. & Kocher, T. D. Genetic and developmental basis of cichlid trophic diversity. Heredity 97, 211–221 (2006).

27. Walker, J. A. A general model of functional constraints on phenotypic evolution. The American Naturalist 170, 681–689 (2007).

28. Futuyma, D. J. & Moreno, G. The Evolution of Ecological Specialization. Annual Review of Ecology and Systematics 19, 207–233 (1988).

29. Field, D. J., et al. Early Evolution of Modern Birds Structured by Global Forest Collapse at the End-Cretaceous Mass Extinction. Current Biology 28, 1825–1831.e2 (2018).

30. Poulson, T. L. & White, W. B. The Cave Environment. Science 165, 971–981 (1969).

31. Wynne, J. J. Cave Biodiversity: Speciation and Diversity of Subterranean Fauna. (JHU Press, 2022).

32. Wright, N. A., Steadman, D. W. & Witt, C. C. Predictable evolution toward flightlessness in volant island birds. Proceedings of the National Academy of Sciences 113, 4765–4770 (2016).

33. Sackton, T. B., et al. Convergent regulatory evolution and loss of flight in paleognathous birds. Science 364, 74–78 (2019).

34. Maderspacher, F. Flightless birds. Current Biology 32, R1155–R1162 (2022).

35. Chippindale, P. T., Bonett, R. M., Baldwin, A. S. & Wiens, J. J. Phylogenetic Evidence for a Major Reversal of Life-History Evolution in Plethodontid Salamanders. Evolution 58, 2809–2822 (2004).

36. Bailey, N. W., Pascoal, S. & Montealegre-Z, F. Testing the role of trait reversal in evolutionary diversification using song loss in wild crickets. Proceedings of the National Academy of Sciences 116, 8941–8949 (2019).

37. Fänge, R. Physiology of the swimbladder. Physiological Reviews 46, 299–322 (1966).

38. Pelster, B. Using the swimbladder as a respiratory organ and/or a buoyancy structure— Benefits and consequences. Journal of Experimental Zoology Part A: Ecological and Integrative Physiology 335, 831–842 (2021).

39. Berenbrink, M. Historical reconstructions of evolving physiological complexity:O2 secretion in the eye and swimbladder of fishes. Journal of Experimental Biology 210, 1641–1652 (2007).

40. Berenbrink, M., Koldkjær, P., Kepp, O. & Cossins, A. R. Evolution of Oxygen Secretion in Fishes and the Emergence of a Complex Physiological System. Science 307, 1752–1757 (2005).

41. Brittain, T. Root effect hemoglobins. Journal of Inorganic Biochemistry 99, 120–129 (2005).

42. Pelster, B. & Weber, R. E. The Physiology of the Root Effect. in Advances in Comparative and Environmental Physiology: Volume 8 (eds Castellini, M. A. et al.) 51–77 (Springer, Berlin, Heidelberg, 1991). doi:10.1007/978-3-642-75900-0_2.

43. Rummer, J. L., McKenzie, D. J., Innocenti, A., Supuran, C. T. & Brauner, C. J. Root Effect Hemoglobin May Have Evolved to Enhance General Tissue Oxygen Delivery. Science 340, 1327–1329 (2013).

44. Rabosky, D. L., et al. An inverse latitudinal gradient in speciation rate for marine fishes. Nature 559, 392–395 (2018).

45. Pietsch, T. W. Oceanic Anglerfishes: Extraordinary Diversity in the Deep Sea. (University of California Press, 2009).

46. Pietsch, T. W. The Genera of Frogfishes (Family Antennariidae). Copeia 1984, 27–44 (1984).

47. Friedman, M. The evolutionary origin of flatfish asymmetry. Nature 454, 209–212 (2008).

48. Harrington, R. C., et al. Phylogenomic analysis of carangimorph fishes reveals flatfish asymmetry arose in a blink of the evolutionary eye. BMC Evolutionary Biology 16, 224 (2016).

49. Arbour, J. H. & Stanchak, K. E. The little fishes that could: smaller fishes demonstrate slow body size evolution but faster speciation in the family Percidae. Biological Journal of the Linnean Society 134, 851–866 (2021).

50. Carlson, R. L. & Wainwright, P. C. The ecological morphology of darter fishes (Percidae: Etheostomatinae). Biological Journal of the Linnean Society 100, 30–45 (2010).

51. Near, T. J., et al. Phylogeny and Temporal Diversification of Darters (Percidae: Etheostomatinae). Systematic Biology 60, 565–595 (2011).

52. McCune, A. R. & Carlson, R. L. Twenty ways to lose your bladder: common natural mutants in zebrafish and widespread convergence of swim bladder loss among teleost fishes. Evolution & Development 6, 246–259 (2004).

53. Bista, I., et al. Genomics of cold adaptations in the Antarctic notothenioid fish radiation. Nat Commun 14, 3412 (2023).

54. Daane, J. M., et al. Historical contingency shapes adaptive radiation in Antarctic fishes. Nature Ecology & Evolution 3, 1102 (2019).

55. Near, T. J., et al. Ancient climate change, antifreeze, and the evolutionary diversification of Antarctic fishes. Proceedings of the National Academy of Sciences 109, 3434–3439 (2012).

56. Arbour, J. H. Hitting rock bottom: exaptation, ecological filtering, and the benthopelagic divergence of percid fishes. Evolution 79, 2057–2071 (2025).

57. Page, L. M. & Swofford, D. L. Morphological correlates of ecological specialization in darters. Environmental biology of Fishes 11, 139–159 (1984).

58. Steen, J. B. 10 The swim bladder as a hydrostatic organ. in Fish physiology vol. 4 413–443 (Elsevier, 1970).

59. Denton, E. J. & Marshall, N. B. The buoyancy of bathypelagic fishes without a gas-filled swimbladder. Journal of the Marine Biological Association of the United Kingdom 37, 753–767 (1958).

60. Evans, J. & Page, L. Distribution and Relative Size of the Swim Bladder in Percina, With Comparisons to Etheostoma, Crystallaria, and Ammocrypta (Teleostei: Percidae). Environmental Biology of Fishes - ENVIRON BIOL FISH 66, 61–65 (2003).

61. Ghezelayagh, A., et al. Prolonged morphological expansion of spiny-rayed fishes following the end-Cretaceous. Nature Ecology & Evolution 6, 1211–1220 (2022).

62. Hughes, L. C., et al. Comprehensive phylogeny of ray-finned fishes (Actinopterygii) based on transcriptomic and genomic data. Proceedings of the National Academy of Sciences of the United States of America 115, 6249–6254 (2018).

63. Musilova, Z., et al. Vision using multiple distinct rod opsins in deep-sea fishes. Science 364, 588–592 (2019).

64. Alfaro, M. E., et al. Explosive diversification of marine fishes at the Cretaceous–Palaeogene boundary. Nature Ecology & Evolution 2, 688–696 (2018).

65. Betancur-R, R., et al. Phylogenetic classification of bony fishes. BMC Evolutionary Biology 17, 162 (2017).

66. Dornburg, A. & Near, T. J. The Emerging Phylogenetic Perspective on the Evolution of Actinopterygian Fishes. Annual Review of Ecology, Evolution, and Systematics 52, 427– 452 (2021).

67. Betancur-R, R., et al. The Tree of Life and a New Classification of Bony Fishes. PLOS Currents Tree of Life 10.1371/currents.tol.53ba26640df0ccaee75bb165c8c26288 (2013) doi:10.1371/currents.tol.53ba26640df0ccaee75bb165c8c26288.

68. Liem, K. F. & Sanderson, S. L. The pharyngeal jaw apparatus of labrid fishes: A functional morphological perspective. Journal of Morphology 187, 143–158 (1986).

69. Burress, E. D. & Wainwright, P. C. Adaptive radiation in labrid fishes: A central role for functional innovations during 65 My of relentless diversification. Evolution 73, 346–359 (2019).

70. Brownstein, C. D., Harrington, R. C., Radchenko, O. & Near, T. J. The many origins of extremophile fishes. Proceedings of the Royal Society B: Biological Sciences 292, 20250217 (2025).

71. Brownstein, C. D., et al. Phylogenomics establishes an Early Miocene reconstruction of reef vertebrate diversity. Science Advances 11, eadu6149 (2025).

72. Bogan, S. N., et al. Temperature and Pressure Shaped the Evolution of Antifreeze Proteins in Polar and Deep Sea Zoarcoid Fishes. Molecular Biology and Evolution 42, msaf219 (2025).

73. Wainwright, P. C. & Longo, S. J. Functional Innovations and the Conquest of the Oceans by Acanthomorph Fishes. Current Biology 27, R550–R557 (2017).

74. Konow, N., Bellwood, D. R., Wainwright, P. C. & Kerr, A. M. Evolution of novel jaw joints promote trophic diversity in coral reef fishes. Biological Journal of the Linnean Society 93, 545–555 (2008).

75. Near, T. J., et al. Resolution of ray-finned fish phylogeny and timing of diversification. Proceedings of the National Academy of Sciences 109, 13698–13703 (2012).

76. Melendez-Vazquez, F., et al. Ecological interactions and genomic innovation fueled the evolution of ray-finned fish endothermy. Science Advances 11, eads8488 (2025).

77. Fricke, R., Eschmeyer, W. N. & Laan, R. V. D. ESCHMEYER’S CATALOG OF FISHES: GENERA, SPECIES, REFERENCES. Electronic Version http://researcharchive.calacademy.org/research/ichthyology/catalog/fishcatmain.asp.

78. Eastman, J. T. The nature of the diversity of Antarctic fishes. Polar Biology 28, 93–107 (2005).

79. Mecklenburg, C. W., Møller, P. R. & Steinke, D. Biodiversity of arctic marine fishes: taxonomy and zoogeography. Marine Biodiversity 41, 109–140 (2011).

80. Wang, K., et al. Morphology and genome of a snailfish from the Mariana Trench provide insights into deep-sea adaptation. Nature Ecology & Evolution 3, 823–833 (2019).

81. Xu, W., et al. Chromosome-level genome assembly of hadal snailfish reveals mechanisms of deep-sea adaptation in vertebrates. eLife 12, RP87198 (2023).

82. Friedman, S. T., et al. Body shape diversification along the benthic–pelagic axis in marine fishes. Proceedings of the Royal Society B: Biological Sciences 287, 20201053 (2020).

83. Eastman, J. T. & DeVries, A. L. Buoyancy adaptations in a swim-bladderless Antarctic fish. Journal of Morphology 167, 91–102 (1981).

84. Near, T. J., Kendrick, B. J., William Detrich, H. & Jones, C. D. Confirmation of neutral buoyancy in Aethotaxis mitopteryx DeWitt (Notothenioidei: Nototheniidae). Polar Biology 30, 443–447 (2007).

85. Doré, M., et al. Evolutionary history of ponerine ants highlights how the timing of dispersal events shapes modern biodiversity. Nature Communications 16, 8297 (2025).

86. Miller, E. C., et al. Reduced evolutionary constraint accompanies ongoing radiation in deepsea anglerfishes. Nat Ecol Evol 9, 474–490 (2025).

87. Friedman, S. T. & Muñoz, M. M. A latitudinal gradient of deep-sea invasions for marine fishes. Nature Communications 14, 773 (2023).

88. Hotaling, S., Borowiec, M. L., Lins, L. S. F., Desvignes, T. & Kelley, J. L. The biogeographic history of eelpouts and related fishes: Linking phylogeny, environmental change, and patterns of dispersal in a globally distributed fish group. Molecular Phylogenetics and Evolution 162, 107211 (2021).

89. Hobbs, R. S., Hall, J. R., Graham, L. A., Davies, P. L. & Fletcher, G. L. Antifreeze protein dispersion in eelpouts and related fishes reveals migration and climate alteration within the last 20 Ma. PLOS ONE 15, e0243273 (2020).

90. Kunzmann, A., Caruso, C. & Prisco, G. di. Haematological studies on a high-Antarctic fish: *Bathydraco marri* Norman. Journal of Experimental Marine Biology and Ecology 152, 243–255 (1991).

91. di Prisco, G., et al. Structure and function of hemoglobin in antarctic fishes and evolutionary implications. Polar Biology 10, 269–274 (1990).

92. Sidell, B. D. & O’Brien, K. M. When bad things happen to good fish: the loss of hemoglobin and myoglobin expression in Antarctic icefishes. Journal of Experimental Biology 209, 1791–1802 (2006).

93. Holeton, G. F. Oxygen uptake and circulation by a hemoglobinless antarctic fish (*Chaenocephalus aceratus* Lonnberg) compared with three red-blooded antarctic fish. Comparative Biochemistry and Physiology 34, 457–471 (1970).

94. Cocca, E., et al. Do the hemoglobinless icefishes have globin genes? Comparative Biochemistry and Physiology Part A: Physiology 118, 1027–1030 (1997).

95. Cocca, E., et al. Genomic remnants of alpha-globin genes in the hemoglobinless antarctic icefishes. Proceedings of the National Academy of Sciences of the United States of America 92, 1817–1821 (1995).

96. Rummer, J. L. & Brauner, C. J. Root Effect Haemoglobins in Fish May Greatly Enhance General Oxygen Delivery Relative to Other Vertebrates. PLoS One 10, e0139477 (2015).

97. Bargelloni, L., Marcato, S. & Patarnello, T. Antarctic fish hemoglobins: Evidence for adaptive evolution at subzero temperature. Proceedings of the National Academy of Sciences 95, 8670–8675 (1998).

98. Huey, R. B. & Kingsolver, J. G. Climate Warming, Resource Availability, and the Metabolic Meltdown of Ectotherms. The American Naturalist 194, E140–E150 (2019).

99. Simpson, G. G. The Major Features of Evolution. (New York, Columbia University Press, 1953).

100. Judd, E. J., et al. A 485-million-year history of Earth’s surface temperature. Science 385, eadk3705 (2024).

101. Keller, G., et al. Environmental changes during the Cretaceous-Paleogene mass extinction and Paleocene-Eocene Thermal Maximum: Implications for the Anthropocene. Gondwana Research 56, 69–89 (2018).

102. Speijer, R., Scheibner, C., Stassen, P. & Morsi, A.-M. Response of marine ecosystems to deep-time global warming: A synthesis of biotic patterns across the Paleocene-Eocene thermal maximum (PETM). Austrian Journal of Earth Sciences 105, 6–16 (2012).

103. Pujalte, V., et al. Impact of the Paleocene-Eocene thermal maximum on the evolution of larger foraminifera: a new look at an old problem. Facies 71, 19 (2025).

104. Aze, T. Unraveling ecological signals from a global warming event of the past. Proceedings of the National Academy of Sciences 119, e2201495119 (2022).

105. Tian, S. Y., Yasuhara, M., Huang, H.-H. M., Condamine, F. L. & Robinson, M. M. Shallow marine ecosystem collapse and recovery during the Paleocene-Eocene Thermal Maximum. Global and Planetary Change 207, 103649 (2021).

106. Thomas, E. Cenozoic Mass Extinctions in the Deep Sea; What Disturbs the Largest Habitat on Earth? Division III Faculty Publications 424, (2007).

107. Kennett, J. P. & Stott, L. D. Abrupt deep-sea warming, palaeoceanographic changes and benthic extinctions at the end of the Palaeocene. Nature 353, 225–229 (1991).

108. Arreguín-Rodríguez, G. J., Thomas, E., D’haenens, S., Speijer, R. P. & Alegret, L. Early Eocene deep-sea benthic foraminiferal faunas: Recovery from the Paleocene Eocene Thermal Maximum extinction in a greenhouse world. PLOS ONE 13, e0193167 (2018).

109. McInerney, F. A. & Wing, S. L. The Paleocene-Eocene Thermal Maximum: A Perturbation of Carbon Cycle, Climate, and Biosphere with Implications for the Future. Annual Review of Earth and Planetary Sciences 39, 489–516 (2011).

110. Silva, H. M. A. & Gallo, V. Taxonomic review and phylogenetic analysis of Enchodontoidei (Teleostei: Aulopiformes). Anais da Academia Brasileira de Ciências 83, 483–511 (2011).

111. da Silva, H. M. A. & Gallo, V. Distributional patterns of enchodontoid fishes in the Late Cretaceous. Cretaceous Research 65, 223–231 (2016).

112. Brownstein, C. D. & Near, T. J. Evolutionary origins of the lampriform pelagic radiation. Zoological Journal of the Linnean Society 201, 422–430 (2024).

113. Ribeiro, E., Davis, A. M., Rivero-Vega, R. A., Ortí, G. & Betancur-R, R. Post-Cretaceous bursts of evolution along the benthic-pelagic axis in marine fishes. Proceedings of the Royal Society B: Biological Sciences 285, 20182010 (2018).

114. Eastman, J. T. The Axes of Divergence for the Evolutionary Radiation of Notothenioid Fishes in Antarctica. Diversity 16, 214 (2024).

115. Raia, P., et al. Progress to extinction: increased specialisation causes the demise of animal clades. Scientific Reports 6, 30965 (2016).

116. Burke, K. D., et al. Pliocene and Eocene provide best analogs for near-future climates. Proceedings of the National Academy of Sciences 115, 13288–13293 (2018).

117. Near, T. J. & Thacker, C. E. Phylogenetic classification of living and fossil ray-finned fishes (Actinopterygii). Bulletin of the Peabody Museum of Natural History 65, 3–302 (2024).

118. Near, T. J., Brownstein, C. D., Thacker, C. E. & Wainwright, P. C. Phylogenetic Taxonomy of Wrasses and Parrotfishes (Labridae). Bulletin of the Peabody Museum of Natural History 66, 263–338 (2025).

119. Queiroz, K. de & Cantino, P. *International Code of Phylogenetic Nomenclature (PhyloCode)*. (CRC Press, Boca Raton, 2020). doi:10.1201/9780429446320.

120. Thines, M., et al. Setting scientific names at all taxonomic ranks in italics facilitates their quick recognition in scientific papers. IMA Fungus 11, 25 (2020).

121. Baraf, L. M., Hung, J. Y. & Cowman, P. F. Phylogenomics of marine angelfishes: diagnosing sources of systematic discordance for an iconic reef fish family (F: Pomacanthidae). Systematic Biology syaf016 (2025) doi:10.1093/sysbio/syaf016.

122. Wood, J. E., Harrington, R. C., Ghezelayagh, A., Geiger, M. F. & Near, T. J. Saltwater tolerance and the Holarctic distribution of freshwater percid fishes. Systematic Biology syag057 (2026) doi:10.1093/sysbio/syag057.

123. Glenn, T. C., et al. Adapterama I: universal stubs and primers for 384 unique dual-indexed or 147,456 combinatorially-indexed Illumina libraries (iTru & iNext). PeerJ 7, e7755 (2019).

124. Katoh, K. & Standley, D. M. MAFFT multiple sequence alignment software version 7: improvements in performance and usability. Molecular Biology and Evolution 30, 772–780 (2013).

125. Del Fabbro, C., Scalabrin, S., Morgante, M. & Giorgi, F. M. An Extensive Evaluation of Read Trimming Effects on Illumina NGS Data Analysis. PLoS ONE 8, e85024 (2013).

126. Bolger, A. M., Lohse, M. & Usadel, B. Trimmomatic: a flexible trimmer for Illumina sequence data. Bioinformatics 30, 2114–2120 (2014).

127. Lohse, M., et al. RobiNA: a user-friendly, integrated software solution for RNA-Seq-based transcriptomics. Nucleic Acids Research 40, W622–W627 (2012).

128. Faircloth, B. C. Illumiprocessor - software for Illumina read quality filtering. Brant Faircloth 10.6079/J9ILL (2011).

129. Bankevich, A., et al. SPAdes: a new genome assembly algorithm and its applications to single-cell sequencing. Journal of Computational Biology 19, 455–477 (2012).

130. Tumescheit, C., Firth, A. E. & Brown, K. CIAlign: A highly customisable command line tool to clean, interpret and visualise multiple sequence alignments. PeerJ 10, e12983 (2022).

131. Minh, B. Q., et al. IQ-TREE 2: New Models and Efficient Methods for Phylogenetic Inference in the Genomic Era. Molecular Biology and Evolution 37, 1530–1534 (2020).

132. Kalyaanamoorthy, S., Minh, B. Q., Wong, T. K. F., von Haeseler, A. & Jermiin, L. S. ModelFinder: fast model selection for accurate phylogenetic estimates. Nature Methods 14, 587–589 (2017).

133. Minh, B. Q., Hahn, M. W. & Lanfear, R. New Methods to Calculate Concordance Factors for Phylogenomic Datasets. Molecular Biology and Evolution 37, 2727–2733 (2020).

134. Bouckaert, R., et al. BEAST 2.5: An advanced software platform for Bayesian evolutionary analysis. PLOS Computational Biology 15, e1006650 (2019).

135. Bouckaert, R., et al. BEAST 2: A Software Platform for Bayesian Evolutionary Analysis. PLoS Computational Biology 10, e1003537 (2014).

136. Gavryushkina, A., et al. Bayesian Total-Evidence Dating Reveals the Recent Crown Radiation of Penguins. Systematic Biology 66: 57–73 (2016) doi:10.1093/sysbio/syw060.

137. Ribeiro, A. C., de Mayrinck, D., Bockmann, F. A. & de Pinna, M. The oldest acanthomorph fossil (Actinopterygii, Teleostei) from the Early Cretaceous of Gondwana (Morro do Chaves Formation, Sergipe–Alagoas Basin, NE Brazil). Papers in Palaeontology 12, e70072 (2026).

138. Chen, W.-J., et al. New insights on early evolution of spiny-rayed fishes (Teleostei: Acanthomorpha). Frontiers in Marine Science 1, 53 (2014).

139. Friedman, M. The Macroevolutionary History of Bony Fishes: A Paleontological View. Annual Review of Ecology, Evolution, and Systematics 53, 353–377 (2022).

140. Rambaut, A., Drummond, A. J., Xie, D., Baele, G. & Suchard, M. A. Posterior Summarization in Bayesian Phylogenetics Using Tracer 1.7. Systematic Biology 67, 901– 904 (2018).

141. Santos, E. C., Friedman, S. T. & Martinez, C. M. Distinct evolutionary signatures underlie body shape diversity across deep sea habitats. Evolution 80, 85–96 (2026).

142. Froese, R. & Pauly, D. FishBase. (2010).

143. Ho, L. S. T. et al. phylolm: Phylogenetic Linear Regression. (2024).

144. Price, S. A., et al. FishShapes v1: Functionally relevant measurements of teleost shape and size on three dimensions. Ecology 103, e3829 (2022).

145. Revell, L. J. phytools 2.0: an updated R ecosystem for phylogenetic comparative methods (and other things). PeerJ 12, e16505 (2024).

146. Harmon, L. J., Weir, J. T., Brock, C. D., Glor, R. E. & Challenger, W. GEIGER: investigating evolutionary radiations. Bioinformatics 24, 129–131 (2008).

147. Mok, hin kiu. Coelomic Organs of Perciform Fishes (teleostei). (City University of New York, United States -- New York, 1978).

148. Dallas Evans, J. & Page, L. M. Distribution and Relative Size of the Swim Bladder in Percina, With Comparisons to Etheostoma, Crystallaria, and Ammocrypta (Teleostei: Percidae). Environmental Biology of Fishes 66, 61–65 (2003).

149. FitzJohn, R. G., Maddison, W. P. & Otto, S. P. Estimating Trait-Dependent Speciation and Extinction Rates from Incompletely Resolved Phylogenies. Systematic Biology 58, 595–611 (2009).

150. Valero-Mora, P. M. ggplot2: Elegant Graphics for Data Analysis. Journal of Statistical Software 35, 1–3 (2010).

151. Czekanski-Moir, J. E. & Rundell, R. J. The Ecology of Nonecological Speciation and Nonadaptive Radiations. Trends in Ecology & Evolution 34, 400–415 (2019).

152. Rundell, R. J. & Price, T. D. Adaptive radiation, nonadaptive radiation, ecological speciation and nonecological speciation. Trends in Ecology & Evolution 24, 394–399 (2009).

153. FitzJohn, R. G. *Diversitree*: comparative phylogenetic analyses of diversification in R. Methods in Ecology and Evolution 3, 1084–1092 (2012).

154. Beaulieu, J., O’Meara, B., Caetano, D., Boyko, J. & Vasconcelos, T. hisse: Hidden State Speciation and Extinction. (2023).

155. Rabosky, D. L., et al. BAMMtools: an R package for the analysis of evolutionary dynamics on phylogenetic trees. Methods in Ecology and Evolution 5, 701–707 (2014).

156. di Prisco, G., Eastman, J. T., Giordano, D., Parisi, E. & Verde, C. Biogeography and adaptation of Notothenioid fish: Hemoglobin function and globin–gene evolution. Gene 398, 143–155 (2007).

