## Supplemental Text, Tables, Figures, and References for "The macroevolutionary impact of an innovation reversal in ray-finned fishes"

**Table of Contents.**

Fossil Calibration Justification List ………. Pg. 1

Supplementary Results and Discussion …... Pg. 14

References………………………………….Pg. 17

Figure Captions ……………………………Pg. 25

**Fossil Calibration Justification List.**

†*Cumbaaichthys oxyrhynchus*

**Phylogenetic Placement and Justification:** †*Cumbaaichthys oxyrhynchus* calibrates pan-*Polymixia*; in analyses presented in this paper, this is the MRCA of *Polymixia lowei* and *Aphredoderus sayanus*. See a full justification of this calibration in (1).

**Stratigraphy**: Lacdes Bois, Northwest Territories, Canada: early Turonian, Late Cretaceous ^1^.

**Fossil tip age**: 93.9 Ma (M. Wilson 1977; Mayr et al. 2019). The node calibration was created such that 97.5% of the probability distribution fell before this date.

†*Iridopristis parrisi*

**Phylogenetic Placement and Justification:** †*Iridopristis parrisi* calibrates the MRCA of *Beryx splendens* and *Holocentrus rufus* and is placed as a pan-holocentrid in a recent phylogenetic

analysis of morphological characters ^3^. See a full justification of this calibration in ^4^.

**Stratigraphy:** Main Fossiliferous Layer, Hornerstown Formation, New Jersey, USA: earliest Danian, Paleocene ^3^, approximately 66.02 Ma ^2^.

**Fossil age**: 66.02 Ma. The node calibration was created such that 97.5% of the probability distribution fell before this date.

†*Gasterorhamphosus zuppichini*

**Phylogenetic Placement and Justification:** †*Gasterorhamphosus zuppichini* calibrates the MRCA of *Scomber scombrus* and *Syngnathus fuscus*. †*G. zuppichini* is the oldest identifiable member of *Syngnathiformes* and is placed in the crown of that order based on the results of phylogenetic analyses of morphological characters ^5^. See a full justification of this calibration in ^5^.

**Stratigraphy**: Nardo, Italy; Late Cretaceous, Santonian-Campanian, 83.6 Ma ^5^.

**Fossil tip age**: 83.6 Ma. The node calibration was created such that 97.5% of the probability distribution fell before this date.

†*Mene purydi*

**Phylogenetic Placement and Justification:** †*Mene purydi* is assignable to pan-*Mene* and calibrates the MRCA of *Betta prima* and *Caranx caninus* in our phylogeny. See a full justification of this calibration in ^6,7^.

**Stratigraphy**: Talara Province, northwestern Perú. Thanetian to Ypresian, Eocene ^8^, minimum 56.0 Ma ^2^.

**Fossil tip age**: 56.0 Ma. The node calibration was created such that 97.5% of the probability distribution fell before this date.

†*Chaychanus gonzalezorum*

**Phylogenetic Placement and Justification:** †*Chaychanus gonzalezorum* is assignable to Pan-*Pomacentridae* based on the following morphological characters: presence of a fused pharyngeal jaw; two spines on the anal fin, both of which are associated with the first anal fin pterygiophore; large supraoccipital crest; three supraneurals; spines and serrations on circumorbital bones; serrated posterior margin of preopercle ^9^. †*C. gonzalezorum* calibrates the MRCA of *Chromis enchrysura* and *Gillellus semicinctus* in our node-dating analyses.

**Stratigraphy**: Belisario Domínguez quarry, Tenejapa-Lacandón geological unit, Chiapas, Mexico; Danian, 63.0  ± 1.5 Ma ^9^.

**Fossil tip age**: 63.0 Ma. The node calibration was created such that 97.5% of the probability distribution fell before this date.

†*Epinephelus casottii*

**Phylogenetic Placement and Justification:** †*Epinephelus cassorti* is assignable to Pan-*Epinephelus* based on the strongly serrated posterior margin of the preopercle, which possesses the characteristic spines of *Epinephelus* ^10^ and calibrates the MRCA of *Mycteroperca rubra* and *Epinephelus hexagonatus* in our node-dating analyses.

**Stratigraphy**: Retznei quarry, Lagenidae Zone, Syria; Badenian, Miocene, 16.5-13 Ma ^10,11^

**Fossil tip age**: 13.0 Ma. The node calibration was created such that 97.5% of the probability distribution fell before this date.

†*Sander* sp. ‘Wood Mountain’

**Phylogenetic Placement and Justification:** †*Sander* sp. ‘Wood Mountain’ is assignable to Pan-*Sander* based on the morphology of vertebral centra and the presence of caniniform teeth ^12^ and calibrates the MRCA of *Perca flavescens* and *Sander lucioperca.*

**Stratigraphy**: Yost Farm, Wood Mountain Formation, Saskatchewan, Canada; middle Miocene, 16.3-13.6 Ma ^12^.

**Fossil tip age**: 13.6 Ma. The node calibration was created such that 97.5% of the probability distribution fell before this date.

*Sander* cf. *lucioperca*

**Phylogenetic Placement and Justification:** *Sander* cf. *lucioperca* is assignable to Pan-*Sander lucioperca* based on the indistinguishability of the bones from living representatives of that species ^13^ and calibrates the MRCA of *Sander marinus* and *Sander lucioperca* in our analysis.

**Stratigraphy**: Kerch, Crimaea, Ukraine; earliest Pliocene ^13^, 5.33 Ma ^2^.

**Fossil tip age**: 5.33 Ma. The node calibration was created such that 97.5% of the probability distribution fell before this date.

†*Proeleginops grandeastmanorum*

**Node calibrated:** †*Proeleginops grandeastmanorum* calibrates the node representing the MRCA of all notothenioids (*Percophis brasiliensis* + *Eleginops maclovinus*) in our analysis. Because this species has been placed in varying positions across *Notothenioidei* and resultantly has unclear affinities to living clades ^7,14–17^, our placement of this fossil calibration on the MRCA of all notothenioids is conservative.

**Stratigraphy**: La Mesata Formation, Seymour Island, Antarctica; Priabonian, Eocene, 33.9 Ma ^2,17^. The La Mesata Formation spans the Eocene, but the marine depositional setting appears in the upper portion of the formation ^18^.

**Fossil tip age**: 33.9 Ma. The node calibration was created such that 97.5% of the probability distribution fell before this date.

†*Raususetarches sakurai*

**Node calibrated:** †*Raususetarches sakurai* calibrates the node representing the MRCA of *Neosebastes thetidis* and *Sebastes vulpes*, and is identifiable as a member of Pan-*Setarchinae* based on the presence of grooved lateral line scales ^19^.

**Stratigraphy**: North bank of Ponshunkarikotangawa River, Koshikawa Formation, Kasuga, Rausu, Hokkaido, Japan; Late Miocene ^19^, 5.33 Ma ^2^.

**Fossil tip age**: 5.33 Ma. The node calibration was created such that 97.5% of the probability distribution fell before this date.

†*Enophrys hoplites*

**Node calibrated:** †*Enophrys hoplites* calibrates the MRCA of *Clinocottus analis* and *Taurulus bubalis* in our analysis. The placement of †*E. hoplites* in Pan-*Enophrys* in crown *Psychrolutidae* based on the following combination of features: membrane bones of head granular and bear small, densely spaced osseus tubercles; upper orbital rim placed dorsal to occiput; elongated, straight, and denticulated upper spine of preopercle; all fin rays except principal caudal rays unbranched; lateral line scales appear as enlarged bony plates; pelvic fin with one spine and three segmented rays ^20^*.*

**Stratigraphy**: Agnevo Formation, 1 km north of the mouth of the Agnevo River, Alexadrovsk region, western coast of Sakhalin Island, Russia; Serravallian-Tortonian ^20^, 13.82-7.246 Ma ^2^.

**Fossil tip age**: 11.608 Ma. The node calibration was created such that 97.5% of the probability distribution fell before this date.

*Gasterosteus aculeatus*

**Node calibrated:** *Gasterosteus aculeatus* is assignable to *Gasterosteus aculeatus* based on meristic counts ^21^ and calibrates the MRCA of *Gasterosteus islandicus* and *G. nipponicus* in our analysis.

**Stratigraphy**: Alta Mira Shale, Monterey Formation Palos Verdes, California; 13.3 to 13.0 Ma ^21^.

**Fossil tip age**: 13.0 Ma. The node calibration was created such that 97.5% of the probability distribution fell before this date.

†*Palaeopholis laevis*

**Node calibrated:** †*Palaeopholis laevis* calibrates the node representing the MRCA of *Ptilichthys goodei* and *Pholidae*. This placement of this species as a member of Pan-*Pholidae* follows Nazarkin ^22^, who identified †*Palaeopholis laevis* as the known sister taxon to *Pholidae* (one step crownward of †*Agnevichthys gretchinae* from the same stratigraphic layer and region) based on the following features: hemonephrapophises present on abdominal vertebrae; absence of pleural ribs; five branchiostegal rays; vertebral count increased. Both these equivalent fossil calibrations are united with Pan-*Pholidae* by the presence of shortened caudal fin rays, unequally amphicoelous vertebrae, and a strongly anteriorly displaced first anal fin pterygiophore ^22^. †*Palaeopholis laevis* is not a crown pholid based on the absence of the following features: branching in procurrent caudal fin rays closest to the principal rays; posterior pterygiophores of the dorsal and anal fins lack rays; palatines edentulous ^22^.

**Stratigraphy**: Agnevo Formation, 1 km north of the mouth of the Agnevo River, Alexadrovsk region, western coast of Sakhalin Island, Russia; Serravallian-Tortonian ^22,23^, 13.82-7.246 Ma ^2^.

**Fossil tip age**: 11.608 Ma. The node calibration was created such that 97.5% of the probability distribution fell before this date.

†*Stichaeopsis sakhalinensis*

**Node calibrated:** †*Stichaeopsis sakhalinensis* calibrates the node representing the MRCA of *Stichaeus punctatus* and *Bryozoichthys marjorius* (*Stichaeidae* sensu stricto). The placement of this species within crown *Stichaeidae* as a member of the genus *Stichaeopsis* is based on the presence of a well-developed system of sensory canals ^24^. However, because *Stichaeopsis* is resolved as polyphyletic in this study, we conservatively calibrate the MRCA of *Stichaeidae* with this fossil. Equivalent fossil calibrations include †*Stichaeus brachigrammus* ^24^ and potentially †*Stichaeus matsubarai* ^25^. †*Nivchia makushoki*, also described from Sakhalin Island, shows some similarity to *Cebidichthys*, which was previously considered a stichaeid ^24^ but is now placed sister to *Dictyosoma* and outside the clade containing *Stichaeidae*+*Zoarcidae*.

**Stratigraphy**: Agnevo Formation, Agnevskaja Svita, Alexadrovsk region, Sakhalin Island, Russia; Serravallian-Tortonian ^22–24^, 13.82-7.246 Ma ^2^.

**Fossil tip age**: 11.608 Ma. The node calibration was created such that 97.5% of the probability distribution fell before this date.

†*Lumpenidae* indet.

**Node calibrated:** †*Lumpenidae* indet. (NSM PV 22683) calibrates the node representing the MRCA of *Lumpenidae* and *Cryptacanthodes*. The placement of NSM PV 22683 as an indeterminate lumpenid is based on the following features: dorsal fin spines equally thick; body covered in a dense coat of scales; posterior dorsal pterygiophore lacks a serial ray ^26^. We conservatively calibrate the MRCA of *Lumpenidae* and its sister taxon using this fossil because the incompleteness of NSM PV 22683 means it cannot be confidently assigned to any living lumpenid genus ^26^.

**Stratigraphy**: Bessho Formation, Nagano Prefecture, Honshu, Japan; Early-Middle Miocene, 13-16 Ma ^26^.

**Fossil tip age**: 13.0 Ma. The node calibration was created such that 97.5% of the probability distribution fell before this date.

†*Zaprora koreana*

**Node calibrated:** †*Zaprora koreana* calibrates the node representing the MRCA of *Zaprora*, *Opisthocentridae*, *Ptilichthys goodei*, and *Pholidae*. The Prowfish *Zaprora silenus* is the only living species in the genus, and so this calibration reflects the placement of †*Zaprora koreana* on the prowfish stem lineage, which is supported by the following features: caudal thorax very deep; dorsal fin spinous; anal fin with a short base and three anterior spines; caudal peduncle conspicuous; neural, haemal, and fin spines and rays very tall ^27^.

**Stratigraphy**: Duho Formation, Pohang, South Korea; middle Miocene, 15 Ma ^27^.

**Fossil tip age**: 15.0 Ma. The node calibration was created such that 97.5% of the probability distribution fell before this date.

†*Anarhichadidae* indet.

**Node calibrated:** †*Anarhichadidae* indet. (ZIN 457p) calibrates the node representing the MRCA of *Anarhichadidae* and *Zoarcidae*, reflecting the uncertain placement of this fossil within or outside the anarhichadid crown clade ^28^. The identity of this fossil as an anarhichadid is supported by the presence of the characteristic dental morphology of this family, which includes canines, conical teeth, and molariform teeth ^28^ placed in a short, deep suspensorium. Additional features uniting ZIN 457p with anarhichadids include: frontals fused; parasphenoid with a developed keel; narrow postorbital skull roof ^28^.

**Stratigraphy**: Kurasi Formation, southwestern Sakhalin Island, Russia; Serravallian–Tortonian, Miocene ^28^, 13.82-7.246 Ma ^2^. The Kurasi Formation is coeval with the Agnevo Formation ^29^.

**Fossil tip age**: 11.608 Ma. The node calibration was created such that 97.5% of the probability distribution fell before this date.

†*Plioplarchus whitei*

**Node calibrated:** †*Plioplarchus whitei* calibrates the node representing the MRCA of *Micropterus dolomieu* and *Elassoma okefenokee* in our analysis. †*P. whitei* is member of *Centrarchidae* based on a phylogenetic analysis of morphological characters and meristic counts ^30^ .

**Stratigraphy**: Sentinel Butte Formation, Billings County, North Dakota, USA; latest Eocene to earliest Oligocene, 34.2 Ma ^30^.

**Fossil tip age**: 34.2 Ma. The node calibration was created such that 97.5% of the probability distribution fell before this date.

†*Micropterus* sp. Snake Creek

**Node calibrated:** †*Micropterus* sp. Snake Creek is a member of Pan-*Micropterus* ^30^ and calibrates the node representing the MRCA of *Micropterus dolomieu* and *Lepomis gibbosus* in our analysis.

**Stratigraphy**: Lower Snake Creek Fauna, Sioux County, Nebraska, USA; early Miocene, 16.0 Ma ^30^.

**Fossil tip age**: 16.0 Ma. The node calibration was created such that 97.5% of the probability distribution fell before this date.

†*Siniperca ikikoku*

**Node calibrated:** †*Siniperca ikikoku* is a member of Pan-*Siniperca* ^31^ and calibrates the node representing the MRCA of *Coreoperca herzi* and *Siniperca knerii* in our analysis. †*S. ikikoku* is assignable to *Sinipercidae* and Pan-*Siniperca* based on the following combination of morphological characters: 11 abdominal and 16 caudal vertebrae; 11 spines and 12 soft rays in dorsal fin; 3 spines and 8 to 9 soft rays in anal fin, cycloid scalation; elongated third dorsal fin spine almost reaches length of fourth and fifth spines; preopercle with four spines along ventral margin; and third to fifth neural spines insert deeply between dorsal pterygiophores.

**Stratigraphy**: Diatomite beds from the Chojabaru Formation at Hachiman, Ashibe, Iki Island, Nagasaki Prefecture, Japan; early middle Miocene, 15.3 Ma ^31^.

**Fossil tip age**: 15.3 Ma. The node calibration was created such that 97.5% of the probability distribution fell before this date.

†*Priscacara serrata*

**Node calibrated:** †*Priscacara serrata* is a member of *Moronidae* based on a phylogenetic analysis of morphological characters ^32^ and calibrates the node representing the MRCA of *Dicentrarchus labrax* and *Morone chrysops* in our analysis.

**Stratigraphy**: Green River Formation, including Lake Uinta and Fossil Butte Member, USA; Eocene, between 56.96 +/-0.12 Ma to 50.5 Ma. Absolute dating of the Fossil Butte Member of the Green River Formation, Wyoming, USA gives an age of 51.66 +/-0.09 Ma ^33^.

**Fossil tip age**: 51.66 Ma. The node calibration was created such that 97.5% of the probability distribution fell before this date.

†*Eoplatax papilio*

**Node calibrated:** †*Eoplatax papilio* is a member of Pan-*Ephippidae* ^34^ and calibrates the node representing the MRCA of *Drepane punctata* and *Platax orbicularius* in our analysis*.* The placement of †*E. papilio* in Pan-*Ephippidae* is supported by the presence of teeth on the ectopterygoids and endopterygoids, the presence of supramaxillae, tricuspid teeth, and nonprotrusive premaxillae.

**Stratigraphy**: Monte Bolca, near Verona, Italy: Ypresian, Eocene, 48.96 to 48.5 Ma ^35^.

**Fossil tip age**: 48.5 Ma. The node calibration was created such that 97.5% of the probability distribution fell before this date.

†*Caucasisciaena ignota*

**Node calibrated:** †*Caucasisciaena ignota* is a member of Pan-*Sciaenidae* ^36^ and calibrates the node representing the MRCA of *Dinoperca petersi* and *Nibea squamosa* in our analysis*.* The placement of †*C. ignota* in Pan-*Sciaenidae* is supported by the following morphological characters: moderately cavernous frontals; supramaxillae absent; single branchiostegal ray on posterior ceratohyal; elongated base of soft dorsal fin; two anal fin spines present; and absence of trisegmented endoskeletal elements of median fins ^36^.

**Stratigraphy**: Various units, Caucasus and Crimea; Early Miocene (lower Burdigalian) ^36^, 20.45 to 15.98 Ma ^2^.

**Fossil tip age**: 15.98 Ma.

†*Hypsocephalus atlanticus*

**Node calibrated:** †*Hypsocephalus atlanticus* is a member of *Lutjanidae* ^37,38^ and calibrates the node representing the MRCA of *Pristipomoides typus* and *Lutjanus campechanus* in our analysis. The placement of †*H. atlanticus* in *Lutjanidae* close to *Hoplopagrus* is supported by the following combination of characters: robust, conical dentition on premaxillae, dentaries, and vomer; basioccipital with vertical and transverse posterior facet; exoccipital articular surface for atlas discontinuous across midline; parasphenoid with globular swelling on posteroventral surface; compressed otic region; dorsal surface of lateral ethmoid lateral to anterior end of frontal compact; convex supraethmoid; morphology of the lateral ethmoid facet for palatine ^37^.

**Stratigraphy**: Milton’s Cave, lower member of the Crystal River Formation, Jackson County, Florida, USA; latest Eocene ^37^ 33.9 Ma ^39^.

**Fossil tip age**: 33.9 Ma.

†*Avitoluvarus eocaenicus*

**Node calibrated:** †*Avitoluvarus eocaenicus* is a member of Pan-*Luvarus* ^40^ and calibrates the node representing the MRCA of *Luvarus imperialis* and *Zanclus cornutus* in our analysis. The placement of †*A. eocaenicus* in Pan-*Luvarus* as a member of †*Avitoluvarus* is supported by the following combination of characters: dentition reduced or lost in adults; median pterygial truss around most of body; pterygial truss shallow and not extensively interdigitated; two or fewer spines in dorsal fin; anal spines absent; all soft dorsal and anal fin rays unsegmented; distal end of first anal fin pterygiophore anteriorly elongated; anteriorly displaced anus; fusion of first four hypurals; broad overlap between caudal fin rays and hypural plate; pelvic fin rudimentary with increasing body size; proximal shafts of anal fin pterygiophores situation in first two interhaemal spaces ^40,41^.

**Stratigraphy**: Uylya-Kushlyuk locality, 2 km northeast of Uylya-Kushlyuk village, Turkmenistan; Danata Formation, uppermost Thanetian-lowermost Ypresian (Paleocene to Eocene) ^6,40,42^. Following Ghezelayagh et al. ^6^, we use an age of 55.8 Ma.

**Fossil tip age**: 55.8 Ma.

†*Eoleiognathus dorsalis*

**Node calibrated:** †*Eoleiognathus dorsalis* is a member of Pan-*Leiognathidae* ^43,44^ and calibrates the node representing the MRCA of *Chaetodon unimaculatus* and *Gazza minuta* in our analysis. The placement of †*E. dorsalis* in Pan-*Leiognathidae* is supported by the following combination of characters: small, protractile mouth; high, pointed supraoccipital crest; origin of dorsal fin anteriorly shifted; 24 vertebrae; non-oligomerized caudal skeleton; iliac process well developed; and meristic counts of the dorsal and anal fins ^43^.

**Stratigraphy**: Monte Bolca, near Verona, Italy: Ypresian, Eocene, 48.96 to 48.5 Ma ^35^.

**Fossil tip age**: 48.5 Ma. The node calibration was created such that 97.5% of the probability distribution fell before this date.

†*Chaetodon ficheuri*

**Node calibrated:** †*Chaetodon ficheuri* is a member of Pan-*Chaetodon* ^45^ and calibrates the node representing the MRCA of *Chaetodon unimaculatus* and *Forcipiger longirostris* in our analysis. The placement of †*C. ficheuri* in *Chaetodontidae* as a member of Pan-*Chaetodon* is supported by the following combination of characters: supraoccipital crest tall; frontals sutured to epiotics; second infraorbital bone excluded from orbit; supracleithral lateral line canal incomplete medially; 24 vertebrae; ctenoid scales arranged in ascending rows from the lower anterior to upper posterior; predorsal formula 0/0+2/1/1; pelvic fin spine and basipterygium interlocked via two small flanges at proximal end of pelvic spine fused through pelvic articular foramen; lateral line scales disposed as smooth arch below the dorsal fin and extend almost to last dorsal fin ray; caudal fin skeleton structure and composition ^45^.

**Stratigraphy**: Massive diatomites at Les Planteurs, Raz-el-Aïn, Saint-Denis du Sig, and St. Eugène, Chelif Basin, northwestern Algeria; early Messinian, ~7 to 6 Ma ^45^.

**Fossil tip age**: 6.0 Ma.

†*Tauichthys aspesae*

**Node calibrated:** †*Tauichthys aspesae* is a member of Pan-*Acanthurinae* ^46,47^ and calibrates the node representing the MRCA of *Ctenochaetus striatus* and *Naso annulatus* in our analysis. The placement of †*T. aspesae* in *Acanthuridae* is supported by the following combination of characters: hypurals unconsolidated; three epurals; first dorsal and anal fin spines elongated such that they protrude prominently outward; absence of laterally expanded pterygial shields around first dorsal and anal fin spine bases; five branchiostegal rays ^47^.

**Stratigraphy**: Monte Bolca, near Verona, Italy: Ypresian, Eocene, 48.96 to 48.5 Ma ^35^.

**Fossil tip age**: 48.5 Ma. The node calibration was created such that 97.5% of the probability distribution fell before this date.

†*Sparnodus vulgaris*

**Node calibrated:** †*Sparnodus vulgaris* is a member of Pan-*Sparidae* ^48^ and calibrates the node representing the MRCA of *Lethrinus nebulosus* and *Pagrus major* in our analysis. The placement of †*S. vulgaris* in Pan-*Sparidae* is supported by phylogenetic analyses of morphological characters ^48^.

**Stratigraphy**: Monte Bolca, near Verona, Italy: Ypresian, Eocene, 48.96 to 48.5 Ma ^35^.

**Fossil tip age**: 48.5 Ma. The node calibration was created such that 97.5% of the probability distribution fell before this date.

*Linophryne* cf. *indica*

**Node calibrated**: *Linophryne* cf. *indica* calibrates the node representing the MRCA of *Linophryne* and *Acentrophryne* in our analysis. This is based on the identification of a *Linophryne* cf. *indica* fossil as representative of a member of the *Linophryne* total clade ^49^.

This fossil is placed in *Linophryne* and resembles *L. indica* based on: prominent sphenotic spine, rounded protuberance on frontal, very large teeth and preopercular spine, single pair of teeth on vomer, absence of dentary symphyseal spine, and enlarged pharyngobranchials ^49^.

**Stratigraphy**: Upper portion of the Capistrano Formation, Los Angeles Basin, California, USA: Middle-Upper Miocene, Neogene 14.0 to 5.333 Ma. ^S50^

**Fossil tip age**: 5.333 Ma.

*Chaenophryne* cf. *melanorhabdus*

**Node calibrated**: *Chaenophryne* cf. *melanorhabdus* calibrates the node representing the MRCA of *Chaenophryne* and *Bertella* in our analysis. This is based on the identification of *Chaenophryne* cf. *melanorhabdus* fossils as representative of a member of the *Chaenophryne* total clade.^49^ This fossil is placed in *Chaenophryne* and resembles *C. melanorhabdus* based on: cancellous bone texture, strongly dorsolaterally convex frontal, neurocranium deepens at the level of the parietal, absence of a sphenotic spine and dentary symphyseal spine, large mandibular dentition, blunt articular spine and reduced angular spine, triangular, posteriorly concave opercle, subopercle with small anterior spine and long spine at upper posterior corner, two anteriorly-directed dorsal and ventral processes near fused first preural and ural centra, long illicium runs over 20% of standard length, and posterior margin of cleithrum sigmoid ^49^.

**Stratigraphy**: Upper portion of the Capistrano Formation, Los Angeles Basin, California, USA: Middle-Upper Miocene, Neogene 14.0 to 5.333 Ma ^50^.

**Fossil node age**: 5.333 Ma. The node calibration was created such that 97.5% of the probability distribution fell before this date.

*Oneirodes* sp.

**Node calibrated:** *Oneirodes* sp. calibrates the node representing the MRCA of *Oneirodes* and *Spiniphryne* in our analysis. This is based on the identification of an *Oneirodes* sp. fossil as representative of a member of the *Oneirodes* total clade ^49^. This fossil is placed in *Oneirodes* based on: frontal dorsolaterally convex, sphenotic spine present and extensively developed, mouth extends beyond orbit, jaw is large and depressible, large pharyngobranchial teeth present, and the ventral half of the subopercle is semicircular ^49^.

**Stratigraphy**: Upper portion of the Capistrano Formation, Los Angeles Basin, California, USA: Middle-Upper Miocene, Neogene 14.0 to 5.333 Ma. ^S50^

**Fossil node age**: 5.333 Ma. The node calibration was created such that 97.5% of the probability distribution fell before this date.

†*Histionotophorus bassani*

**Node calibrated:** †*Histionotophorus bassani* calibrates the node representing the MRCA of *Brachionichthys* and *Rhyncherus* in our analysis. A second, equivalent-age fossil calibration is given by †*Orrichthys longimanus* ^51^. These genera are identifiable as members of the clade *Brachionichthyidae* based on the presence of two elongated radials on the pectoral fine, curved ceratobranchials I through IV, five branchiostegal rays, absence of vomerine and palatine teeth, and the combination of a horizontal to oblique mouth, a flat ventral vomer surface, a broad quadrate articular head, two pharyngobranchials, a slightly curved ventral column, absence of epurals, and three cephalic dorsal fin spines ^51^. This placement is supported by phylogenetic analysis of morphological characters ^51^.

**Stratigraphy**: Monte Bolca, near Verona, Italy: early-late Ypresian, Eocene, Paleogene 48.96 to 48.5 Ma ^35^.

**Fossil tip age**: 48.5 Ma. The node calibration was created such that 97.5% of the probability distribution fell before this date.

†*Neilpeartia ceratoi*

**Node calibrated:** †*Neilpeartia ceratoi* ^52^ calibrates the node representing the MRCA of the clade containing species in the genera *Abantennarius, Antennarius*, *Antennatus*, *Fowlerichthys, Histrio*, and *Nudiantennarius* in our analysis. †*Eophryne barbutii* may be an equivalent fossil calibration ^53^, but †*Neilpeartia ceratoi* represents the most likely member of this crown clade among the Monte Bolca forms ^52^. †*Neilpeartia ceratoi* is identified as an antennariid based on: ectopterygoid triradiate, endopterygoid present, epural present, spatulate post-maxillary process of the premaxilla, reduced opercle, dorsoventrally deep body shape, and enlarged third dorsal fin spine ^52^. Phylogenetic analysis of morphological characters placed †*Neilpeartia ceratoi* sister to the genus *Fowlerichthys* ^52^*.*

**Stratigraphy**: Monte Bolca, near Verona, Italy: early-late Ypresian, Eocene, Paleogene 48.96 to 48.5 Ma ^35^.

**Fossil tip age**: 48.5 Ma. The node calibration was created such that 97.5% of the probability distribution fell before this date.

†*Antennarius monodi*

**Node calibrated:** †*Antennarius monodi* calibrates the crown of *Fowlerichthys* (formerly the “*Antennarius*” *ocellatus* group)^54^ .The assignment of †*Antennarius monodi* to *Antennariidae* is based on: triradiate ectopterygoid present, a keel-like posteromedial process of the vomer, a spatulate post-maxillary process of the premaxilla, and a reduced opercle ^54^. Assignment to the “*Antennarius*” *ocellatus* group is based on the presence of close-set bifid dermal spinules covering the skin, bifurcated caudal fine rays, the presence of an epural, and meristic similarities.

**Stratigraphy**: Diatomites of Raz-el-Aïn, near Oran, north-east Algeria: late Miocene, Messinian)^54^, which is 7.246 to 5.333 Ma ^55^.

**Fossil tip age**: 5.333 Ma. The node calibration was created such that 97.5% of the probability distribution fell before this date.

†*Caruso brachysomus*

**Node calibrated:** †*Caruso brachysomus* calibrates the node representing the MRCA of the clade containing *Lophius* and *Sladenia* (=crown *Lophiidae*)^56^. †*Eosladenia caucasica* from the Middle Eocene of the northern Caucasus provides a penecontemporaneous fossil calibration ^56,57^. The assignment of †*Caruso brachysomus* to total clade Lophiidae is based on: absence of a mesethmoid, an autogenous ascending process of the premaxilla, fusion of the ectopterygoid and endopterygoid, prominent anterodorsal process of the subopercle articulates with anteroventral margin of opercle via connective tissue, restriction of the dentition on the fifth ceratobranchial to discrete rows on lateral and medial margins, presence of a cleithral spine, skin unornamented, and eight caudal fin rays. †*Caruso brachysomus* is identified as a crown lophiid based on the presence of a strongly bifurcated opercle, and as sister to *Sladenia* by a narrow interorbital skull roof and a posteriorly directed expanded distal end of the posteriormost dorsal fin pterygiophore. These character combinations are based on a comprehensive phylogenetic analysis of lophiids using morphological data ^56^.

**Stratigraphy**: Monte Bolca, near Verona, Italy: early-late Ypresian, Eocene, Paleogene 48.96 to 48.5 Ma ^35^.

**Fossil tip age**: 48.5 Ma. The node calibration was created such that 97.5% of the probability distribution fell before this date.

†*Prodiodon erinaceus*

**Node calibrated:** †*Prodiodon erinaceus* calibrates the node representing the MRCA of the clade containing *Diodon* and *Canthigaster*. The placement of †*Prodiodon erinaceus* follows the phylogenetic analysis conducted by ^58^, although equivalent calibrations include †*Heptadiodon echinus*, and †*Zignodon fornasieroae* ^58^.

**Stratigraphy**: Monte Bolca, near Verona, Italy: early-late Ypresian, Eocene, Paleogene 48.96 to 48.5 Ma ^35^.

**Fossil tip age**: 48.5 Ma. The node calibration was created such that 97.5% of the probability distribution fell before this date.

†*Moclaybalistes danekrus*

**Node calibrated:** †*Moclaybalistes danekrus* calibrates the node representing the MRCA of the clade containing *Balistes* and *Triacanthus*. The placement of †*Moclaybalistes danekrus* follows the phylogenetic analysis conducted in Close et al. ^58^, although penecontemporaneous calibrations include †*Eospinus daniltshenkoi* (see ^58^).

**Stratigraphy**: Monte Bolca, near Verona, Italy: early-late Ypresian, Eocene, Paleogene 48.96 to 48.5 Ma. ^35^.

**Fossil tip age**:48.5 Ma. The node calibration was created such that 97.5% of the probability distribution fell before this date.

†*Ctenoplectus williamsi*

**Node calibrated:** †*Ctenoplectus williamsi* is a member of Pan-*Triodon* in *Tetraodontoidei* that calibrates the node representing the MRCA of the clade containing *Triodon macropterus* and *Aracana aurita.* The identity of †*Ctenoplectus williamsi* as a member of Pan-*Triodon* is supported by: jaws form acute arrowhead shapes in dorsal and ventral views, dental microstructure consists of simple, round elements densely stacked together, the presence of osteodentinic posterointernal walls and absence of anteroexternal walls in the dental microstructure, and the presence of posttemporals, ribs, pelves, and spines in the dorsal fin ^58^.

**Stratigraphy**: London Clay Formation, England, UK: Ypresian, Eocene, Paleogene 53 Ma ^58^.

**Fossil tip age**: 53 Ma. The node calibration was created such that 97.5% of the probability distribution fell before this date.

†*Synagropoides steparenkorum*

**Node calibrated:** †*Synagropoides steparenkorum* is a member of Pan-*Acropomatidae* ^59^ and calibrates the node representing the MRCA of *Symphysanodon octoactinus* and *Synagrops spinosus* in our analysis. The placement of †*S. steparenkorum* in Pan-*Acropomatidae* is supported by the following combination of characters: two separate dorsal fins; small number of rays in second dorsal fin and anal fin; three anal fin spines; vertebral formula 10+15; non-oligomerized caudal skeleton; two supernumerary spines in second dorsal fin ^59^.

**Stratigraphy**: Gomyi Luch, Krasnodar krai, Kuma Horizon, Turkmenistan; Bartonian Stage of the Eocene ^59^, 41.0 to 37.7 Ma ^2^.

**Fossil tip age**: 37.7 Ma.

†*Astroscopus countermani* (= “*Uranoscopus*” sp.)

**Node calibrated:** *Astroscopus countermani* calibrates the node representing the MRCA of the clade containing *Astroscopus y-graecum* and *Kathetostoma averruncus* (*Uranoscopidae*). The identity of †*A. countermani* as a member of *Astroscopus* is established by the shape of the neurocranium, obliteration of the orbital foramen, and the presence of a large tuberosity on the hyomandibula dorsolateral face ^60^. Equivalent calibrations are cf. *Uranoscopus* sp./ *Uranoscopidae indet.* ^61^.

**Stratigraphy**: St. Marys Formation of Maryland, USA: at latest Tortonian Stage of the Miocene 7.25 Ma ^61^.

**Fossil tip age**: 7.25 Ma. The node calibration was created such that 97.5% of the probability distribution fell before 7.25 Ma.

†*Phyllopharyngodon longipinnis*

**Node calibrated:** †*Phyllopharyngodon longipinnis* calibrates the node representing the MRCA of *Labridae.* A placement in *Labridae* is supported by the presence of a single posteriorly oriented supraneural, developed pharyngeal jaws, and cycloid scales ^62–64^, and a placement on the stem of *Hypsigenyinae* is supported by the presence of phyllodont dentition ^62^. Figure 3 in ^62^ appears to show several additional synapomorphies of *Labridae* present in †*P. longipinnis*: parasphenoid with well-developed adductor process and pharyngeal apophysis, the close association of the nasal and frontal bones, and the coalescence of the articular and ascending processes of the premaxilla ^63^.

**Stratigraphy**: Monte Bolca, near Verona, Italy: early-late Ypresian, Eocene, Paleogene 48.96 to 48.5 Ma ^35^.

**Fossil tip age**: 48.5 Ma. The node calibration was created such that 97.5% of the probability distribution fell before 48.5 Ma.

†*Trigonodon jugleri*

**Node calibrated:** †*Trigonodon jugleri* calibrates the node representing the MRCA of the clade containing *Pseudodax moluccanus*, *Bodianus* spp., *Clepticus* spp., and *Semicossyphus* spp. †*Trigonodon jugleri* is placed in *Hypsigenyinae* as sister to *Pseudodax moluccanus* based on the following characters: single modified giant tooth in the dentary, phyllodont dentition in the upper pharyngeal tooth plates oriented along multiple oblique rows, and nodular dentition border the ovoid margin of the lower pharyngeal jaw ^64,65^.

**Stratigraphy**: Various formations, Austria: Burdigalian (Ottnangian), Early Miocene ^64,65^, 20.43-15.97 Ma ^2^

**Fossil tip age**: 15.97 Ma. The node calibration was created such that 97.5% of the probability distribution fell before 15.97 ma.

†*Bolbometopon* sp.

**Node calibrated:** †*Bolbometopon* sp. calibrates the node representing the MRCA of the clade containing *Bolbometopon muricatum* and *Cetoscarus* spp. †*Bolbometopon* sp. is identifiable to the genus *Bolbometopon* based on characteristic dental morphology, which includes elongated dentition that lacks thick cemented covering and wherein only the distal face of each tooth is visible on the dental surface such that the teeth form a mosaic pattern, and the exposed surface of each tooth appears square with a dorsally-placed nodule ^64,66^.

**Stratigraphy**: East Shore of Dutch Bay, NW Province, Sri Lanka: Late Miocene (?Tortonian-Messinian Stages) 11.6-5.33 Ma ^2^.

**Fossil tip age**: 7.25 Ma [the Tortonian-Messinian boundary]. The node calibration was created such that 97.5% of the probability distribution fell before 7.25 Ma.

†*Calotomus preisli*

**Node calibrated: :** †*Calotomus preisli* calibrates the node representing the MRCA of crown *Calotomus*. †*Calotomus preisli* is identifiable to the *Scarinae* based on the presence of upper pharyngeal plates with 1-3 tooth rows and the presence of a lateral canine ^66^ and placed in *Calotomus* based on the combination of the following features: non-coalesced oral jaw teeth, upper pharyngeal teeth short and gracile, and the presence of a conical tooth on the medial face of the premaxilla next to the symphysis ^66^. The presence of a low premaxillary tooth count and the absence of discrete rows of premaxillary teeth serve to unite †*Calotomus preisli* with *C. spinidens*.

**Stratigraphy**: Leitha Limestone, St. Margarethen, Burgenland, Austria: Upper Badenian, Miocene ^66,67^. The Badenian stretches from 16.30 to 12.83 Ma ^68^.

**Fossil tip age**: 12.83 ma. The node calibration was created such that 97.5% of the probability distribution fell before 12.83 Ma.

†*Wainwrightdabrus agassizi*

**Node calibrated:** †*Wainwrightdabrus agassizi* is an equivalent calibration for the node representing the MRCA of the clade containing *Labrus* and *Tautoga*. *Wainwrightdabrus agassizi* is placed in *Labrinae* based on: serrated posterior margin of the preopercle, unshortened first vertebral neural spine, >25 total vertebrae, >10 dorsal fin spines, a Y-shaped lower pharyngeal jaw, a slightly curved supraneural, and a steeply ventrally deflected posterior lateral line ^67^. Overall skeletal structure, in particular the general morphology of the suspensorium, in †*Wainwrightdabrus agassizi* compares favorably with *Labrus* spp. among *Labrinae*, though †*Wainwrightdabrus* differs from all members of this genus in numerous discrete morphological and meristic characters ^67^.

**Stratigraphy**: Leitha Limestone, St. Margarethen, Burgenland, Austria: Upper Badenian, Miocene ^66,67^. The Badenian stretches from 16.30 to 12.83 Ma ^68^.

**Fossil tip age**: 12.83 Ma.

†*Symphodus westneati*

**Node calibrated:** †*Symphodus westneati* calibrates the node representing the MRCA of the clade containing *Symphodus* and *Tautogalabrus.* †*Symphodus westneati* is assignable to *Labrinae* based on the presence of a posteriorly serrated preopercle, five branchiostegal rays, an unshortened neural spine of the first vertebra, and >25 vertebrae, and to *Symphodus* based on the presence of three anal fin spines, shortened oral jaws, the presence of a single row of teeth, and general skeletal morphology ^67^.

**Stratigraphy**: Leitha Limestone, St. Margarethen, Burgenland, Austria: Upper Badenian, Miocene ^66,67^/ The Badenian stretches from 16.30 to 12.83 Ma ^68^.

**Fossil tip age**: 12.83 Ma. The node calibration was created such that 97.5% of the probability distribution fell before 12.83 Ma.

†*Coris sigismundi*

**Node calibrated:** †*Coris sigismundi* calibrates the node representing the MRCA of the clade *Julidinae.* Because living *Coris* are strongly inferred as paraphyletic in this paper, we cannot assign †*Coris sigismundi* to a particular clade in *Julidinae.* †*Coris sigismundi* is assignable to *Julidinae* based on: presence of a frontal recess, preopercle with entire ventral and posterior margins, shortened first neural spine supported by an autogenous neural arc, heavily reduced fifth hypural, posteroventral spine on urohyal, and abrupt ventral deflection of the lateral line below the fleshy dorsal fin ^67^. †*Coris sigismundi* is united with wrasses traditionally placed in the genus *Coris* by the pattern of oral and pharyngeal dentition, the presence of six branchiostegal rays, the total vertebral count of 25, formulae of the median and paired fins, enlarged haemal arc and spine of the first caudal vertebra, and the vertical development of the spatulate prezygapophyses of the abdominal vertebrae ^67^.

**Stratigraphy**: Leitha Limestone, St. Margarethen, Burgenland, Austria: Upper Badenian, Miocene ^66,67^. The Badenian stretches from 16.30 to 12.83 Ma ^68^.

**Fossil tip age**: 12.83 ma. The node calibration was created such that 97.5% of the probability distribution fell before 12.83 Ma.

**Supplementary Results and Discussion.**

***Phylogenetic Results.–***Our phylogenomic analyses inferred relationships among *Eupercaria* that are very similar to those recovered previously using ultraconserved elements ^6,69–72^ and exons ^73–79^(Figures S1-S20). In the 75% taxon-locus occupancy matrix, which included 995 UCE loci, the total number of sites was 263,535 (average of 264.86 per locus), the total number of parsimony-informative sites (PIS) was 137,947 (average of 138.64 per locus, comprising 52% of all sites), and the nucleotide base composition was 27.9% A, 22.2% C, 22.0% G, and 27.8% T. The total lengths of gene trees ranged from 0.27 to 20.88, with a mean length of 5.15 and a median length of 4.72. Comparisons of different metrics of node support and the full taxon-locus occupancy matrix are in Figures S21 and S22, respectively.

We infer that *Perciformes* sensu Near and Thacker^7^ is the first order-level clade to diverge in *Eupercaria* (Figure S1). Earlier analyses that used small sets of nuclear markers and mitochondrial DNA occasionally inferred that *Perciformes* was placed within a clade formed by the other taxonomic orders in *Eupercaria* ^80–82^, but this hypothesis can largely be rejected by the consilience observed across studies of much larger sets of loci ^6,70,73,76^. A remaining issue is the position of mojarras, which comprise *Gerreidae*. This is a rogue taxon that has shifted positions across phylogenetic analyses that inferred evolutionary relationships using both small sequence sets ^81–85^ and genome-wide marker data ^6,76,78^. Alternatively concatenating loci or estimating a species tree topology modifies the placement of *Gerreidae* in phylogenetic analyses of UCEs ^72^. In the expanded *Eupercaria* phylogeny that we present here, *Gerreidae* is inferred to be the sister lineage of *Acanthuriformes* when loci are concatenated. Thus, although some uncertainty remains regarding ingroup ordinal relationships among *Eupercaria*, the deepest divergences in the lineage and the monophyly of all order-level clades appears to be stable (Figure S1).

Below the level of taxonomic orders, the monophyly of most taxonomic families of *Eupercaria* as delimited in the phylogenetic rank-free classification ^7^ are supported in our expanded phylogeny. Our phylogeny resolves the monophyly of taxonomic families across all five taxonomic orders in *Eupercaria* in a manner entirely consistent with the phylogenetic rank-free taxonomies proposed in previous studies ^7,86^.

***Perciformes.*–**In previous systematic treatments, incomplete species sampling has been considered a roadblock to a robust taxonomy, particularly in *Perciformes* ^87^. Owing to the increased taxonomic sample of genera and species in our phylogeny, including species and genera traditionally included in *Serranidae* (now widely recognized as paraphyletic) ^6,7,76,87–90^, our phylogeny clarifies the identity and composition of major clades in this taxonomic order. The following lineages of *Perciformes* are consistently recovered as monophyletic and comprise the major bins of species diversity in this taxonomic order: *Epinephelidae* (groupers, soapfishes, and painted basslets), *Anthiadidae* (basslets), *Serranidae* (hamlets and relatives), *Acanthistius* (wirrahs), *Bembropidae* (duckbills), *Niphon spinosus* (Sawedged Perch), *Trachinidae* (weevers), *Percidae* (perches, walleyes, and darters), *Notothenioidei* (Antarctic icefishes and relatives), *Platycephalidae* (flatheads), *Scorpaenoidea* (rockfishes, stonefishes, ghost flatheads, and Mote sculpin), *Bembridae* (deepwater flatheads), *Triglidae* (sea robins), *Anoplopomatidae* (sablefishes), *Cottoidea* (sculpins, snailfishes, and greenlings), *Gasterosteidae* (sticklebacks), and *Zoarcoidea* (eelpouts, wolffishes, and pricklebacks). The relationships among these lineages are inconsistently recovered across different analyses of genome-wide marker data ^6,76,78,87^. However, *Niphon spinosus*, *Percidae*, and *Trachinidae* form a clade and *Epinephelidae* and *Anthiadidae* are sister lineages, as in previous analyses ^6,76,87^. Because we sample multiple species of *Acanthistius* and infer that this genus represents its own long branch that is not clearly placed in any taxonomic family of *Perciformes*, our analyses ameliorate concerns about the placement of this lineage in studies with smaller species samples ^87^.

Several notable results in *Perciformes* deserve mention (Figure S2-S4). First, we find that the genera *Anthias*, *Cephalopholis*, *Centropristis*, *Epinephelus*, *Pseudanthias*, and *Serranus* are broadly para- and polyphyletic, as is traditional *Serranidae*, which comprises the clades *Acanthistius, Anthiadidae, Grammistinae, Epinephelinae*, *Liopropomatinae*, and *Serraninae*. The resolution of these clades matches that in Ghezelayagh et al. ^6^: *Anthiadidae* and *Epinephelidae* are sister taxa, *Liopropomatinae* is a grade leading to *Grammistinae*, and *Serranidae* sensu stricto is sister to *Bembropidae* (Figures S2-S3).

As in previous analyses, *Epinephelus* contains species in the genera *Anyperodon* and *Cromileptes* ^91,92^ *Paranthias* is contained within *Cephalopholis* ^89^, and *Gracila albomarginata* and *Cephalopholis igarashiensis* form a clade exclusive of other *Cephalopholis* ^89^. Intriguingly, *Hyporthodus* is placed deep within *Cephalopholis*, which may be the result of introgressive hybridization known to occur among species of the former genus with other groupers ^93^. *H. nigritus*, the representative of *Hyporthodus* in our analyses, is also one of the most deeply divergent species in the genus in previous phylogenetic analyses focusing on that genus ^94^, and so it might simply be that the limited earlier analyses of this species did not adequately sample other grouper lineages such that its placement in *Cephalopholis* would be recovered. In *Anthiadidae*, we resolve three major lineages as in Tang and Chen ^90^, although unlike that study *Hypoplectrodes* is placed sister to their clade III (*Odontanthias* and allies) + clade I (*Caesioperca* and allies). As in Tang and Chen ^90^, species in *Paranthias* are distributed across these clades. The lack of species overlap between our study and Tang and Chen ^90^ is illustrative of the need for a comprehensive phylogenomic analysis of anthiadids to resolve their systematics. In *Serranidae*, species assigned to *Serranus* are nested in four different clades: *S. tigrinus* and *S. baldwini* are sister taxa, *Serranus cabrilla* Linneaus 1758, the type species, is sister to a clade containing species in the genera *Bullisichthys, Paralabrax*, and *Schultzea*, *Serranus notospilus* is sister to *Centropristis fuscula*, and *Serranus luciopercanus* is sister to species in the genus *Hypoplectrus* (Figure S3).

Deeper into the perciform tree, we also fail to recover monophyly of North American and Asian walleyes and relatives in the genus *Sander* (Figure S3). Instead, North American species (classically placed in the genus *Stizostedion*) form the sister lineage to Asian *Sander* and the clade containing *Romanichthys* and *Zingel.* The paraphyly of *Sander* is strongly supported and deserves additional study, especially since an upcoming study of ultraconserved elements that largely overlaps in taxon sampling with this one (Wood et al., in review) finds *Sander* to be monophyletic. Secondly, we infer that the neutrally buoyant notothenioid genera *Aethotaxis* *Dissostichus,* and *Pleuragramma* do not form a clade (Figure S4; contra ^95^). This implies that neutral buoyancy has evolved at least twice within *Notothenioidei*. We also highlight our high taxon sampling for *Notothenioidei* (71% of genera, 30% of species), which gives us an additional degree of confidence in rejecting neutrally buoyant notothenioid monophyly.

***Centrarchiformes.*–**Our phylogeny of *Centrarchiformes* is congruent with previous studies ^6^ except in the placement of *Oplegnathus* as sister to the clade formed by *Kyphosidae*, *Kuhlia*, and *Terapontidae* rather than the clade formed *Dichistius* and *Caesioscorpis*. Of note is that all genera of the temperate perches in *Percichthyidae* are reciprocally monophyletic, a result inferred thanks to increased species-level sampling.

***Labriformes.*–**Our phylogeny of *Labriformes* is congruent with previous studies ^6,72^ and no major new phylogenetic relationships are inferred by us. The relationships of the major lineages in *Labridae* and *Uranoscopoidei* should be considered stable.

***Acropomatiformes.*–**Although we resolve two major lineages in *Acropomatiformes* (clade I consisting of *Acropomatidae*, *Ostracoberyx*, *Scombrops*, *Symphysanodon*, *Howellidae*, and *Epigonidae*; clade II consisting of *Polyprionidae*, *Pempheridae*, *Stereolepis, Banjos*, *Pempheridae*, *Glaucosoma*, *Bathyclupeidae*, *Creedidae*, *Pentacerotidae*, *Hemerocoetidae*, and allies) as in previous studies ^6,70^, our expanded taxon sampling allows us to infer the relationships of a handful of morphologically abberant species with previously contested affinities. For example, we infer that Grape-Eyed Seabass *Hemilutjanus macropthalmos* and Longfin Pike *Dinolestes lewini* are members of a weakly supported lineage that also includes giant sea basses (*Stereolepis*) and armourheads (*Pentacerotidae*). We also infer that *Synagropidae* is sister to *Bathyclupeidae*. Instead of being sister to *Acropomatidae* ^6^, *Ostracoberyx* is sister to all other lineages in clade I.

***Acanthuriformes.*–**Although we infer a very similar phylogeny of *Acanthuriformes* to previous studies using ultraconserved elements ^6^, our phylogeny differs from those published previously in several key respects. First, angelfishes (*Pomacanthidae*) are placed sister to all other acanthuroids, rather than sister to butterflyfishes (*Chaetodontidae*) and ponyfishes (*Leiognathidae*). *Callanthiadidae* is placed sister to the clade containing *Siganus*, *Tetraodontoidei*, *Lophioidei*, *Priacanthidae*, *Scatophagidae*, and allies, instead of sister to *Lethrinidae* and *Sparidae* ^6^.

***Divergence Times.*–**The divergence times of major lineages in *Eupercaria* that we estimate are broadly congruent with those estimated in earlier studies. However, some estimated ages, such as the crown age of *Perciformes* at 51.06 Ma (95% HPD: 40.63, 70.73 Ma), are younger than recent analyses with dense taxonomic sampling.^6,75^ The time-calibrated phylogeny of *Eupercaria* that we present is also incompatible with the interpretation of several Cretaceous fishes as putative tetraodontoids,^96^ which has been controversial.^6,8^ We note that, as of now, the placement of these fossils in *Tetraodontoidei* has not truly been tested in phylogenetic analyses. Although these fossils have been included in phylogenetic analyses of morphological character data, these taxon-character matrices either exclusively include tetraodontoids^96^ or one non-tetraodontoid^97^ species. The same practices have led to the incorrect identification of fossil lizards as birds^98^ and Triassic reptiles of unclear affinity as members of the anguimorph squamate crown clade.^99^ Although a proper analysis of the affinities of these Cretaceous fish fossils remains to be performed, their affinities to a deeply-nested lineage in *Eupercaria* should be viewed with proper skepticism given the lack of supporting evidence from both the fossil record^8^ and time-calibrated molecular phylogenies.^6,70,73,76,81^

The earliest representative crown perciform fossils are ‘scorpaenioids’ of uncertain affinity from the Eocene of Monte Bolca, Italy,^100^ which is congruent with the Ypresian age for the rapid diversification of perciform lineages in our time-calibrated phylogeny (Figure 1). Given that we infer that genera traditionally placed in *Serranidae* are a paraphyletic grade of early-diverging perciforms (Figure 1, Supplementary Information), putative serranids known from the Paleocene of Mexico ^101^ may instead represent stem-lineage perciforms. Even though we use nearly all of the same fossil calibrations as previous studies that have estimated mid- to early Late Cretaceous divergence times for families of anglerfishes and pufferfishes in *Acanthuriformes* ^69,79^, our divergence time estimates for these lineages in the new time-calibrated phylogeny indicate they are latest Cretaceous to Paleocene in age. The older divergence times estimated in previous studies may be attributable to the use of wide priors for the age of the root node to account for the missing fossil record of order-level clades in *Eupercaria* and outgroups ^8,102,103^.

**References.**

1.	Murray, A. Mid-Cretaceous acanthomorph fishes with the description of a new species from the Turonian of Lac des Bois, Northwest Territories, Canada. *Vertebrate Anatomy Morphology Palaeontology* **1**, 101–115 (2016).

2.	Gradstein, F. M., Ogg, J. G., Schmitz, M. & Ogg, G. *The Geologic Time Scale 2020*. (Elsevier Science, Amsterdam, The Netherlands, 2021).

3.	Andrews, J. V., Schein, J. P. & Friedman, M. An earliest Paleocene squirrelfish (Teleostei: Beryciformes: Holocentroidea) and its bearing on the timescale of holocentroid evolution. *Journal of Systematic Palaeontology* **21**, 2168571 (2023).

4.	Brownstein, C. D., Dornburg, A. & Near, T. J. Cenozoic evolutionary history obscures the Mesozoic origins of acanthopterygian fishes. *Evolution* qpaf040 (2025) doi:10.1093/evolut/qpaf040.

5.	Brownstein, C. D. Syngnathoid Evolutionary History and the Conundrum of Fossil Misplacement. *Integr Org Biol* **5**, obad011 (2023).

6.	Ghezelayagh, A. *et al.* Prolonged morphological expansion of spiny-rayed fishes following the end-Cretaceous. *Nature Ecology & Evolution* **6**, 1211–1220 (2022).

7.	Near, T. J. & Thacker, C. E. Phylogenetic classification of living and fossil ray-finned fishes (Actinopterygii). *Bulletin of the Peabody Museum of Natural History* **65**, 3–302 (2024).

8.	Friedman, M., V. Andrews, J., Saad, H. & El-Sayed, S. The Cretaceous–Paleogene transition in spiny-rayed fishes: surveying “Patterson’s Gap” in the acanthomorph skeletal record André Dumont medalist lecture 2018. *Geol. Belg.* https://doi.org/10.20341/gb.2023.002 (2023) doi:10.20341/gb.2023.002.

9.	Cantalice, K. M., Alvarado-Ortega, J. & Bellwood, D. R. †*Chaychanus gonzalezorum* gen. et sp. nov.: A damselfish fossil (Percomorphaceae; Pomacentridae), from the Early Paleocene outcrop of Chiapas, Southeastern Mexico. *Journal of South American Earth Sciences* **98**, 102322 (2020).

10.	Ma, K. Y., Craig, M. T., Choat, J. H. & van Herwerden, L. The historical biogeography of groupers: Clade diversification patterns and processes. *Molecular Phylogenetics and Evolution* **100**, 21–30 (2016).

11.	Schultz, O. Ein Zackenbarsch (Epinephelus, Serranidae, Pisces) aus dem Mittel-Miozän von Retznei, Steiermark. *Joannea Geologie und Paläontologie* **2**, (2000).

12.	Murray, A. & Divay, J. First evidence of percids (Teleostei: Perciformes) in the Miocene of North America. *Canadian Journal of Earth Sciences* **48**, 1419–1424 (2011).

13.	Kovalchuk, O. M. & Murray, A. M. Late Miocene and Pliocene Pikeperches (Teleostei, Percidae) of Southeastern Europe. *vrpa* **36**, (2016).

14.	Near, T. J. Estimating divergence times of notothenioid fishes using a fossil-calibrated molecular clock. *Antarctic Science* **16**, 37–44 (2004).

15.	Dornburg, A., Federman, S., Lamb, A. D., Jones, C. D. & Near, T. J. Cradles and museums of Antarctic teleost biodiversity. *Nat Ecol Evol* **1**, 1379–1384 (2017).

16.	Hotaling, S., Borowiec, M. L., Lins, L. S. F., Desvignes, T. & Kelley, J. L. The biogeographic history of eelpouts and related fishes: Linking phylogeny, environmental change, and patterns of dispersal in a globally distributed fish group. *Molecular Phylogenetics and Evolution* **162**, 107211 (2021).

17.	Balushkin, A. Protoeleginops grandeastmanorum Gen. et sp. nov. (Perciformes, Notothenioidei, Eleginopsidae) from the Late Eocene of Seymour Island (Antarctica) is a fossil Notothenioid, not a gadiform. *Journal of Ichthyology* **34**, 10–23 (1994).

18.	Marenssi, S. A., Net, L. I. & Santillana, S. N. Provenance, environmental and paleogeographic controls on sandstone composition in an incised-valley system: the Eocene La Meseta Formation, Seymour Island, Antarctica. *Sedimentary Geology* **150**, 301–321 (2002).

19.	Yabumoto, Y. & Nazarkin, M. V. A New Miocene Scorpaenoid Fish, Raususetarches sakurai gen. et sp. nov. (Teleostei: Scorpaeniformes) from Rausu, Hokkaido, Japan. *jpal* **25**, 93–104 (2021).

20.	Nazarkin, M. V. A new horned sculpin (Pisces: Cottidae) from the Miocene of Sakhalin Island, Russia. *Paleontol. J.* **51**, 77–86 (2017).

21.	Bell, M. A., Stewart, J. D. & Park, P. J. The World’s Oldest Fossil Threespine Stickleback Fish. *cope* **2009**, 256–265 (2009).

22.	Nazarkin, M. Gunnels (Perciformes, Pholidae) from the Miocene of Sakhalin Island. *Journal of Ichthyology* **42**, 279–288 (2002).

23.	Nazarkin, M., Carnevale, G. & Bannikov, A. A New Greenling (Teleostei, Cottoidei) from the Miocene of Sakhalin Island, Russia. *Journal of Vertebrate Paleontology* **33**, 794–803 (2013).

24.	Nazarkin, M. New Stichaeid Fishes (Stichaeidae, Perciformes) from Miocene of Sakhalin. *Journal of Ichthyology* **38**, 279–291 (1998).

25.	Yabumoto, Y. & Uyeno, T. Late Mesozoic and Cenozoic fish faunas of Japan. *Island Arc* **3**, 255–269 (1994).

26.	Nazarkin, M. & Yabumoto, Y. New fossils of Neogene pricklebacks (Actinopterygii: Stichaeidae) from East Asia. *Zoosystematica Rossica* **24**, 128–137 (2015).

27.	Nam, K.-S. & Nazarkin, M. Fossil prowfish, Zaprora koreana , sp. nov. (Pisces, Zaproridae), from the Neogene of South Korea. *Journal of Vertebrate Paleontology* **38**, 1–5 (2018).

28.	Nazarkin, M. V. & Platonov, V. V. Fossil Wolffish (Anarhichadidae) From the Miocene Deposits of Sakhalin Island. *J. Ichthyol.* **60**, 109–113 (2020).

29.	Nazarkin, M. V. The Structure of the Miocene Northwestern Pacific Ichthyofauna as Revealed By Two Fossil Fish Assemblages From Sakhalin Island, Russia. *jpal* **25**, 366–374 (2021).

30.	Near, T. J. & Kim, D. Phylogeny and time scale of diversification in the fossil-rich sunfishes and black basses (Teleostei: Percomorpha: Centrarchidae). *Molecular Phylogenetics and Evolution* **161**, 107156 (2021).

31.	Yabumoto, Y. Siniperca ikikoku, a New Species of Freshwater Percoid Fish from the Miocene of Iki Island, Nagasaki, Japan. *jpal* **24**, 226–237 (2020).

32.	Whitlock, J. A. Phylogenetic Relationships of the Eocene Percomorph Fishes †priscacara and †mioplosus. *Journal of Vertebrate Paleontology* **30**, 1037–1048 (2010).

33.	Smith, M. E., Carroll, A. R. & Singer, B. S. Synoptic reconstruction of a major ancient lake system: Eocene Green River Formation, western United States. *GSA Bulletin* **120**, 54–84 (2008).

34.	Cavalluzzi, M. R. Osteology, phylogeny, and biogeography of the marine fish family Ephippidae (Perciformes, Acanthuroidei), with comments on sister group relationships. https://scholarworks.wm.edu/handle/internal/3849 (2000).

35.	Friedman, M. & Carnevale, G. The Bolca Lagerstätten: shallow marine life in the Eocene. *Journal of the Geological Society* **175**, 569–579 (2018).

36.	Bannikov, A. F., Carnevale, G. & Landini, W. A new Early Miocene genus of the family Sciaenidae (Teleostei, Perciformes) from the eastern Paratethys. *Comptes Rendus Palevol* **8**, 535–544 (2009).

37.	Swift, C. C., Swift, C. C. & Ellwood, B. Hypsocephalus atlanticus, a new genus and species of Lutjanid fish from marine Eocene limestones of northern Florida. *Contributions in science* **230**, 1--29 (1972).

38.	Frédérich, B. & Santini, F. Macroevolutionary analysis of the tempo of diversification in snappers and fusiliers (Percomorpha: Lutjanidae). *Belgian Journal of Zoology* **147**, (2017).

39.	Rincon-Sandoval, M. *et al.* Evolutionary determinism and convergence associated with water-column transitions in marine fishes. *Proceedings of the National Academy of Sciences* **117**, 33396–33403 (2020).

40.	Bannikov, A. & Tyler, J. A new species of the luvarid fish genus †Avitoluvarus (Acanthuroidei, Perciformes) from the Eocene of the Caucasus in Southwest Russia. *Proceedings of the Biological Society of Washington* **114**, 579–588 (2001).

41.	Bannikov, A. F. & Tyler, J. C. *Phylogenetic Revision of the Fish Families Luvaridae and †Kushlukiidae (Acanthuroidei), with a New Genus and Two New Species of Eocene Luvarids*. (Smithsonian Institution, 1995).

42.	Prokofiev, A. M. A remarkable new genus of Carangidae fron1 the Upper Paleocene of Turkmenistan (Osteichthyes: Perciformes). https://www.zin.ru/Journals/zsr/content/2002/zr_2002_11_1_Prokofiev_2.pdf (2002).

43.	Bannikov, A. The new genus Eoleiognathus for the percoid fish Pygaeus dorsalis Agassiz from the Eocene of Bolca in northern Italy, a putative ponyfish (Perciformes, Leoignathidae). *Studi e ricerche sui giacimenti terziari di Bolca* **15**, (2014).

44.	Kovalchuk, O. M., Świdnicka, E. & Stefaniak, K. Early Miocene Ponyfishes (Acanthuriformes, Leiognathidae) of the Carpathian Basin. *Paleontol. J.* **55**, 421–428 (2021).

45.	Carnevale, G. Morphology and biology of the Miocene butterflyfish Chaetodon ficheuri (Teleostei: Chaetodontidae). *Zoological Journal of the Linnean Society* **146**, 251–267 (2006).

46.	Siqueira, A. C., Bellwood, D. R. & Cowman, P. F. Historical biogeography of herbivorous coral reef fishes: The formation of an Atlantic fauna. *Journal of Biogeography* **46**, 1611–1624 (2019).

47.	Tyler, J. & Bannikov, A. A new species of the surgeon fish genus Tauichthys from the Eocene of Monte Bolca, Italy (Perciformes, Acanthuridae). *Bollettino del Museo Civico di Storia Naturale di Verona Geologia Paleontologia Preistoria* **24**, 29–36 (2000).

48.	Day, J. Evolutionary relationships of the Sparidae (Teleostei: Percoidei): integrating fossil and Recent data. *Earth and Environmental Science Transactions of the Royal Society of Edinburgh* **93**, 333–353 (2002).

49.	Carnevale, G., Pietsch, T. W., Takeuchi, G. T. & Huddleston, R. W. Fossil ceratioid anglerfishes (Teleostei: Lophiiformes) from the Miocene of the Los Angeles Basin, California. *Journal of Paleontology* **82**, 996–1008 (2008).

50.	Critelli, S., Rumelhart, P. E. & Ingersoll, R. V. Petrofacies and provenance of the Puente Formation (middle to upper Miocene), Los Angeles Basin, Southern California; implications for rapid uplift and accumulation rates. *Journal of Sedimentary Research* **65**, 656–667 (1995).

51.	Carnevale, G. & Pietsch, T. W. Eocene handfishes from Monte Bolca, with description of a new genus and species, and a phylogeny of the family Brachionichthyidae (Teleostei: Lophiiformes). *Zoological Journal of the Linnean Society* **160**, 621–647 (2010).

52.	Carnevale, G., Pietsch, T. W., Bonde, N., Leal, M. E. C. & Marramà, G. †Neilpeartia ceratoi, gen. et sp. nov., a new frogfish from the Eocene of Bolca, Italy. *Journal of Vertebrate Paleontology* **40**, e1778711 (2020).

53.	Carnevale, G. & Pietsch, T. W. An Eocene Frogfish from Monte Bolca, Italy: The Earliest Known Skeletal Record for the Family. *Palaeontology* **52**, 745–752 (2009).

54.	Carnevale, G. & Pietsch, T. W. Filling the gap: a fossil frogfish, genus Antennarius (Teleostei, Lophiiformes, Antennariidae), from the Miocene of Algeria. *Journal of Zoology* **270**, 448–457 (2006).

55.	Ogg, J. G., Ogg, G. M. & Gradstein, F. M. 15 - Neogene. in *A Concise Geologic Time Scale* (eds Ogg, J. G., Ogg, G. M. & Gradstein, F. M.) 203–210 (Elsevier, 2016). doi:10.1016/B978-0-444-59467-9.00015-7.

56.	Carnevale, G. & Pietsch, T. W. †Caruso, a new genus of anglerfishes from the Eocene of Monte Bolca, Italy, with a comparative osteology and phylogeny of the teleost family Lophiidae. *Journal of Systematic Palaeontology* **10**, 47–72 (2012).

57.	Bannikov, A. The first discovery of an anglerfish (Teleostei, Lophiidae) in the Eocene of the Northern Caucasus. *Paleonotological Journal* **38**, 420–425 (2004).

58.	Close, R. A., Johanson, Z., Tyler, J. C., Harrington, R. C. & Friedman, M. Mosaicism in a new Eocene pufferfish highlights rapid morphological innovation near the origin of crown tetraodontiforms. *Palaeontology* **59**, 499–514 (2016).

59.	Bannikov, A. A new Middle-Eocene marine percoid (Perciformes, Percoidei) from the Northern Caucasus. *Journal of Ichthyology* **42**, 695–700 (2002).

60.	Carnevale, G., Godfrey, S. J. & Pietsch, T. W. Stargazer (Teleostei, Uranoscopidae) cranial remains from the Miocene Calvert Cliffs, Maryland, U.S.A. (St. Marys Formation, Chesapeake Group). *Journal of Vertebrate Paleontology* **31**, 1200–1209 (2011).

61.	Carnevale, G. & Godfrey, S. J. Miocene bony fishes of the Calvert, Choptank, St. Marys and Eastover Formations, Chesapeake Group, Maryland and Virginia. *Smithsonian Contributions to Paleobiology* **100**, (2018).

62.	Bellwood, D. A new fossil fish Phyllopharyngodon longipinnis gen. et sp. nov. (family Labridae) from the Eocene, Monte Bolca, Italy. *Studi e Ricerche sui Giacimenti Terziari di Bolca* **6**, 149–160 (1990).

63.	Bannikov, A. F. & Carnevale, G. Bellwoodilabrus landinii n. gen., n. sp., a new genus and species of labrid fish (Teleostei, Perciformes) from the Eocene of Monte Bolca. *geod* **32**, 201–220 (2010).

64.	Bellwood, D. R., Schultz, O., Siqueira, A. C. & Cowman, P. F. A review of the fossil record of the Labridae. *Annalen des Naturhistorischen Museums in Wien. Serie A für Mineralogie und Petrographie, Geologie und Paläontologie, Anthropologie und Prähistorie* **121**, 125–194 (2019).

65.	Schultz, O. & Bellwood, D. Trigonodon oweni and Asima jugleri are different parts of the same species Trigonodon jugleri, a Chiseltooth Wrasse from the Lower and Middle Miocene in Central Europe (Osteichthyes, Labridae, Trigonodontinae). *Ann. Naturhist. Mus. Wien* **105**, (2004).

66.	Bellwood, D. & Schultz, O. A Review of the Fossil Record of the Parrotfishes (Labroidei: Scaridae) with a Description of a New Calotomus Species from the Middle Miocene (Badenian) of Austria. *Annalen des Naturhistorischen Museums in Wien* **92**, (1991).

67.	Carnevale, G. Middle Miocene wrasses (Teleostei, Labridae) from St.Margarethen (Burgenland, Austria) [121-159. *Palaeontographica Abteilung A* 124–160 (2015) doi:10.1127/pala/304/2015/124.

68.	Hohenegger, J., Ćorić, S. & Wagreich, M. Timing of the Middle Miocene Badenian Stage of the Central Paratethys. *Geologica Carpathica* **65**, 55–66 (2014).

69.	Brownstein, C. D. *et al.* Synergistic innovations enabled the radiation of anglerfishes in the deep open ocean. *Current Biology* **34**, 2541-2550.e4 (2024).

70.	Alfaro, M. E. *et al.* Explosive diversification of marine fishes at the Cretaceous–Palaeogene boundary. *Nature Ecology & Evolution* **2**, 688–696 (2018).

71.	Brownstein, C. D., Harrington, R. C., Radchenko, O. & Near, T. J. The many origins of extremophile fishes. *Proceedings of the Royal Society B: Biological Sciences* **292**, 20250217 (2025).

72.	Brownstein, C. D. *et al.* Phylogenomics establishes an Early Miocene reconstruction of reef vertebrate diversity. *Science Advances* **11**, eadu6149 (2025).

73.	Musilova, Z. *et al.* Vision using multiple distinct rod opsins in deep-sea fishes. *Science* **364**, 588–592 (2019).

74.	Matschiner, M., Böhne, A., Ronco, F. & Salzburger, W. The genomic timeline of cichlid fish diversification across continents. *Nat Commun* **11**, 5895 (2020).

75.	Melendez-Vazquez, F. *et al.* Ecological interactions and genomic innovation fueled the evolution of ray-finned fish endothermy. *Science Advances* **11**, eads8488 (2025).

76.	Hughes, L. C. *et al.* Comprehensive phylogeny of ray-finned fishes (Actinopterygii) based on transcriptomic and genomic data. *Proceedings of the National Academy of Sciences of the United States of America* **115**, 6249–6254 (2018).

77.	Hughes, L. C., Nash, C. M., White, W. T. & Westneat, M. W. Concordance and Discordance in the Phylogenomics of the Wrasses and Parrotfishes (Teleostei: Labridae). *Systematic Biology* **72**, 530–543 (2023).

78.	Hughes, L. C. *et al.* Exon probe sets and bioinformatics pipelines for all levels of fish phylogenomics. *Mol Ecol Resour* **21**, 816–833 (2021).

79.	Miller, E. C. *et al.* Reduced evolutionary constraint accompanies ongoing radiation in deep-sea anglerfishes. *Nat Ecol Evol* **9**, 474–490 (2025).

80.	Near, T. J. *et al.* Resolution of ray-finned fish phylogeny and timing of diversification. *Proceedings of the National Academy of Sciences* **109**, 13698–13703 (2012).

81.	Near, T. J. *et al.* Phylogeny and tempo of diversification in the superradiation of spiny-rayed fishes. *Proceedings of the National Academy of Sciences* **110**, 12738–12743 (2013).

82.	Wainwright, P. C. *et al.* The Evolution of Pharyngognathy: A Phylogenetic and Functional Appraisal of the Pharyngeal Jaw Key Innovation in Labroid Fishes and Beyond. *Systematic Biology* **61**, 1001–1027 (2012).

83.	Betancur-R, R. *et al.* The Tree of Life and a New Classification of Bony Fishes. *PLOS Currents Tree of Life* https://doi.org/10.1371/currents.tol.53ba26640df0ccaee75bb165c8c26288 (2013) doi:10.1371/currents.tol.53ba26640df0ccaee75bb165c8c26288.

84.	Betancur-R, R. *et al.* Phylogenetic classification of bony fishes. *BMC Evolutionary Biology* **17**, 162 (2017).

85.	Rabosky, D. L. *et al.* An inverse latitudinal gradient in speciation rate for marine fishes. *Nature* **559**, 392–395 (2018).

86.	Near, T. J., Brownstein, C. D., Thacker, C. E. & Wainwright, P. C. Phylogenetic Taxonomy of Wrasses and Parrotfishes (Labridae). *Bulletin of the Peabody Museum of Natural History* **66**, 263–338 (2025).

87.	Santos, E. C. *et al.* Ecological axes of skull diversification in a massive vertebrate radiation. 2026.06.19.733456 Preprint at https://doi.org/10.64898/2026.06.19.733456 (2026).

88.	Pondella II, D. J., Craig, M. T. & Franck, J. P. C. The phylogeny of *Paralabrax* (Perciformes: Serranidae) and allied taxa inferred from partial 16S and 12S mitochondrial ribosomal DNA sequences. *Molecular Phylogenetics and Evolution* **29**, 176–184 (2003).

89.	Craig, M. & Hastings, P. A molecular phylogeny of the groupers of the subfamily Epinephelinae (Serranidae) with a revised classification of the Epinephelini. *Ichthyological Research* **54**, 1–17 (2007).

90.	Tang, C.-N. & Chen, W.-J. A 40-year taxonomic enigma: multigene phylogeny resolves the polyphyly of Plectranthias (Perciformes: Anthiadidae) and supports a revised taxonomy. *Zool J Linn Soc* **205**, zlaf148 (2025).

91.	Zhuang, X., Qu, M., Zhang, X. & Ding, S. A Comprehensive Description and Evolutionary Analysis of 22 Grouper (Perciformes, Epinephelidae) Mitochondrial Genomes with Emphasis on Two Novel Genome Organizations. *PloS one* **8**, e73561 (2013).

92.	Wang, C. *et al.* Comparative Analysis of Four Complete Mitochondrial Genomes of Epinephelidae (Perciformes). *Genes* **13**, (2022).

93.	Kim, Y., Park, J. Y., Huynh, T., Kim, K.-R. & Bang, I.-C. Complete mitochondrial genome of the hybrid grouper Hyporthodus septemfasciatus (♀)×Epinephelus moara (♂) (Perciformes, Serranidae) and results of a phylogenetic analysis. *Mitochondrial DNA Part B* **6**, 771–773 (2021).

94.	DiBattista, J. D. *et al.* Genomic and life-history discontinuity reveals a precinctive lineage for a deep-water grouper with gene flow from tropical to temperate waters on the west coast of Australia. *Ecological Genetics and Genomics* **9**, 23–33 (2018).

95.	Eastman, J. T. & Voskoboinikova, O. S. Osteology provides insight into the biology of the enigmatic Antarctic notothenioid fish Gvozdarus svetovidovi. *Polar Biol* **47**, 1137–1149 (2024).

96.	SANTINI, F. & TYLER, J. C. A phylogeny of the families of fossil and extant tetraodontiform fishes (Acanthomorpha, Tetraodontiformes), Upper Cretaceous to Recent. *Zoological Journal of the Linnean Society* **139**, 565–617 (2003).

97.	Arcila, D. & Tyler, J. C. Mass extinction in tetraodontiform fishes linked to the Palaeocene–Eocene thermal maximum. *Proceedings of the Royal Society B: Biological Sciences* **284**, 20171771 (2017).

98.	Bolet, A. *et al.* Unusual morphology in the mid-Cretaceous lizard Oculudentavis. *Current Biology* **31**, 3303-3314.e3 (2021).

99.	Brownstein, C. D. *et al.* The affinities of the Late Triassic *Cryptovaranoides* and the age of crown squamates. *Royal Society Open Science* **10**, 230968 (2023).

100.	Marramà, G., Bannikov, A. & Carnevale, G. An Eocene scorpionfish from Monte Postale (Bolca Lagerstätte, northeastern Italy). *Bollettino della Societa Paleontologica Italiana* **59**, 105–112 (2020).

101.	Cantalice, K. M., Alvarado-Ortega, J. & Alaniz-Galvan, A. *Paleoserranus lakamhae* gen. et sp. nov., a Paleocene seabass (Perciformes: Serranidae) from Palenque, Chiapas, southeastern Mexico. *Journal of South American Earth Sciences* **83**, 137–146 (2018).

102.	Patterson, C. An overview of the early fossil record of acanthomorphs. *Bulletin of Marine Science* https://www.semanticscholar.org/paper/An-overview-of-the-early-fossil-record-of-Patterson/870f1380d1dda4230507a50b5517dcd091fe096b (1993).

103.	Friedman, M. The Macroevolutionary History of Bony Fishes: A Paleontological View. *Annual Review of Ecology, Evolution, and Systematics* **53**, 353–377 (2022).

**Figure Captions.**

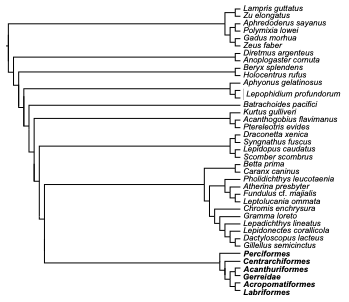

**Figure S1. Phylogeny of *Eupercaria* I: Backbone*.*** Maximum likelihood phylogeny based on

995 UCE sequences and inferred in IQ-TREE2. %BS = percent ultrafast bootstrap support.

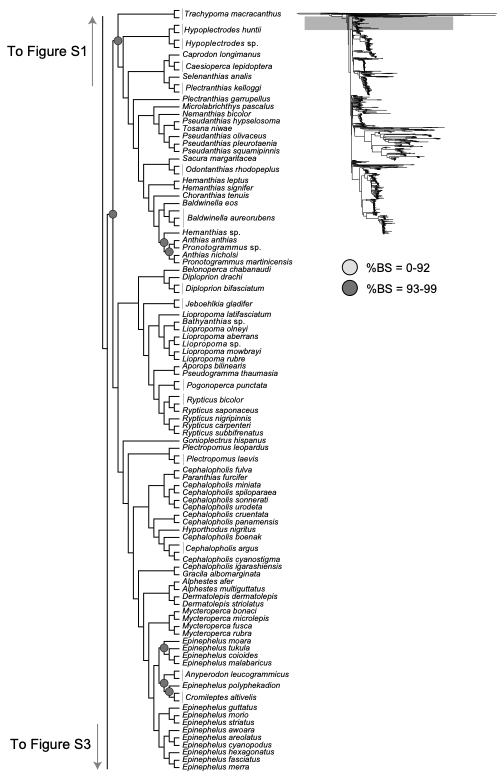

**Figure S2. Phylogeny of *Eupercaria* II: *Anthiadidae* and *Epinephelidae.*** Maximum likelihood

phylogeny based on 995 UCE sequences and inferred in IQ-TREE2. %BS = percent ultrafast

bootstrap support.

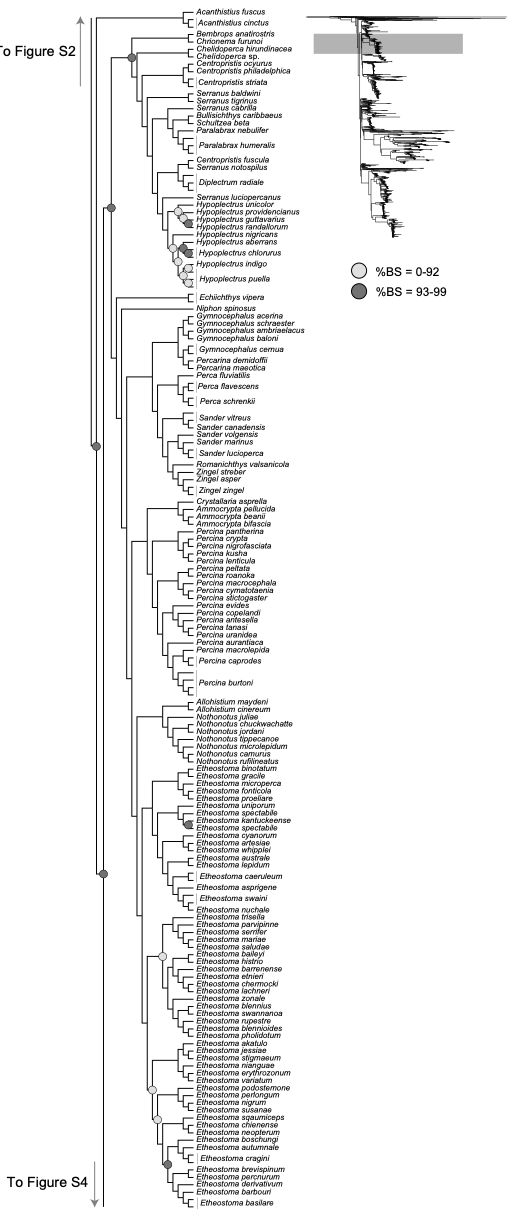

**Figure S3. Phylogeny of *Eupercaria* III: *Bembropidae* to *Percidae.*** Maximum likelihood

phylogeny based on 995 UCE sequences and inferred in IQ-TREE2. %BS = percent ultrafast

bootstrap support.

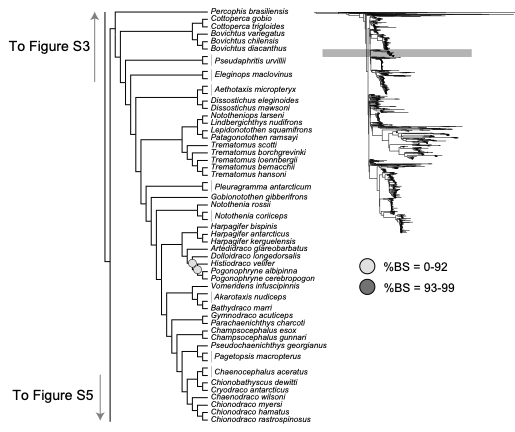

**Figure S4. Phylogeny of *Eupercaria* IV: *Notothenioidei.*** Maximum likelihood

phylogeny based on 995 UCE sequences and inferred in IQ-TREE2. %BS = percent ultrafast

bootstrap support.

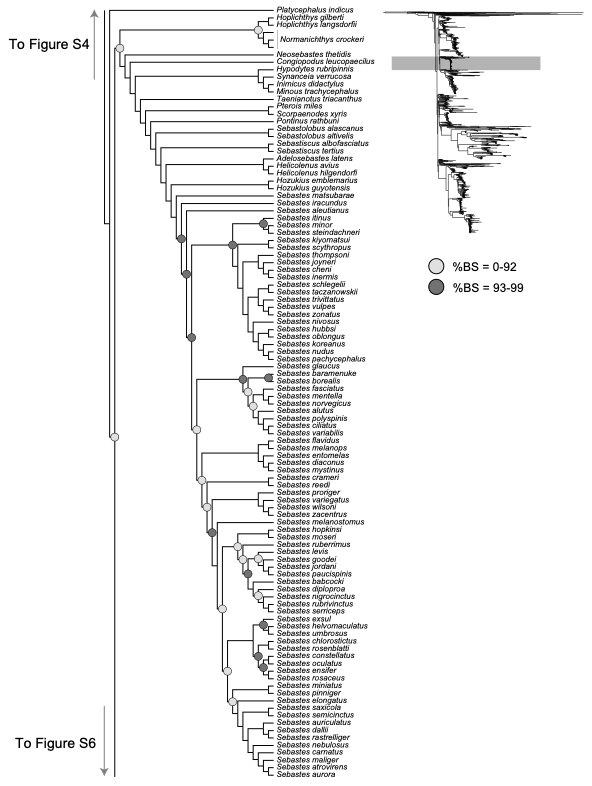

**Figure S5. Phylogeny of *Eupercaria* V: *Platycephalus* to *Scorpaenidae.*** Maximum likelihood

phylogeny based on 995 UCE sequences and inferred in IQ-TREE2. %BS = percent ultrafast

bootstrap support.
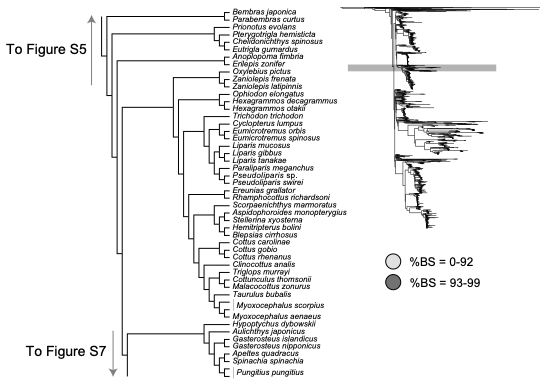

**Figure S6. Phylogeny of *Eupercaria* VI: *Anoplopomatidae* to *Cottoidea.*** Maximum likelihood

phylogeny based on 995 UCE sequences and inferred in IQ-TREE2. %BS = percent ultrafast

bootstrap support.

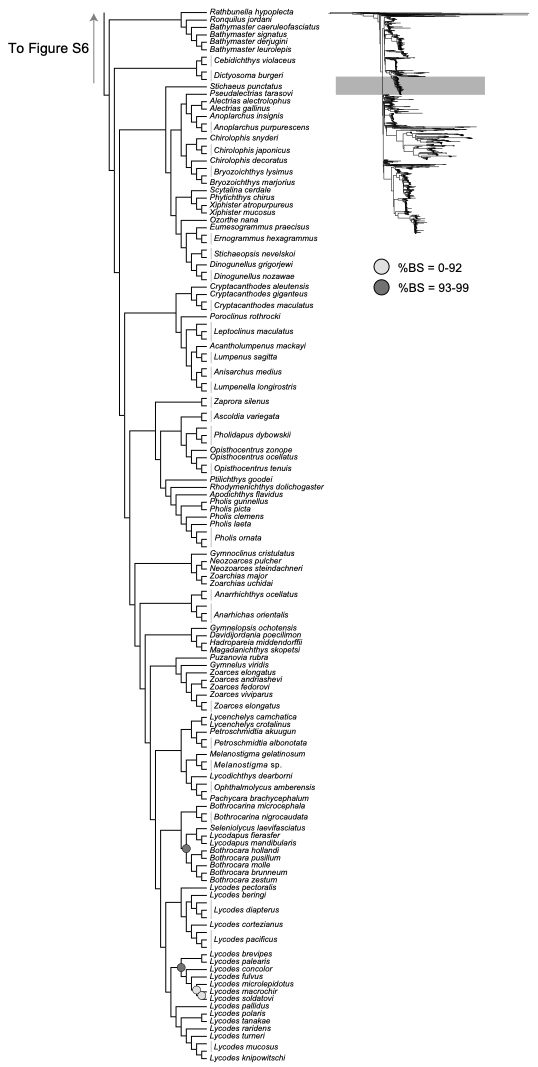

**Figure S7. Phylogeny of *Eupercaria* VII: *Zoarcoidea.*** Maximum likelihood

phylogeny based on 995 UCE sequences and inferred in IQ-TREE2. %BS = percent ultrafast

bootstrap support.

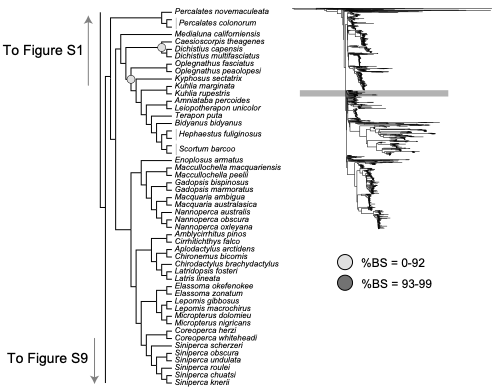

**Figure S8. Phylogeny of *Eupercaria* VIII: *Centrarchiformes.*** Maximum likelihood

phylogeny based on 995 UCE sequences and inferred in IQ-TREE2. %BS = percent ultrafast

bootstrap support.

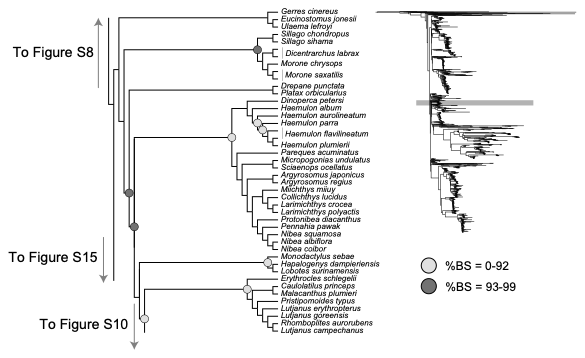

**Figure S9. Phylogeny of *Eupercaria* IX: *Gerreidae* to *Lutjanidae.*** Maximum likelihood

phylogeny based on 995 UCE sequences and inferred in IQ-TREE2. %BS = percent ultrafast

bootstrap support.

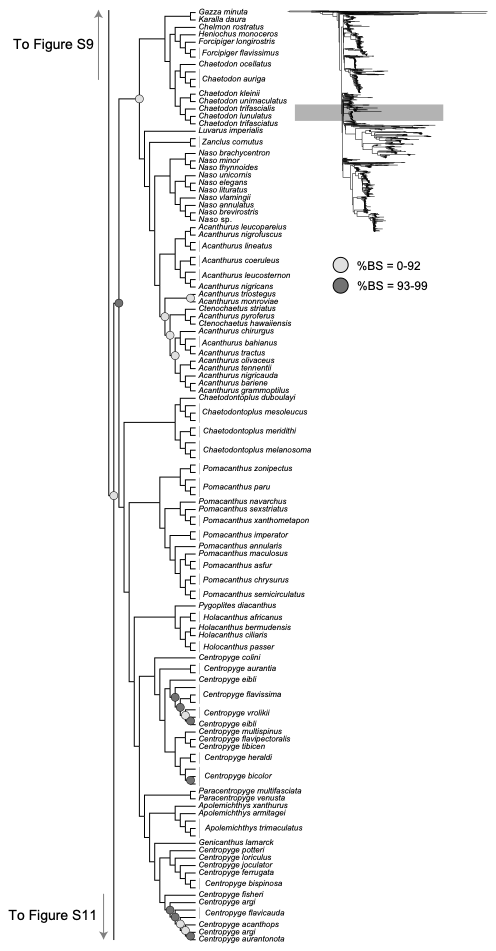

**Figure S10. Phylogeny of *Eupercaria* X: *Acanthuroidei.*** Maximum likelihood

phylogeny based on 995 UCE sequences and inferred in IQ-TREE2. %BS = percent ultrafast

bootstrap support.

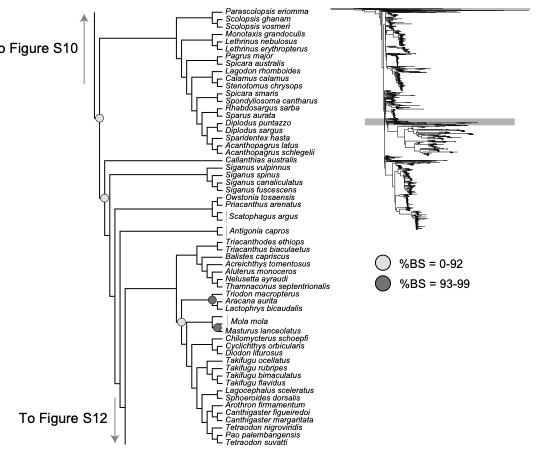

**Figure S11. Phylogeny of *Eupercaria* XI: *Nemipteridae* to *Tetraodontidae.*** Maximum

likelihood phylogeny based on 995 UCE sequences and inferred in IQ-TREE2. %BS = percent

ultrafast bootstrap support.

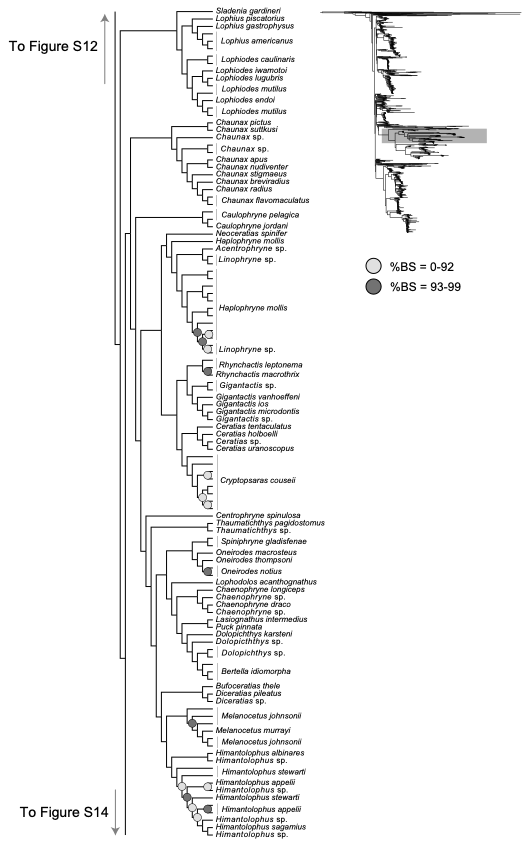

**Figure S12. Phylogeny of *Eupercaria* XII: *Lophiidae* to *Ceratioidea.*** Maximum

likelihood phylogeny based on 995 UCE sequences and inferred in IQ-TREE2. %BS = percent

ultrafast bootstrap support.

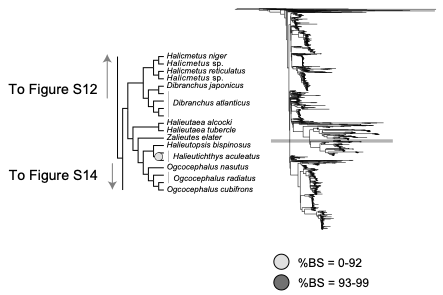

**Figure S13. Phylogeny of *Eupercaria* XIII: *Ogcocephalidae.*** Maximum

likelihood phylogeny based on 995 UCE sequences and inferred in IQ-TREE2. %BS = percent

ultrafast bootstrap support.

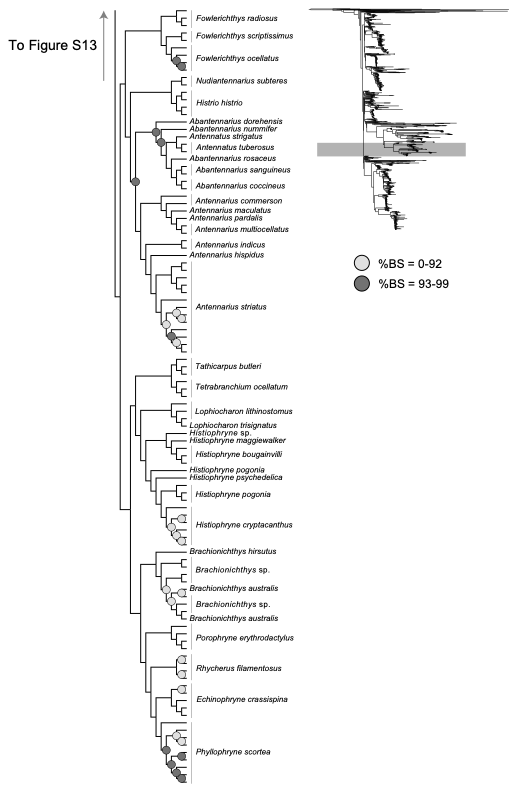

**Figure S14. Phylogeny of *Eupercaria* XIV: *Antennariidae.*** Maximum

likelihood phylogeny based on 995 UCE sequences and inferred in IQ-TREE2. %BS = percent

ultrafast bootstrap support.

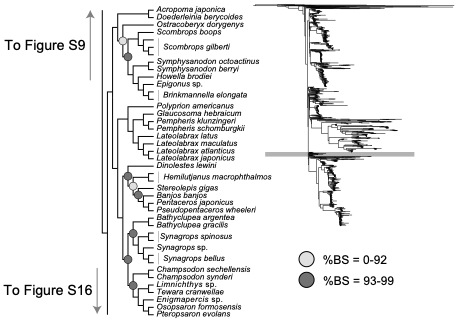

**Figure 15. Phylogeny of *Eupercaria* XV: *Acropomatiformes.*** Maximum

likelihood phylogeny based on 995 UCE sequences and inferred in IQ-TREE2. %BS = percent

ultrafast bootstrap support.

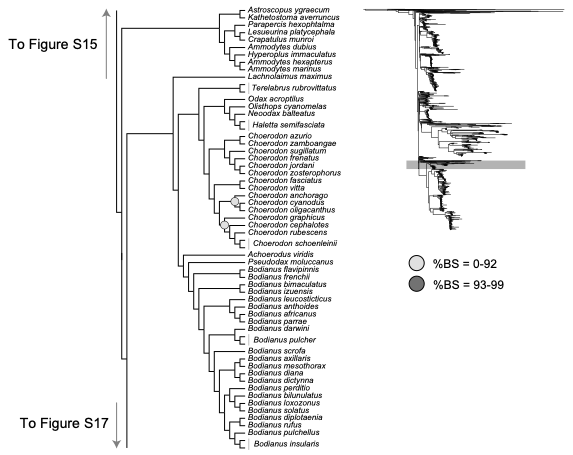

**Figure S16. Phylogeny of *Eupercaria* XVI: *Uranoscopoidei* to *Hypsigenyinae.*** Maximum

likelihood phylogeny based on 995 UCE sequences and inferred in IQ-TREE2. %BS = percent

ultrafast bootstrap support.

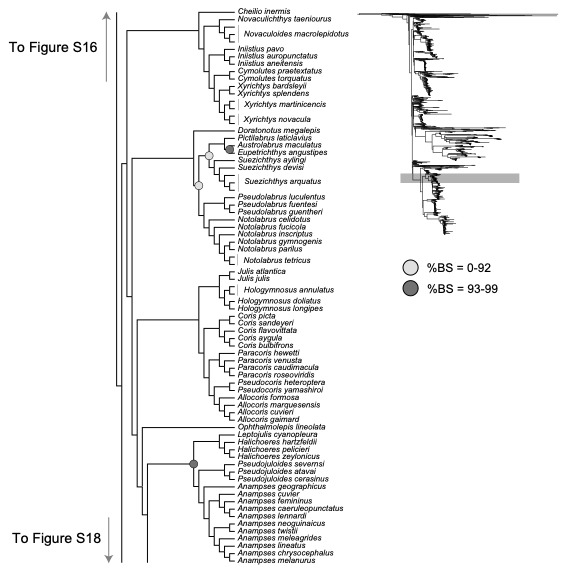

**Figure S17. Phylogeny of *Eupercaria* XVII: *Cheilio* to *Anampses.*** Maximum

likelihood phylogeny based on 995 UCE sequences and inferred in IQ-TREE2. %BS = percent

ultrafast bootstrap support.

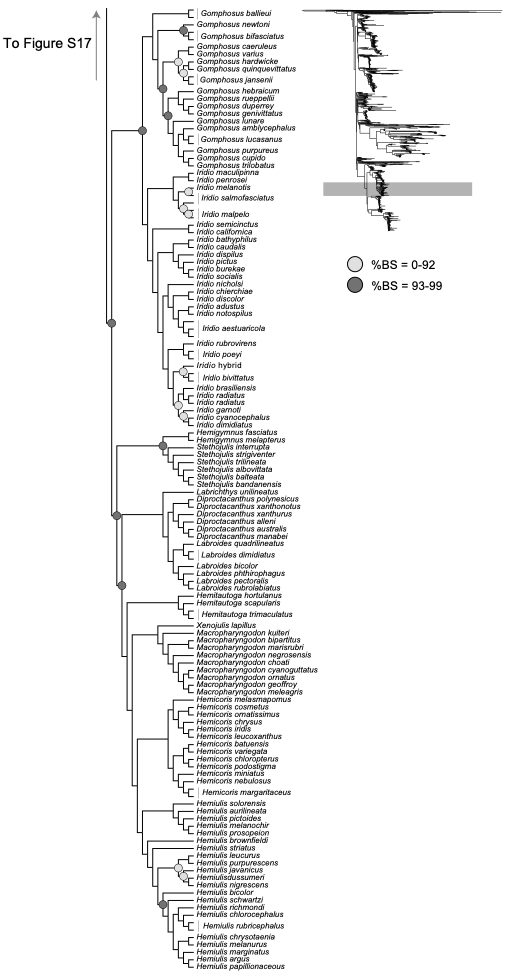

**Figure S18. Phylogeny of *Eupercaria* XVIII: *Gomphosus* to *Hemiulis.*** Maximum

likelihood phylogeny based on 995 UCE sequences and inferred in IQ-TREE2. %BS = percent

ultrafast bootstrap support.

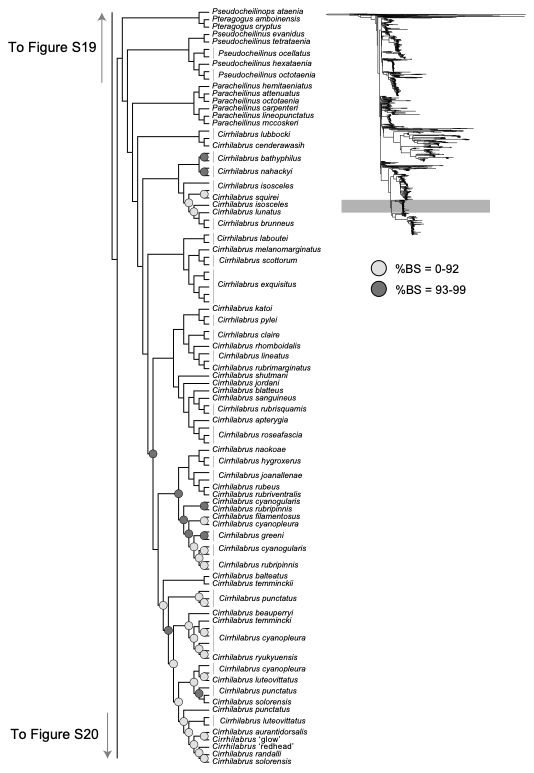

**Figure S19.** **Phylogeny of *Eupercaria* XIX: *Cirrhilabrinae.*** Maximum

likelihood phylogeny based on 995 UCE sequences and inferred in IQ-TREE2. %BS = percent

ultrafast bootstrap support.

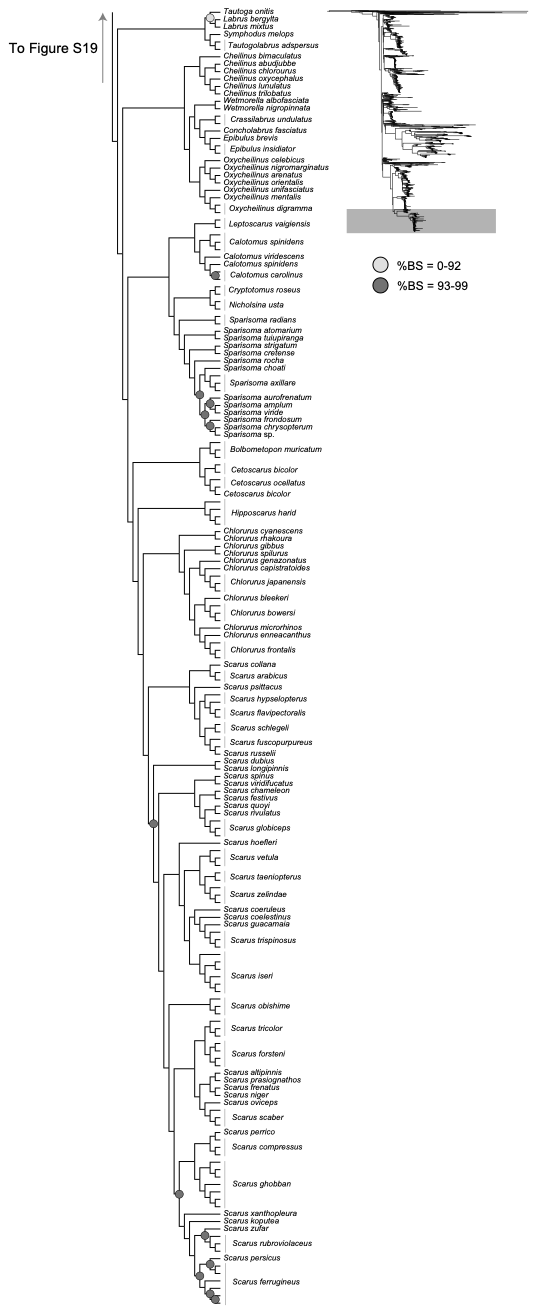

**Figure S20. Phylogeny of *Eupercaria* XX: *Labrinae* to *Scarinae.*** Maximum

likelihood phylogeny based on 995 UCE sequences and inferred in IQ-TREE2. %BS = percent

ultrafast bootstrap support.

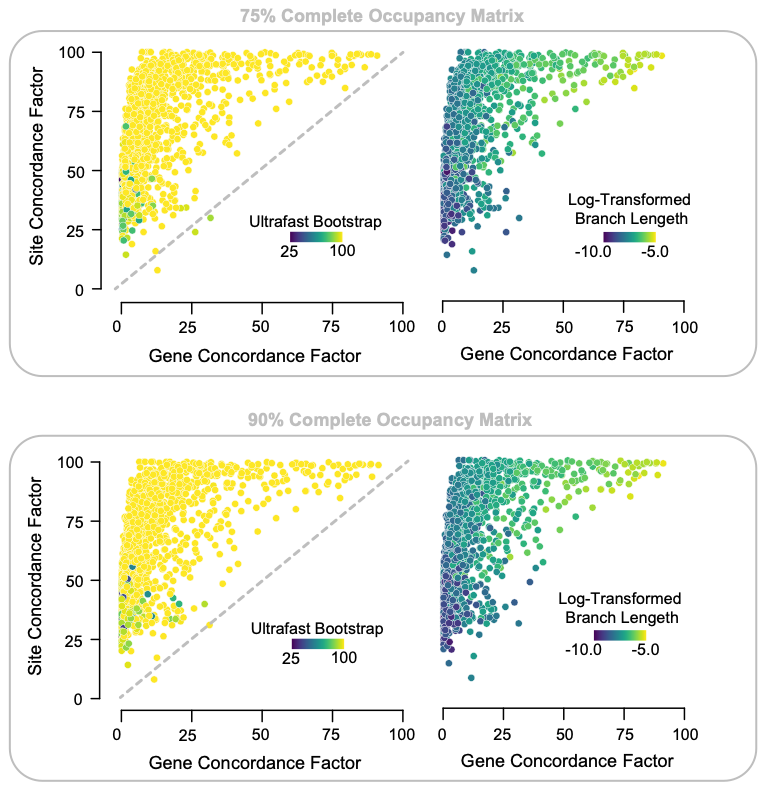

**Figure S21. Concordance Factors.** Relationships of gene and site concordance factors,

bootstrap supports, and log-transformed branch lengths for analyses of the 75% complete (top

row) and 95% complete (bottom row) matrices.

**Figure S22. Taxon-occupancy matrix for the UCE dataset.**

**Table S1. Newly Sequenced Tissues.**

| **Genus** | **Species** | **Specimen #** |
| --- | --- | --- |
| *Acanthistius* | *cinctus* | YFTC 23786 |
| *Acanthistius* | *cinctus* | YFTC 26895 |
| *Acanthistius* | *fuscus* | DRR 111234 |
| *Aethotaxis* | *micropteryx* | YFTC 15293 |
| *Aethotaxis* | *micropteryx* | YFTC 15294 |
| *Alphestes* | *afer* | BLZ 6053 |
| *Alphestes* | *multiguttatus* | DRR 113720 |
| *Anoplarchus* | *purpurescens* | YFTC 16440 |
| *Anthias* | *nicholsi* | YFTC 22491 |
| *Anyperodon* | *leucogrammicus* | DRR 113670 |
| *Anyperodon* | *leucogrammicus* | PI 0203 |
| *Aporops* | *bilinearis* | MAR Q277 |
| *Baldwinella* | *aureorubens* | YFTC 26223 |
| *Baldwinella* | *aureorubens* | YFTC 26364 |
| *Baldwinella* | *aureorubens* | YFTC 26454 |
| *Baldwinella* | *eos* | MOP 110015 |
| *Bathyanthias* | sp. | MOC 11791 |
| *Belonoperca* | *chabanaudi* | MBIO 18501 |
| *Brinkmannella* | *elongata* | YFTC 22524 |
| *Brinkmannella* | *elongata* | NV |
| *Bullisichthys* | *caribbaeus* | CUR12304 |
| *Caesioperca* | *lepidoptera* | YFTC 26890 |
| *Caesioperca* | *lepidoptera* | YFTC 26891 |
| *Caesioscorpis* | *theagenes* | YFTC16937 |
| *Caprodon* | *longimanus* | YFTC26894 |
| *Centropristis* | *fuscula* | CUR 13255 |
| *Centropristis* | *ocyurus* | YFTC 17799 |
| *Centropristis* | *ocyurus* | YFTC 1779 |
| *Centropristis* | *philadelphica* | FCC 8113 |
| *Centropristis* | *striata* | YFTC 12184 |
| *Cephalopholis* | *argus* | YFTC 12559 |
| *Cephalopholis* | *boenak* | YFTC 16959 |
| *Cephalopholis* | *cyanostigma* | YFTC 23415 |
| *Cephalopholis* | *fulva* | BLZ 8273 |
| *Cephalopholis* | *igarashiensis* | YFTC 16960 |
| *Cephalopholis* | *miniata* | YFTC 16961 |
| *Cephalopholis* | *sonnerati* | YFTC 16962 |
| *Cephalopholis* | *spiloparaea* | YFTC 16963 |
| *Cephalopholis* | *urodeta* | YFTC 16964 |
| *Chelidoperca* | sp. | PHI 422 |
| *Choranthias* | *tenuis* | CUR 11397 |
| *Cromileptes* | *altivelis* | YFTC 16967 |
| *Dermatolepis* | *dermatolepis* | DRR 110674 |
| *Dermatolepis* | *striolatus* | DRR 113687 |
| *Dinolestes* | *lewini* | YFTC 24264 |
| *Diplectrum* | *radiale* | YFTC 25735 |
| *Diplectrum* | *radiale* | YFTC 25736 |
| *Diplectrum* | *radiale* | YFTC 25737 |
| *Diploprion* | *bifasciatum* | YFTC 16941 |
| *Diploprion* | *bifasciatum* | YFTC 16942 |
| *Synagrops* | sp. | YFTC 22291 |
| *Epigonus* | sp. | YFTC 26491 |
| *Epinephelus* | *areolatus* | PI 0323 |
| *Epinephelus* | *fasciatus* | PI 0060 |
| *Epinephelus* | *guttatus* | BAH 8008 |
| *Epinephelus* | *hexagonatus* | YFTC 23377 |
| *Epinephelus* | *merra* | PI 0035 |
| *Epinephelus* | *morio* | BLZ 7860 |
| *Epinephelus* | *striatus* | BLZ 8233 |
| *Eumicrotremus* | *orbis* | YFTC 17737 |
| *Gonioplectrus* | *hispanus* | CUR 13081 |
| *Gracila* | *albomarginata* | PI 0323 |
| *Hemanthias* | *leptus* | CUR 13060 |
| *Hemanthias* | *signifer* | MOP 110012 |
| *Hemanthias* | sp. | YFTC 24876 |
| *Hemilutjanus* | *macrophthalmos* | YFTC 17239 |
| *Hemilutjanus* | *macrophthalmos* | YFTC 17240 |
| *Hypoplectrodes* | *huntii* | YFTC 26888 |
| *Hypoplectrodes* | *huntii* | YFTC 26889 |
| *Hypoplectrodes* | sp. | YFTC 26892 |
| *Hypoplectrodes* | sp. | YFTC 26893 |
| *Hypoplectrus* | *aberrans* | YFTC 16214 |
| *Hypoplectrus* | *puella* | YFTC 11472 |
| *Hyporthodus* | *nigritus* | DRR 112405 |
| *Jeboehlkia* | *gladifer* | CUR 13152 |
| *Lateolabrax* | *atlanticus* | YFTC 25777 |
| *Liopropoma* | *aberrans* | CUR 13260 |
| *Liopropoma* | *latifasciatum* | PHI 336 |
| *Liopropoma* | *mowbrayi* | CUR 13101 |
| *Liopropoma* | *olneyi* | CUR 12060 |
| *Liopropoma* | sp. | DRR 110786 |
| *Mycteroperca* | *bonaci* | BLZ 8234 |
| *Mycteroperca* | *fusca* | DRR 112802 |
| *Mycteroperca* | *microlepis* | YFTC 11502 |
| *Mycteroperca* | *rubra* | DRR 112427 |
| *Normanichthys* | *crockeri* | YFTC 25859 |
| *Normanichthys* | *crockeri* | YFTC 25860 |
| *Odontanthias* | *rhodopeplus* | PHI 325 |
| *Odontanthias* | *rhodopeplus* | PHI 326 |
| *Paralabrax* | *humeralis* | YFTC 17242 |
| *Paralabrax* | *humeralis* | YFTC 17243 |
| *Paralabrax* | *humeralis* | YFTC 17244 |
| *Paralabrax* | *nebulifer* | YFTC 11485 |
| *Paranthias* | *furcifer* | DRR 112262 |
| *Pholidapus* | *dybowskii* | YFTC 16371 |
| *Plectranthias* | *garrupellus* | CUR 12028 |
| *Plectranthias* | *kelloggi* | YFTC 16977 |
| *Plectranthias* | *kelloggi* | YFTC 16979 |
| *Plectropomus* | *laevis* | YFTC 16983 |
| *Plectropomus* | *laevis* | YFTC 16984 |
| *Pleuragramma* | *antarcticum* | YFTC 15126 |
| *Pleuragramma* | *antarcticum* | YFTC 15127 |
| *Pogonoperca* | *punctata* | MARQ 001 |
| *Pogonoperca* | *punctata* | PHI 007 |
| *Pronotogrammus* | *martinicensis* | CUR 13211 |
| *Pronotogrammus* | sp. | YFTC 24861 |
| *Pseudanthias* | *bicolor* | YFTC16986 |
| *Pseudanthias* | *hypselosoma* | PHI 334 |
| *Pseudanthias* | *olivaceus* | MBIO 18461 |
| *Pseudanthias* | *pascalus* | YFTC 12576 |
| *Pseudanthias* | *pleurotaenia* | YFTC 16988 |
| *Pseudogramma* | *thaumasia* | DRR 113838 |
| *Rypticus* | *bicolor* | DRR 110539 |
| *Rypticus* | *carpenteri* | BLZ 8013 |
| *Rypticus* | *saponaceus* | CV 11085 |
| *Sacura* | *margaritacea* | YFTC16993 |
| *Schultzea* | *beta* | CUR 12041 |
| *Selenanthias* | *analis* | YFTC 16940 |
| *Serranus* | *baldwini* | BLZ 8310 |
| *Serranus* | *luciopercanus* | CUR 11395 |
| *Serranus* | *notospilus* | CUR 13045 |
| *Synagrops* | *bellus* | YFTC 22534a |
| *Synagrops* | *bellus* | YFTC 22534b |
| *Synagrops* | *spinosus* | YFTC 22520 |
| *Synagrops* | *spinosus* | YFTC 26491 |
| *Tosana* | *niwae* | YFTC 16996 |
| *Trachypoma* | *macracanthus* | YFTC 26897 |
| *Trachypoma* | *macracanthus* | YFTC 26898 |
| *Ulaema* | *lefroyi* | YFTC 11525 |
| *Zoarces* | sp. | YFTC 16363 |

**Table S2. Loadings for PCA of body shape.** Lambda = 0.9425204. Percentages at top indicate percentage of variance explained by each component.

| Measurement | PC1  (54.21%) | | PC2 (21.81%) | PC3 (14.16%) | PC4 (6.24%) | PC5 (3.58%) |
| --- | --- | --- | --- | --- | --- | --- |
| Max body depth/SL | -0.7409413 | -0.6029209 | | -0.0701574 | -0.0420251 | 0.28426082 |
| Max width/SL | -0.773815 | 0.3065312 | | 0.40524829 | -0.3767293 | -0.0331348 |
| Head depth/SL | -0.8178033 | -0.4762879 | | -0.0554089 | 0.10242894 | -0.3013067 |
| Lower jaw length/SL | -0.5663835 | 0.4347891 | | -0.6937526 | -0.0941739 | 0.00259842 |
| Mouth width/SL | -0.7570637 | 0.4659886 | | 0.23332103 | 0.38600285 | 0.0791975 |

**Table S3. Ancestral State Reconstruction Model Fitting.** Ancestral state model fitting for the water column ecology and depth traits, when considered both as three-state and two-state characters. Best-fit models are in bold.

| Model | | log(L) | d.f. | | AIC | weight |
| --- | --- | --- | --- | --- | --- | --- |
| Ecology (three state), ER | -498.834 | | | 1 | 999.668 | 0.1924759 |
| Ecology (three state), ARD | | **-492.4** | | **6** | **996.8** | **0.8075241** |
| Ecology (two state), ER | | -296.4165 | | 1 | 594.833 | 0.6216557 |
| Ecology (two state), ARD | | **-295.9131** | | **2** | **595.8262** | **0.3783443** |
| Depth (three state), ER | | -631.416 | | 1 | 1264.832 | 2.37E-40 |
| Depth (three state), ARD | | **-535.1752** | | **6** | **1082.35** | **1.00E+00** |
| Depth (two state), ER | | -202.716 | | 1 | 407.4321 | 0.08064819 |
| Depth (two state), ARD | | **-199.2825** | | **2** | **402.5649** | **0.91935181** |

**Table S4. SSE Model Fitting.** Model fitting for SSE analyses conducted using the R package *hisse*. Numbers are AIC scores. Best-fit model in each case is bolded.

| Trait | Null Model | BiSSE | HiSSE | HiSSE, CID |
| --- | --- | --- | --- | --- |
| Swimbladder | 8492.937 | 8482.202 | **8201.836** | 8211.924 |
| Swimbladder Subset | 4018.481 | 4013.266 | **3920.328** | 3924.96 |
| Water Column Ecology | 9068.024 | 9067.074 | **8721.023** | 8768.281 |
| Water Column Ecology Subset | 4313.09 | 4303.911 | **4187.262** | 4216.639 |
| Depth | 8194.563 | 8189.764 | **7824.115** | 7905.153 |
| Depth Subset | 3842.12 | 3844.753 | **3677.795** | 3754.102 |
